# Generation and functional analysis of defective viral genomes during SARS-CoV-2 infection

**DOI:** 10.1101/2022.09.22.509123

**Authors:** Terry Zhou, Nora J. Gilliam, Sizhen Li, Simone Spaudau, Raven M. Osborn, Christopher S. Anderson, Thomas J. Mariani, Juilee Thakar, Stephen Dewhurst, David H. Mathews, Liang Huang, Yan Sun

## Abstract

Defective viral genomes (DVGs) have been identified in many RNA viruses as a major factor influencing antiviral immune response and viral pathogenesis. However, the generation and function of DVGs in SARS-CoV-2 infection are less known. In this study, we elucidated DVG generation in SARS-CoV-2 and its relationship with host antiviral immune response. We observed DVGs ubiquitously from RNA-seq datasets of *in vitro* infections and autopsy lung tissues of COVID-19 patients. Four genomic hotspots were identified for DVG recombination and RNA secondary structures were suggested to mediate DVG formation. Functionally, bulk and single cell RNA-seq analysis indicated the IFN stimulation of SARS-CoV-2 DVGs. We further applied our criteria to the NGS dataset from a published cohort study and observed significantly higher DVG amount and frequency in symptomatic patients than that in asymptomatic patients. Finally, we observed unusually high DVG frequency in one immunosuppressive patient up to 140 days after admitted to hospital due to COVID-19, first-time suggesting an association between DVGs and persistent viral infections in SARS-CoV-2. Together, our findings strongly suggest a critical role of DVGs in modulating host IFN responses and symptom development, calling for further inquiry into the mechanisms of DVG generation and how DVGs modulate host responses and infection outcome during SARS-CoV-2 infection.

**Importance:** Defective viral genomes (DVGs) are ubiquitously generated in many RNA viruses, including SARS-CoV-2. Their interference activity to full-length viruses and IFN stimulation provide them the potential for novel antiviral therapies and vaccine development. SARS-CoV-2 DVGs are generated through the recombination of two discontinuous genomic fragments by viral polymerase complex and the recombination is also one of the major mechanisms for the emergence of new coronaviruses. Focusing on the generation and function of SARS-CoV-2 DVGs, these studies identify new hotspots for non-homologous recombination and strongly suggest that the secondary structures within viral genomes mediate the recombination. Furthermore, these studies provide the first evidence for IFN stimulation activity of *de novo* DVGs during natural SARS-CoV-2 infection. These findings set up the foundation for further mechanism studies of SARS-CoV-2 recombination and provide the evidence to harness DVGs’ immunostimulatory potential in the development of vaccine and antivirals for SARS-CoV-2.

## Introduction

Respiratory tract infection of severe acute respiratory syndrome coronavirus 2 (SARS-CoV-2) results in varying immunopathology underlying coronavirus disease 2019 (COVID-19). Its symptoms vary from asymptomatic infection to milder/moderate disease and further critical illness, including respiratory failure and death. Immune responses in COVID-19 patients of various disease severities have been studied (Lega, Naviglio et al. 2020, Chiale, Greene et al. 2022, Dadras, Afsahi et al. 2022). In general, broad induction of IFN responses and antiviral genes are associated with milder/moderate COVID-19, whereas severe COVID-19 is often characterized by a blunt early IFN responses and elevated proinflammatory cytokine expression in nasopharyngeal mucosa (Kwon, Kim et al. 2020, Liu, Li et al. 2020, Gozman, Perry et al. 2021, Janssen, Grondman et al. 2021, Vanderbeke, Van Mol et al. 2021). Investigation of how IFN responses are induced by SARS-CoV-2 infection, especially early IFN stimulation in some patients, requires further study.

During SARS-CoV-2 infection, in addition to full-length viral genomes and single nucleotide mutations, three major types of viral RNAs are generated from non-homologous recombination that are critical for viral pathogenesis, including subgenomic mRNAs (sgmRNAs), structural variants (SVs), and defective viral genomes (DVGs). The viral replication-transcription complex performs recombination at specific transcription regulatory sequences (TRSs) to generate a set of sgmRNAs, which subsequently translate into viral structural proteins (van Hemert, van den Worm et al. 2008, Dufour, Mateos-Gomez et al. 2011, Sola, Almazán et al. 2015, Brant, Tian et al. 2021). SVs comprise small insertion/deletions that allow the variant genome to independently replicate and transmit. Numerous SVs have been described including small deletions in viral spike protein that alter the fitness and virulence of SARS-CoV-2 isolates (Davidson, Williamson et al. 2020, Li, Wu et al. 2020, Majumdar and Niyogi 2021, Wang, Lau et al. 2021). Different from sgmRNAs and SVs, SARS-CoV-2 DVGs contain large internal deletions and have recombination positions distinct from TRSs while retaining 5’ and 3’ genomic untranslated regions (UTRs) (Gribble, Stevens et al. 2021).

This type of DVGs, also known as defective viral or interfering RNAs (D-RNAs), is widely generated during replication of most positive sense RNA viruses (Huang 1973, Marcus and Sekellick 1977) and influenza (Nayak, Chambers et al. 1985), and their replication relies on viral machinery provided by co-infected homologous full-length viruses (Huang and Baltimore 1970, Brian and Spaan 1997, Wu and Brian 2010). When accumulated to a high level, DVGs can interfere with full-length viral genome production by stealing essential viral elements from full-length viruses (Roux, Simon et al. 1991, Vignuzzi and López 2019). This interference activity has been reported for influenza viruses (De and Nayak 1980) and multiple non-SARS-CoV-2 coronaviruses (CoVs), such as SARS-CoV (Raman and Brian 2005), mouse hepatitis virus (MHV) (Makino, Fujioka et al. 1985), bovine CoV (Hofmann, Sethna et al. 1990), avian infectious bronchitis virus (IBV) (Pénzes, Wroe et al. 1996), transmissible gastroenteritis virus (Méndez, Smerdou et al. 1996), and middle east respiratory syndrome CoV (MERS-CoV) (Gribble, Stevens et al. 2021). In addition to interference activity, DVGs from influenza A virus have strong IFN stimulation (Kupke, Riedel et al. 2019) and are reported to promote viral persistence *in vitro* (De and Nayak 1980, Moscona 1991, Frensing, Heldt et al. 2013). More importantly, DVGs are largely observed in nasal samples from patients positive for influenza and their abundance is negatively correlated with patients’ disease severity, indicating the critical roles of DVGs in host responses and clinical outcome (Vasilijevic, Zamarreño et al. 2017). The current approach to identify DVGs from SARS-CoV-2 infection is through short-read and long-read next generation deep sequencing (NGS). Several algorithms, such as DI-tector (Beauclair, Mura et al. 2018), VODKA (Viral Opensource DVG Key Algorithm) (Sun, Kim et al. 2019), and ViReMa, (Viral-Recombination-Mapper) (Routh and Johnson 2014), and metasearch tool DVGfinder (Olmo-Uceda, Muñoz-Sánchez et al. 2022) are developed to specifically detect the reads containing the recombination sites of DVGs. Using these approaches, DVGs are documented in SARS-CoV-2 infected Vero E6 cells (Chaturvedi, Vasen et al. 2021, Rand, Kupke et al. 2021) and in nasal samples of COVID-19 patients (Xiao, Lidsky et al. 2021). Long-read NGS, such as full length iso-seq and nanopore direct RNA-seq, further confirmed that substantial TRS-independent deletions identified from short-read NGS are from SARS-CoV-2 genomes and maintain two genomic ends (Gribble, Stevens et al. 2021, Wong, Ngan et al. 2021). Additionally, identical deletions are found in various transcripts encoding distinct sgmRNAs (Wong, Ngan et al. 2021), strongly suggesting that even deletions in sgmRNAs are likely to be originated from viral genomes, since deletions existing in the viral genome can be used as the template to generate a set of sgmRNAs with the same deletions during transcription.

Despite DVGs playing such an important role in viral pathogenesis, their function in SARS-CoV-2 biology is less known. Recent reports show that synthetic SARS-CoV-2 DVGs (named therapeutic interfering particles, TIPs) exhibit substantial reduction on viral load across different viral variants when delivered in hamsters (Chaturvedi, Vasen et al. 2021) and mice (Xiao, Lidsky et al. 2021) pre- or shortly after infection, demonstrating the potential of SARS-CoV-2 DVGs as a new class of antiviral intervention by interfering genomic replication. No reports have been identified for the role of DVGs in IFN responses and viral persistence for SARS-CoV-2 infection so far. Interestingly, a COVID-19 cohort study (Wong, Ngan et al. 2021) indicates that the abundance of TRS-independent deletions (>20nts) is significantly more in symptomatic patients than that in asymptomatic patients, suggesting a potential role of DVGs in modulating host responses and symptom development in COVID-19 patients.

As our interest lies with the generation of DVGs, in relation to viral pathogenesis rather than sgmRNAs or smaller deletions in SVs, we used a pipeline based on ViReMa combined with sequence filtering via RStudio to specifically identify TRS-independent DVGs with deletion lengths larger than 100nts. We identified DVGs with varying degrees of junction frequency, termed J_freq_, from multiple NGS datasets that are either publicly available or from our own infections. Interestingly, we found DVG junctions consistently clustered in several genomic hotspots among different NGS datasets and secondary structures within viral genome are likely to guide the recombination. Functionally, we found that with similar infection level, samples with more DVG reads had enhanced type I/III IFN responses than samples with less or no DVGs, indicating the potential IFN stimulation of SARS-CoV-2 DVGs. In support, analysis of single cell RNA-Seq from infected primary human lung epithelial cells showed an earlier primary IFN expression (IFNB and IFNL1) in DVG+ cells than in DVG− cells. Finally, we applied our DVG analysis to several published NGS datasets from nasal samples of COVID-19 patients. We found persistent DVG reads with unusually high frequency in one immunosuppressive patient and higher DVG abundance in symptomatic patients than asymptomatic patients. Taken together, our analyses demonstrate critical roles of DVGs in modulating host IFN responses, viral persistence, and clinic outcome for SARS-CoV-2 infection.

## Results

### DVGs are ubiquitously produced during SARS-CoV-2 infection both *in vitro* and in patients

To examine whether DVGs can be detected universally during SARS-CoV-2 infections, we used the ViReMa pipeline (Virus Recombination Mapper) combining with R filtering (Fig. S6) to specifically map the DVG recombinant sites (Fig. 1A) in multiple next generation sequencing (NGS) datasets. As reported previously, ViReMa can agnostically detect RNA recombination events and reported these junction positions in BED files. Reported junction positions include sgmRNAs, of which their junctions contain leader transcriptional-regulatory signal (TRS-L, within the first 85 nts of leader), and other recombinant RNAs with their jumping positions that are far away from TRS-L. We defined our targeted DVGs as TRS-L independent RNA species bearing deletions larger than 100 nts (Fig. 1A). Use these criteria, we first examined DVGs in 4 publicly available *in vitro* infected NGS datasets with various cell types, MOIs, viral stocks, and sample origins (Table S1). We found that DVGs can be detected in all examined datasets ranging from several counts to several thousand counts (Fig. 1B). As the infection level varied significantly among different datasets, we normalized DVG levels by junction frequency (J_freq_), a ratio of DVG counts over virus counts. DVG counts were the total number of DVG reads obtained from ViReMa and meeting the above criteria, whereas virus counts were the total amount of reads fully aligned to the reference viral genome. We observed two ranges of J_freq_, <0.1% and 0.1%-1%. A549-ACE2 infected cells have the highest J_freq_, whereas infections in NHBE varied. In addition, either total RNA or polyA enriched RNA were used for NGS for Calu3-total RNA and Calu3-polyA, respectively. Both samples had very similar J_freq_, suggesting J_freq_ is robust to different library preparation methods. Interestingly, we detected DVGs, although with low J_freq_, in the supernatants collected from infected Vero E6 cells, suggesting that certain DVG species generated within infected cells, potentially the DVGs containing packaging signals, were able to be packaged into virions and released out into supernatants.

**Figure 1.**
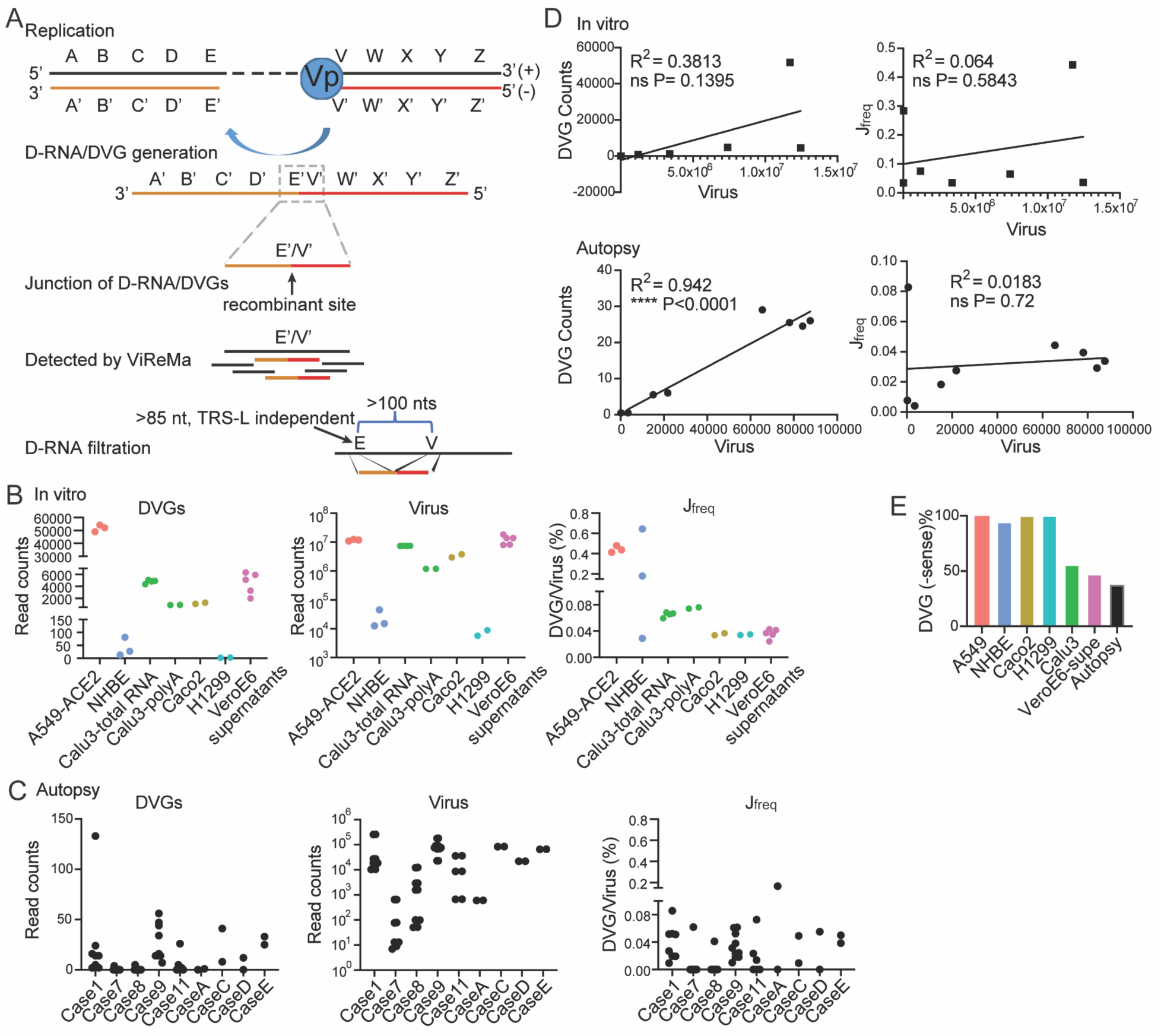
DVGs were ubiquitously generated in SARS-CoV-2 in vitro infections and autopsy tissues of COVID-19 patients. **(A)** Schematic representation of DVG generation from positive sense viral genome and the general principle of ViReMa identification of deletion DVGs. The V’ site represents the break point and the E’ site represents the rejoin point of the viral polymerase in the formation of DVGs. The gray dashed box marks the recombinant site that distinguishes DVGs from full length viral genomes, which are identified by ViReMa, and further filtered using two criteria shown in the graph. **(B)** The DVG read counts, viral read counts, and J_freq_ percentages were graphed for each of the in vitro samples including the infected cells and supernatants. **(C)** The DVG read counts, viral read counts, and J_freq_ percentages were graphed for autopsy lung tissues of 9 DVG + COVID-19 patients. Each case represented one patients and different dots represented RNA-seq from the different location of the same lung tissues. **(D)** The correlations between DVG read counts and viral read counts were plotted for both the in vitro and autopsy samples. ****p<0.0001 by Pearson’s correlation. **(E)** The percentage of −sense DVG among total DVGs in in vitro and autopsy samples were shown.

We then examined DVGs in autopsy tissues from patients that unfortunately died from COVID-19 complications (GSE150316). We analyzed lung, heart, jejunum, liver, and kidney from 19 cases and DVGs were observed in only lung tissues in 9 cases (Fig. 1C). Their DVG counts were close to the level observed in infections in NHBE cells but much less compared to infections in cell lines, such as A549-ACE2, Vero E6, Calu3, and Caco2. J_freq_ from autopsy tissues were mostly less than 0.1%, comparable with the lower range of J_freq_ observed from *in vitro* infections. Next, we sought to examine the relationship between DVG production and viral replication. Interestingly, we observed strong positive correlation between DVG counts and virus counts for autopsy tissues, but not for *in vitro* infections (Fig. 1D). In addition, J_freq_ was not significantly correlated with virus replication level. It is noted that both negative sense (−sense) and positive sense (+sense) DVGs were detected in all NGS datasets. The percentage of −sense DVGs dominated in most *in vitro* infected NGS using total RNA to prepare the library (Fig. 1E). Together with the previous reports in nasal specimens of COVID-19 patients (Xiao, Lidsky et al. 2021) and our own analysis, we concluded that DVGs are ubiquitously generated during SARS-CoV-2 infection *in vitro* and in patients.

### Recombination sites of SARS-CoV-2 DVGs were clustered in certain genomic hotspots

To characterize positions of DVGs’ recombination sites, we graphed the actual junction positions of all identified DVGs from *in vitro* infections from different cells and DVG+ autopsy tissues. As both +sense and −sense DVGs were identified, we examined their distributions separately and first analyzed the junction positions of −sense DVGs. Interestingly, we found that their generation were clustered in three conserved genomic hotspots, indicated as junction areas A, B, C (green boxes in Fig. 2A and 2B). Among them, area B was observed in all infections and area A was largely observed in infected cells but absent in the supernatants from infected Vero E6 cells. As DVGs formed in junction area A contained the largest deletion compared to B and C, it is possible that DVGs within area A lack the package signal and thus were less efficiently released into supernatants. To further identify the genomic hotspots for DVG break and rejoin points, we graphed their locations separately based on the junction frequency per DVG position. We identified one major hotspot for break point, corresponding to genomic positions 28200-29750 (highlighted in grey dashed box in Fig. 2C, details in Fig. 2D). Additionally, three major rejoin hotspots were identified including 700-2500 (red box), 6500-8200 (yellow box), and 27000-29400 (green box). When comparing the distribution between −sense and +sense DVGs, we observed that rejoin points, V, of +sense DVGs shared the same hotspots with break point, V’, of −sense DVGs (Fig. S1A-D vs Fig. 2A). This suggests that the junction positions of −sense and +sense DVGs are correlated, likely resulting from their self-replication (Fig. S1E). Finally, we ought to examine whether common DVGs can be detected from different infection or different autopsy tissues. We only identified common DVGs from different *in vitro* infections within the same RNA-seq dataset (likely used the same viral stock for infections, Table S2). We did not find any common DVGs from different autopsy tissues. Taken together, our analysis from multiple NGS datasets indicated that SARS-CoV-2 DVGs are not generated randomly, rather they are formed at specific genomic regions.

**Figure 2.**
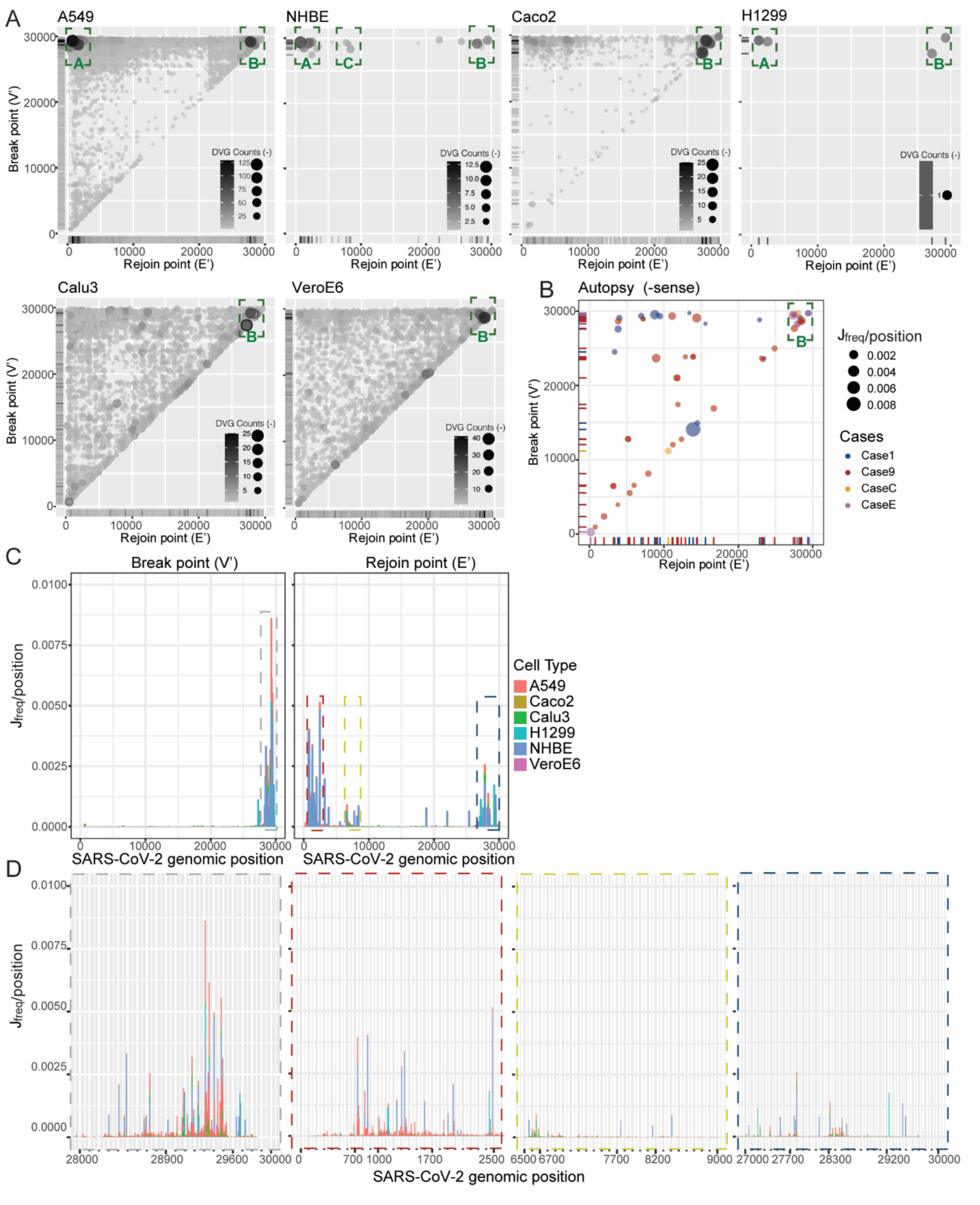
Four genomic hotspots were identified for DVG formation. Break point (V’) and rejoin point (E’) distributions for −sense DVGs from in vitro samples **(A)** and autopsy samples **(B)**. Circle size and color intensity indicated the DVG counts. The green dashed boxes represented hotspots clustered with DVG junctions. **(C)** Break point (V’) and rejoin point (E’) distributions by J_freq_ per position for all in vitro samples. The dashed boxes indicated hotspots with high concentrations of break or rejoin points. The width of each bar represented 300 nts. **(D)** Detailed positions of 4 identified hotspots clustered with DVG break and rejoin points. The color of the dashed outline around each graph indicated the corresponding hotspot with the same color in (C). The width of each bar represented 10 nts.

### The RNA structure distance between SARS-CoV-2 DVG junction positions is shorter than any two random SARS-CoV-2 genomic positions

Ziv et al. developed COMRADES (Ziv, Gabryelska et al. 2018), which can probe RNA base pairing inside cells, and applied it to detect short- and long-range interactions along the full-length SARS-CoV-2 genome (Ziv, Price et al. 2020). Interestingly, the positions of SARS-CoV-2 DVG junctions correlated well with the pairings found by COMRADES (red arches in Fig. 3A), which suggests a role of RNA secondary structures in the formation of DVGs. The paired bases bring distant nucleotides in the primary sequence close and make it possible for the breaking and rejoining actions to occur around those close pairs. To further study the relationship between DVG junctions and the identified secondary structure within the SARS-CoV-2 genome, we calculated the structural distance between DVG junction positions, which is the shortest distance between two nucleotides by traversing the backbone and base pairs (red solid path in Fig. 3B) (Clote, Ponty et al. 2012). We further extended this definition to allow competing base pairs from alternative secondary structures since many RNAs are known to populate multiple conformations in equilibrium and Ziv et al.’s data included alternative conformations of SARS-CoV-2.

**Figure 3.**
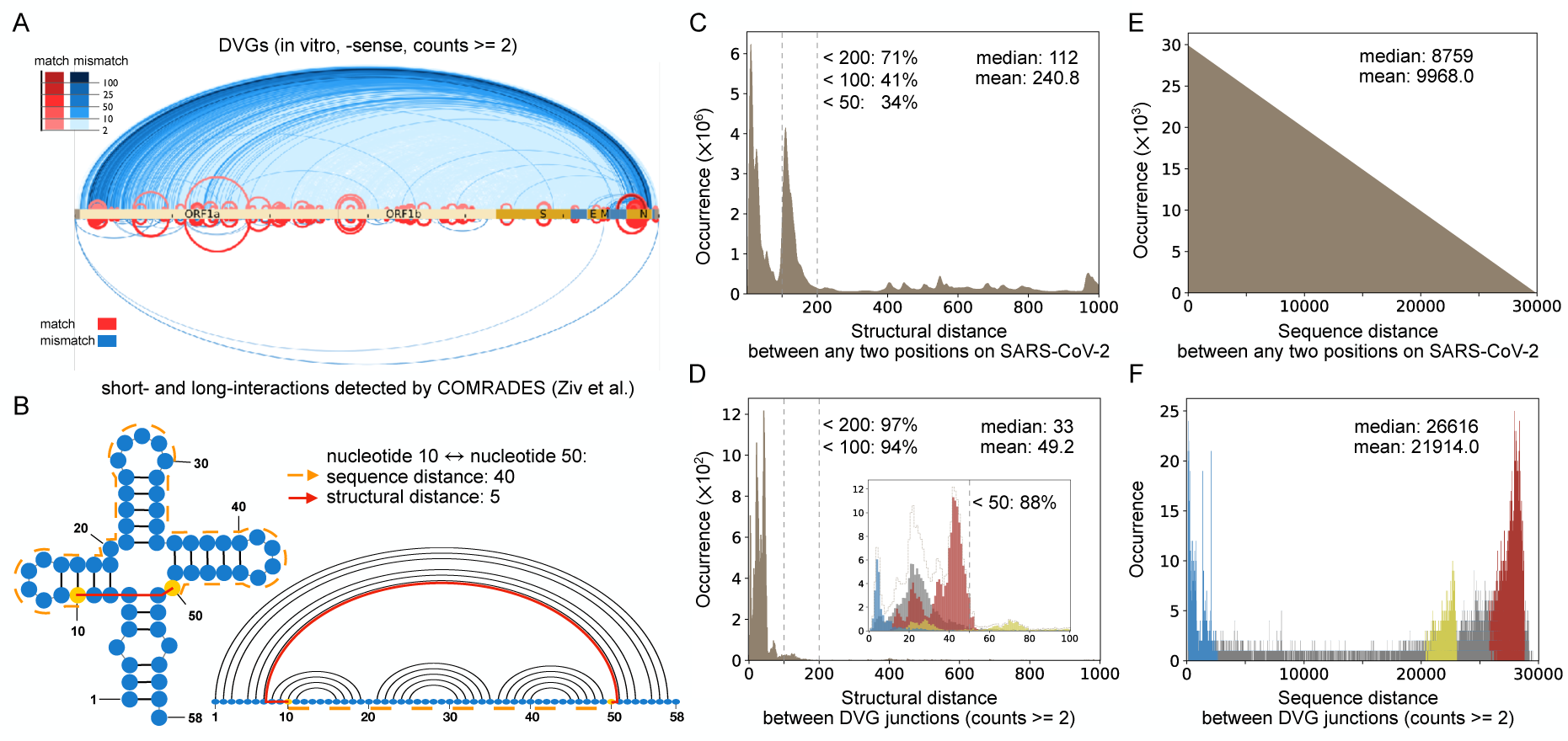
The correlation between DVGs and secondary structures. **(A)** Comparison between DVG junction positions (top, in vitro, −sense DVGs) and chimeric reads from COMRADES (bottom) along full-length SARS-CoV-2 genome (Ziv, Price et al. 2020). The red arches represented DVG positions that match COMRADES crosslinks and the blue arches represented positions that do not match crosslinks. **(B)** Example to compare sequence distance and structural distance. The structural distance between nucleotides 10 to 50 is only 5 (red solid path that includes a connection across a base pair), while the sequence distance is 40 (orange dashed path). **(C–D)** The distribution of all structural distances between any two positions in SARS-CoV-2 (**C**), and between SARS-CoV-2 DVG junction positions **(D)**. The percent of distances less than 50, 100 and 200 were indicated, respectively. **(E–F)** As a negative control, the distribution of all sequence distances between any two positions in SARS-CoV-2 **(E)**, and between SARS-CoV-2 DVG junction positions **(F)**. The mean and median distances of all distributions were annotated in **C–F**. In **(D)** and **(F)**, the blue, yellow, and red bars corresponded to three hotspots annotated in Fig. 2C, respectively, while the grey bars were out of the range of these detected hotspots. The inset in **(D)** distinguished structural distance’s distributions of three hotspots and the rest up to a structural distance of 100. The dashed contour in the inset represented the sum of all distributions for the same structural distance, and it was with the same shape as the major figure in **(D)**. In both (**C)** and **(E)**, the total occurrence of all distances equals the number of any two positions along SARS-CoV-2, and in **(D)** and **(F)**, the total occurrence of all distances is the same as the number of DVG data points (with counts 2 or above).

We first analyzed the distribution of all structural distances between any two nucleotides in SARS-CoV-2 (counts >= 2), where 41% of the distances were under 100 (Fig. 3C) with a long tail up to 1200. The median distance of the distribution was 112. However, for the structural distances only between SARS-CoV-2 DVG junction positions, the peak of the distribution shifted to the left with a smaller median value 33, and the vast majority (94%) of distances were less than 100 (Fig. 3D). Therefore, the structural distances between DVG junction positions were substantially shorter than the distances between any two random positions, which indicated a strong correlation between secondary structures and DVGs formation. Moreover, we observed that the larger the cutoff value for DVG counts, the greater the proportion of distances under 100 and the smaller the mean distance (Fig. S2). As a negative control, we also evaluated the sequence distance, which is the distance between nucleotides only based on their positions along the primary sequence; in fact, it is a special case of structural distance without any secondary structure. We analyzed the sequence distance between any two nucleotides in SARS-CoV-2 and between SARS-CoV-2 DVG junction positions (Fig. 3E and 3F), respectively. The distribution of sequence distances between any two nucleotides on SARS-CoV-2 was a triangular distribution. Most of the distances between DVG junctions were clustered similarly as the hotspots previously observed (Fig. 2C vs Fig. 3F), which is completely different from the distribution of structural distances of DVG junctions that has its peak on the left (Fig. 3C and 3D).

### SARS-CoV-2 DVGs specifically enhanced type I/III IFN responses

To understand the dynamics of SARS-CoV-2 DVGs during infection and how that affects host responses and viral replication, we infected PHLE cells from donors of different age groups with SARS-CoV-2 Hong Kong strain (SARS-CoV-2/human/HKG/VM20001061/2020) at MOI of 5. Mock and infected cells were harvested at different time points post infection (hpi) followed by bulk RNA-seq-ViReMa analysis. We observed DVGs as early as 48 hpi in cells from infants and younger adults, whereas in the elderly sample, we did not detect DVGs until 72 hpi (Fig. 4A), suggesting that DVG generation may be delayed in the elderly who are more likely to display severe symptoms when infected. We observed the same genomic hotspots for DVG junction regardless of their age groups and time points (Fig. S3A-S3D). Strikingly, those hotspots were consistent with the ones identified from different cell lines (Fig. 2), autopsy lung tissues (Fig. 2), and the following single cell RNA-seq analysis (Fig. S3E). Again, we observed that V (rejoin point of +sense DVGs) and V’ (break point of −sense DVGs) shared the same hotspots and E (break point of +sense DVGs) and E’ (rejoin point of −sense DVGs) shared the same hotspots (Fig. S3A vs S3B), indicating that our identified recombination sites were likely from DVGs capable of replication.

**Figure 4.**
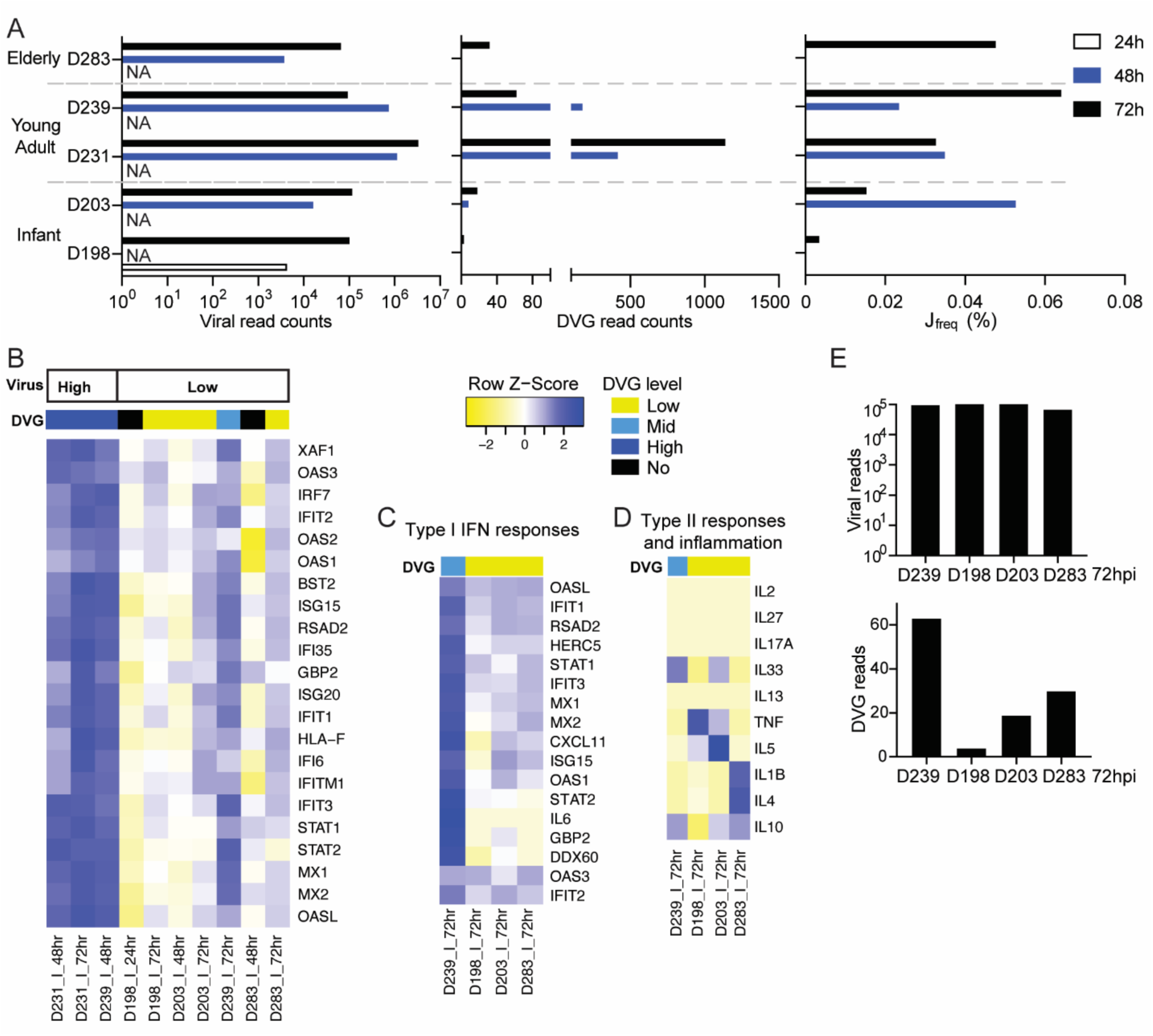
DVGs influence type I/III interferon responses in infected PHLE cells. PHLE cells of donors from different age groups were infected with SARS-CoV-2 at MOI of 5. Samples were harvested at designated time points post infection. **(A)** Viral read counts, DVG read counts, and J_freq_ were graphed for all samples, grouped by donor’ age group and time points. NA indicated that the samples were not available for RNA-seq and thus no data were collected. **(B)** Differential expression levels of genes related to type I interferon responses were graphed as heat map for all infected samples. Samples were grouped by viral infection level. DVG levels of each sample were indicated by different color codes on top of the heatmap. Four infected samples at 72 hpi with similar level of viral counts were selected to compare their IFN responses **(C)** and other gene expression unrelated to type I/III IFN responses **(D)**. **(E)** The viral and DVG read counts for the selected 4 infected samples (D198, D203, D239, and D283) were graphed.

In order to examine the role of DVGs in host responses, we grouped our infected samples based on their DVG counts and viral counts. Three samples (D231_I_48hr, D231_I_72hr, and D239_I_48hr) were significantly higher in both viral counts and DVG counts and thus categorized as High group (marked dark blue in Fig. 4B and S4A). When compared this group with the rest infected samples, one cluster of genes (pink cluster) were identified as upregulated in the High group. Gene Ontology (GO) enrichment analysis of this cluster was highly enriched in genes involved in type I IFN antiviral responses (Fig. S4B). A heatmap focusing on type I/III IFN related genes confirmed that samples in High group had enhanced gene expression compared to the rest of samples (Fig. 4B). In order to test if the IFN stimulation is specific to DVGs, we selected 4 samples at 72 hpi with similar levels of viral replication but different level of DVGs (Fig. 4E) to compare their type I/III IFN responses. We observed that the sample with more DVGs exhibited enhanced antiviral responses than samples with less DVGs (Fig. 4C), but this enhancement was not observed for genes in other pathways such as type II responses and inflammation (Fig. 4D). Although we cannot perform proper statistical analysis due to limited sample size, these data, for the first time, suggest that SARS-CoV-2 DVGs enhance IFN production as observed previously in other RNA viruses (Kupke, Riedel et al. 2019).

### Primary IFNs were expressed earlier in DVG+ cells with moderate infection

To understand DVG generation and their host responses at single cell level, we obtained one single cell RNA-seq dataset using adult NHBE cells with infection at MOI of 0.01 (GSE166766). Consistent with the previous observations, viral counts, DVG counts and J_freq_ at 2 dpi were all significantly increased compared to that at 1 dpi, but not significantly different from 3 dpi (Fig. 5A). Major cell types enriched with DVGs were ciliated cells, basal cells, and SLC16A7+ (red in Fig. 5B, grouping of cell types were based on the markers used in original publication). Among these three cell types, ciliated cells had the most DVG+ cells, whereas SLC16A7+ cells had the highest percentage of DVG+ (Fig. 5C). All DVG+ cells contained at least 1 viral count (virus positive cells) and total viral counts were significantly higher in DVG+ cells than DVG− cells at all three time points (Fig. 5D). Only about 1% of virus positive cells at 1 dpi (n=60) were DVG+. Therefore, we focused on the DVG+ population at 2 dpi (n=348) and 3 dpi (n=725) to analyze their host responses. Differential expression tests were then performed using three different methods in Seurat (MAST, Wilcox, and DEseq2) between DVG+ and DVG− groups within virus positive cells. Significantly more genes were identified as downregulation in DVG+ cells than genes that were upregulated at both time points (adj_pvalue < 0.01 and logFC > 0.25) and similar enriched pathways were observed from GO analysis. Specifically, the ribosomal cytoplasmic translation (host protein synthesis) was largely inhibited in DVG+ cells, possibly due to their higher level of viral replication (more expression of NSP1) than DVG− cells (2 dpi: upper panel in Fig. 6A; 3 dpi: Fig. S5A). Despite of this, pathways such as transcription from RNA polymerase II promoter, TNF and NF-kB, and apoptosis were significantly enriched in the upregulated genes. Importantly, defense to virus and chemokines were also observed in the upregulation list, consistent with the results from bulk RNA-seq (2 dpi: bottom panel in Fig. 6A, 3 dpi: Fig. S5B). Next, we specifically examined the expression level of representative genes related to type I/III IFN pathways between DVG− and DVG+ viral positive cells, including two primary IFNs (IFNB1 and IFNL1), ISGs and chemokines selected from the differentially expressed gene list. To better control viral loads, we further categorized virus positive cells (cells with virus count ≥1) based on their viral counts as three groups: low (viral counts ≤10), moderate (10< viral counts <20000 for 1 dpi and 2 dpi; 10< viral counts <10000 for 3 dpi), and high (viral counts ≥20000 at 1 dpi and 2 dpi; viral counts ≥10000 at 3 dpi). DVGs were identified majorly in moderate (∼12%) and high groups (>84%), and extremely small percentage (<0.2%) of low infected cells generated DVGs. Two primary IFNs were predominantly expressed only in moderate viral group regardless of DVG presence. However, DVG+ cells expressed two primary IFNs 1 day earlier than DVG− cells (2 dpi vs 3 dpi, moderate group in Fig. 6B), suggesting a role of DVGs in stimulating primary IFNs early. In support, ISGs showed similar trend. As IFN related genes are zero-inflated, we performed comparisons for both the expression level of cells expressing interested genes (gene counts >0, named as non-zero cells) and their percentages within DVG+ and DVG− groups. Briefly, the average expression of ISGs (non-zero cells) was all significantly enhanced in DVG+ cells within moderate group at 2 dpi but this enhancement was partially lost at 3 dpi despite of higher percentage of DVG+ cells expressing IFNs and ISGs at 3 dpi relative to that of DVG− cells (Fig. 6C and 6D). Different from moderate group, high viral group had minimal expression of all IFN related genes, further confirming IFN pathways were suppressed in highly infected cells (Fig. 6A, 6B). Low viral group predominantly expressed ISGs rather than two primary IFNs at all time points (Fig. 6E), suggesting they are the secondary responders to initial type I/III IFN production. Taken together, our analysis strongly suggests that DVG+ cells with moderate infection were the first responders to viral infection, quickly expressing primary IFNs and subsequentially alerting neighboring cells to express ISGs.

**Figure 5.**
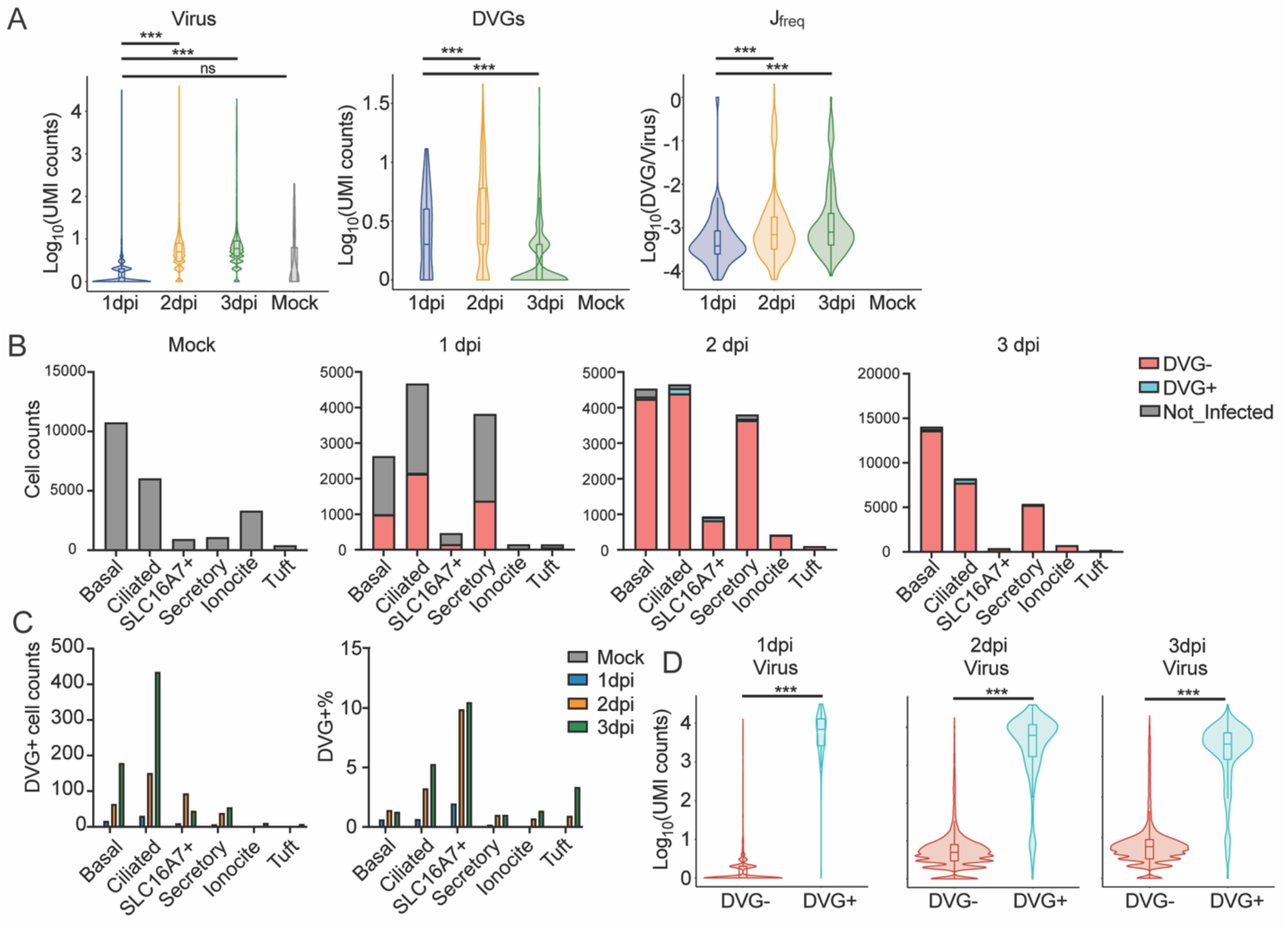
DVG generation in infected NHBE cells from single cell level. **(A)** Violin plots of log transformed viral UMI counts, DVG UMI counts, and J_freq_ for 1 dpi, 2 dpi, 3 dpi, and mock samples. **(B)** Bar plots of cell counts of uninfected cells, DVG− infected cells, and DVG+ cells within different cell type for mock, 1 dpi, 2 dpi, 3 dpi samples. Infected cells were cells with viral UMI over 1 and DVG+ cells were the ones with DVG UMI over 1. All DVG+ cells had at least 1 viral UMI. **(C)** Bar plots of DVG+ cell counts and DVG+ percentages per cell type for mock, 1 dpi, 2 dpi, and 3 dpi samples. **(D)** Violin plots of log transformed viral counts for DVG+ and DVG− viral positive cells. *** p < 0.01, ** p < 0.05 by two-sided Wilcoxon Rank Sum test.

**Figure 6.**
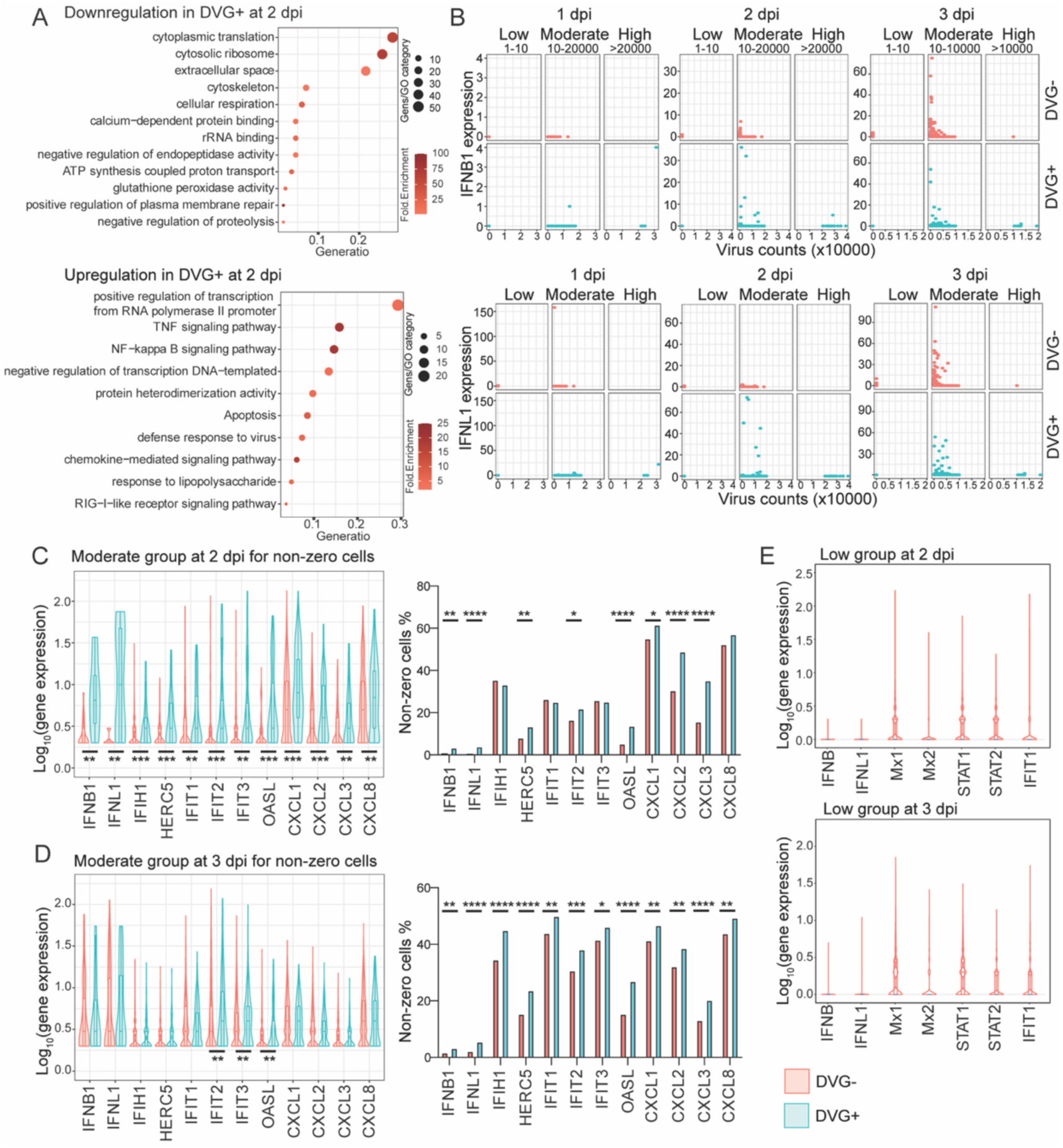
DVG+ cells expressed primary IFNs earlier than DVG− cells. **(A)** Gene ontology analysis of genes that were downregulated (Top) and upregulated (Bottom) in DVG+ cells relative to DVG− cells at 2 dpi. Circle size represented number of genes in each pathway. Gene ratio represented the ratio of number of genes in that pathway to the number of genes in the entire cluster. **(B)** Gene expression of IFNB1 and IFNL1 (Y-axis) were correlated with viral UMI level (X-axis) within each virus counts group. Virus groups with their counts criteria were indicated on top of the graph. Each dot represented individual cell and were colored based on their presence of DVGs. **(C-D)** In the moderate virus group, expression level of IFNB, IFNL1, selected ISGs and chemokines for non-zero (gene counts > 0) cells and percentage of non-zero cells within DVG+ and DVG− groups were compared and graphed as violin plots at 2 dpi **(C)** and 3 dpi **(D)**. *** p < 0.01, ** p < 0.05 by two-sided Wilcoxon Rank Sum test. **(E)** Expression level of IFNB, IFNL1, and selected ISGs for DVG− cells with low virus group at 2 dpi and 3 dpi were graphed as violin plots. **** P<0.0001, *** p < 0.001, ** p < 0.01, * p < 0.05 by Fisher’s exact test.

### Symptomatic COVID-19 patients had higher amount and Jfreq of SARS-CoV-2 DVGs than asymptomatic patients

As SARS-CoV-2 DVGs can stimulate early expression of primary IFNs, the question of whether DVG generation is associated to COVID-19 disease severity was asked. We identified a publicly available NGS dataset (PRJNA690577) investigating subgenomic RNAs and their protein expression from symptomatic vs asymptomatic COVID-19 patients and the authors also indicated more deletions with length over 20 nts in symptomatic patients than asymptomatic patients (Wong, Ngan et al. 2021). To better examine the DVG (larger deletions) level between two patient groups, we applied our criteria to this dataset and found a distinguished increased DVG counts (both −sense and +sense, Fig. 7A) and subsequent higher J_freq_ (Fig. 7C) in symptomatic individuals compared to asymptomatic patients on average. Additionally, our method also confirmed the original finding that the read counts for genomic RNA was significantly lower in symptomatic patients than that in asymptomatic patients (Fig. 7B). These data imply the potential role of DVGs in COVID-19 symptom development.

**Figure 7.**
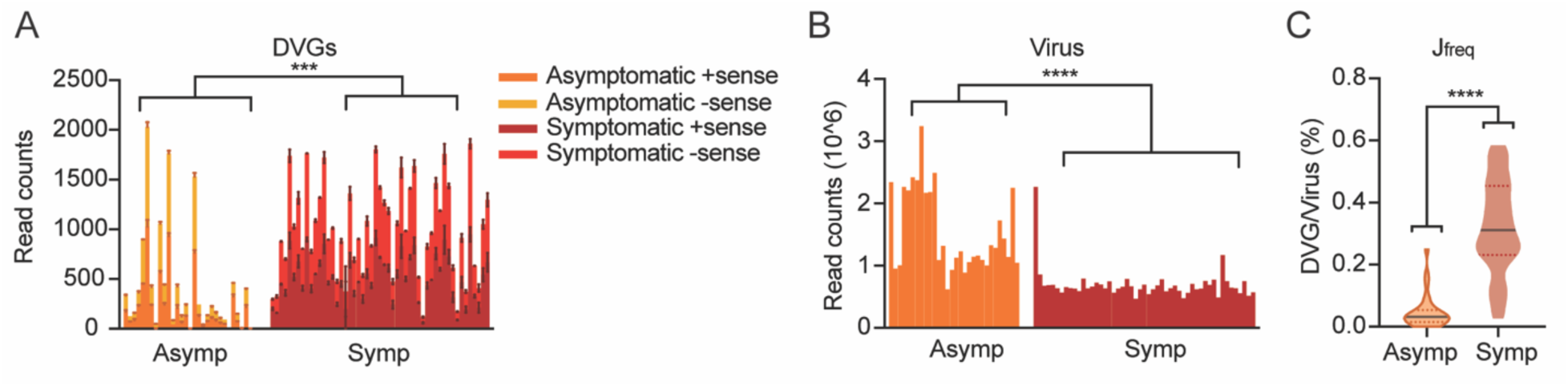
Symptomatic COVID-19 patients had higher amount and Jfreq of SARS-CoV-2 DVGs than asymptomatic patients. Samples of various collection methods including nasopharyngeal (n = 42), anterior nasal (n = 35), and oropharyngeal (n = 5) were used from NGS dataset PRJNA690577. Symptomatic samples (n = 51) were collected from patients presented at the hospital with symptoms consistent with COVID-19 while asymptomatic samples (n = 30) were collected from patients who did not have symptoms consistent with COVID-19 and were found through contact tracing and workforce screening. DVG read counts **(A)**, viral read counts **(B)**, and J_freq_ **(C)** percentages were calculated and graphed for all symptomatic and asymptomatic samples. **** p < 0.0001, *** p < 0.001 by two-sided Mann-Whitney test.

### High DVG Jfreq was observed in one COVID-19 persistent patient

SARS-CoV-2 can develop persistent infections in immunosuppressive patients (Caccuri, Messali et al. 2022, Quaranta, Fusaro et al. 2022), and DVGs have been reported to facilitate viral persistence (Sun and López 2017). To examine whether DVGs are associated with persistent SARS-CoV-2 infection in patient, we identified one NGS dataset, where nasal samples were taken at 9 time points from one immunosuppressive patient who was infected with SARS-CoV-2 and was positive for virus up to 140 days since the first hospital admission (PRJEB47786). We detected DVGs in all 9 time points, but the amount of DVGs were not always correlated with total virus counts (Fig. 8A and 8B). More interestingly, J_freq_ of DVGs from the samples in this patient were at least 10 times higher than the number we observed in *in vitro* infections and autopsy tissues (Fig. 8C vs Fig. 1B, 4A, and 5A) with highest J_freq_ up to nearly 20% at 56 days post initial admission to hospital. We noticed that the method used in this dataset was tiled-PCR using ARTIC V3 followed by Illumina sequencing, which is different from all the previous bulk and single cell RNA-seq we examined. To test whether the high J_freq_ was due to the different approach and potentially because of nasal samples, we found another NGS dataset with nasal samples of normal COVID-19 patients using tiled-PCR (ARTIC V1 and V3) followed by Illumina sequencing (PRJNA707211). We found that the J_freq_ of each patient sample was below 1%, within the range observed from previous in vitro and autopsy NGS (Fig. 8D vs Fig. 1B, 4A, and 5A). This strongly suggests that the high J_freq_ of DVGs in this patient was not due to the amplification and sequencing methods, but rather may be associated with the suppression status of patient’s immune system and persistent viral infection.

**Figure 8.**
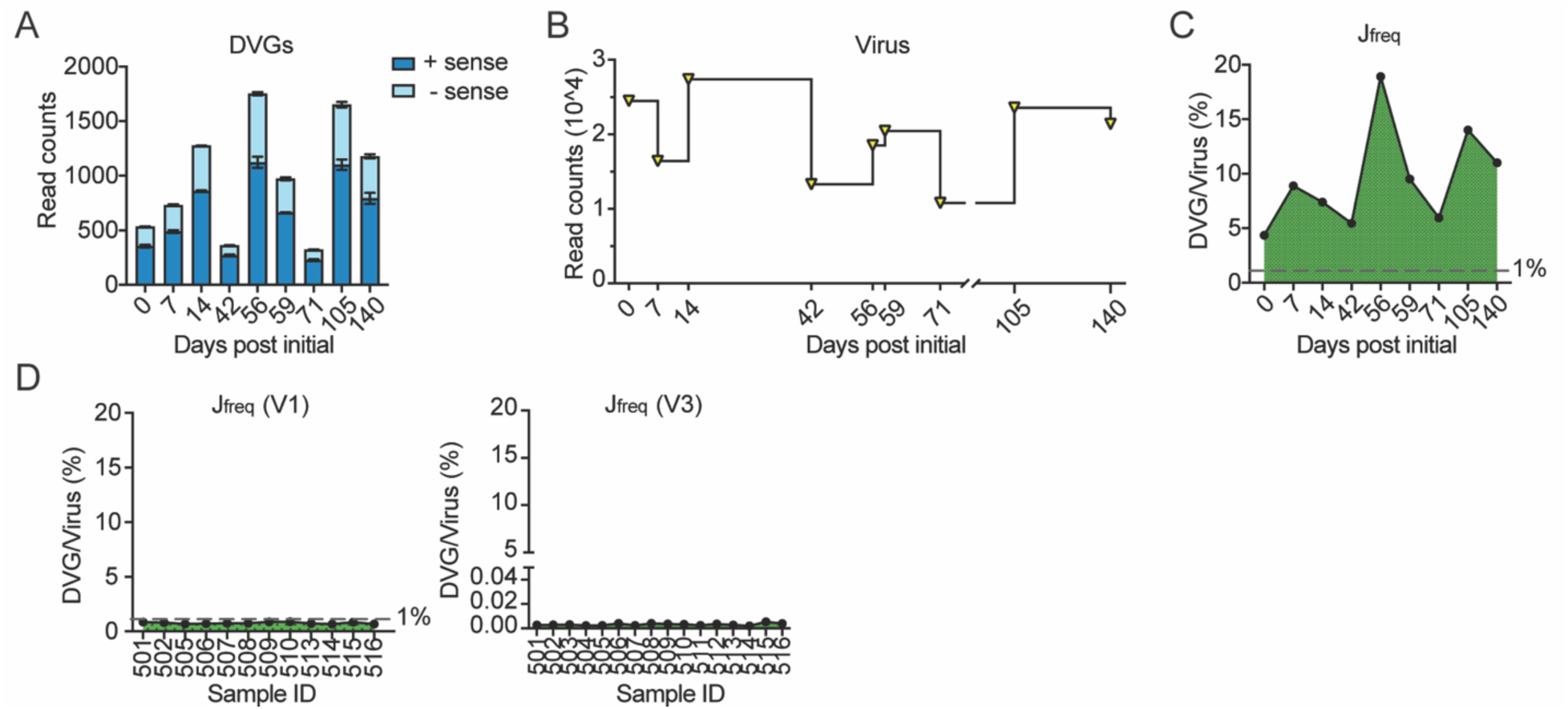
High DVG Jfreq was observed in one SARS-CoV-2 persistent patient. Nasal samples were collected from one immunosuppressive COVID-19 patient with persistent viral infection at 9 different time points. DVGs were identified from the NGS dataset (ERP132087/PRJEB47786) of the nasal samples from this patient. DVG read counts **(A)**, viral read counts **(B)**, and J_freq_ **(C)** percentages were calculated and graphed for samples at each time points. **(D)** J_freq_ of samples in another NGS dataset (PRJNA707211) utilizing the same amplification and sequencing methods demonstrated a much smaller J_freq_ than the SARS-CoV-2 persistent patient, comparable to J_freq_ levels found SARS-CoV-2-infected in vitro and autopsy samples.

## Discussion

It has been well-documented that DVGs are universally generated across single stranded RNA viruses both *in vitro* and *in vivo*, such as Respiratory Syncytial Virus (RSV), measles, influenza, Ebola, Dengue, CoVs, and many more. For SARS-CoV-2, DVGs are resulted from non-homologous recombination and are previously observed in infected Vero cells (Chaturvedi, Vasen et al. 2021) and nasal samples of COVID-19 patients (Xiao, Lidsky et al. 2021). In Vero cells, SARS-CoV-2 is reported to be more than 10 times more recombinogenic than other CoVs, such as MERS-CoV (Gribble, Stevens et al. 2021) and junctions of SARS-CoV-2 DVGs are most commonly flanked at U-rich RNA sequences, suggesting a novel mechanism by which viral polymerases use to generate DVGs. Interestingly, recombination is also proposed to be critical for coronavirus diversity and emergence of SARS-CoV-2 and other zoonotic CoVs. To further understand the recombination positions of SARS-CoV-2 DVGs, we expanded DVG analyses to 4 more commonly used cells lines for SARS-CoV-2 studies, primary human lung epithelial cells (NHBE), and autopsy tissues from patients died of complications of COVID-19, further confirming that DVGs are ubiquitously produced during SARS-CoV-2 infections. Importantly, we identified specific genomic hotspots for DVG recombinant sites that are not only consistent in *in vitro* and in patient samples, but also shared between +sense and −sense DVGs. These results imply two points: 1) DVG recombination is not random in SARS-CoV-2 and certain mechanisms are utilized to regulate their production; and 2) our identified +sense DVGs and −sense DVGs are correlated with each other, likely due to the self-replication in between. One limitation of our analyses using short-read NGS is that short reads are <400 bp long and thus junction reads are less likely to cover the entire DVG sequence. Despite of this, the replication capability of identified DVGs strongly suggest that the 5’ UTR and 3’ UTR are retained in our identified DVGs, as two UTRs are essential for genome replication. More analysis from long read sequencing data are needed to further confirm full sequences of DVGs.

Based on the secondary structures identified by COMRADES crosslinking in the +sense viral genome (Ziv, Gabryelska et al. 2018), we calculated the structural distance between two recombination sites of any −sense DVGs and surprisingly found an association between DVG break and rejoin points with short structural distance (Fig. 3C, D), as mediated by RNA base pairing. The relatively short structural distance, as compared to the sequence length, indicates that DVGs form when the viral polymerase falls off the template during replication and then rejoins the viral template at a position close in space, which can be quite distant in sequence. This strongly suggests that the recombination of viral polymerase complex can be guided by the secondary structures within viral genomes. As the structures formed within the −sense strand are expected to be different from those in +sense strand (because folding stability is strand-direction dependent and G-U pairs map to A-C mismatches in the complementary strand), we postulate that DVG generation is initiated as −sense by viral polymerase complex using +sense viral genomes as template and −sense DVGs are then used as templates to replicate +sense DVGs (Fig. S1E). More investigations on the secondary structures in both strands of viral genomes and their role in viral recombination are needed to further test this hypothesis.

The presence of DVGs on host response and viral replication were additionally explored. It was observed that samples with moderate and high amounts of DVGs exhibited enhanced antiviral responses than samples with low amounts of DVGs. From scRNA-seq analysis, IFN pathways were suppressed in highly infected cells and primary IFNs were stimulated earlier in moderately infected cells with DVGs than the ones without DVGs. These data suggest DVG generation earlier on in infection can enhance antiviral response more quickly, which is critical for mounting adequate and in-time immune response. The mechanisms by which DVGs enhance IFN responses are unknown. DVGs from RSV and influenza can function as primary triggers to directly stimulate type I IFN production through RIG-I like receptors (Sun and López 2017). It is previously reported that SARS-CoV-2 RNAs can be recognized by MDA5 (Thorne, Reuschl et al. 2021, Znaidia, Demeret et al. 2022) and we showed that the expression of MDA5 (IFIH1) was elevated in DVG+ cells at 2 dpi (Fig. 6C). Therefore, it is possible that SARS-CoV-2 DVGs stimulate type I/III IFNs through MDA5. Alternatively, if DVGs do not directly stimulate IFN production, they can suppress the expression of viral-encoding IFN antagonists by large deletions, resulting in an earlier and higher IFN expression in DVG+ cells. Indeed, IFN antagonists are encoded in NSP1, NSP3, NSP5, NSP12, NSP13, NSP14, NSP15, ORF3a, ORF3b, ORF6, ORF7a, ORF7b, ORF8, ORF9b, N, and M (Lei, Dong et al. 2020, Xia, Cao et al. 2020, Han, Zhuang et al. 2021, Wong, Cheung et al. 2022, Znaidia, Demeret et al. 2022) and most of them are within the deletion regions based on our conserved genomic hotspots for DVG recombination sites (Fig. 2A and 2B). Nevertheless, the higher IFN expression in DVG+ samples/cells suggest the critical role of DVGs in modulating host responses and sequential disease severity of COVID-19.

To further explore the role of DVGs in COVID-19 severity, we take advantage of one published NGS dataset that investigates sgmRNA levels in patients with differing clinical severity (Wong, Ngan et al. 2021). They observed a reduction of viral sgmRNAs and viral deletions larger than 20 nts but an increased viral genomic RNA level in nasal samples from asymptomatic patients. As deletions with a cutoff of 20 nts may not represent the viral genomes that are defective, we applied our criteria to this dataset and found that the abundance and J_freq_ of DVGs containing deletions larger than 100 nts were similarly reduced in asymptomatic patients compared to symptomatic patients. A significant difference in DVG production between patients with and without symptoms leads us to posit that quantity and J_freq_ of DVGs contribute to the heterogeneity of both disease outcomes and presentation of symptoms in infected individuals, potentially through modulating host immune responses. As sgmRNAs and DVGs were both reduced in asymptomatic group in this cohort study, we wonder whether sgmRNAs production is always positively correlated with DVG generation. To examine this, we quantified TRS-dependent junction reads (recombination sites <85) from the ViReMa output in infected PHLE cells from different age groups as the estimation of sgmRNAs (dataset used in Fig. 4). Interestingly, we did not observe any positive correlation. Specifically, D198 with the least DVG amount among all samples at 72 hpi had more sgmRNAs counts (n=385) than D239 (n=32), which again confirm that DVGs, rather than sgmRNAs, specifically stimulate IFN responses. Why do symptomatic patients generate more DVGs? It is possible that the IFN response induced by DVGs lead to subsequential expression of cytokines, such as IL6, which is known to be an important mediator for immune-induced fever, as shown in blood monocytes for SARS-CoV-2 infection (Junqueira, Crespo et al. 2021). However, rapid and controlled immune response will lead to milder symptoms, whereas prolonged and uncontrolled immune response will lead to severe symptoms and even death (Janssen, Grondman et al. 2021). Future studies with higher symptom scoring resolution, such as mild/moderate, severe, and death, could elucidate the potential associations of DVG abundance and/or frequency with viral load, IFN responses, and COVID-19 disease severity.

Analysis of DVG presence in longitudinal clinical samples describe the kinetics of the DVG population across entire infection course. For one NGS dataset, we were surprised to find one immunosuppressed patient generating DVGs consistently in every collected time point over a period of 140 days, and J_freq_ of these samples being at least 10-fold higher than all previous analyzed datasets (>1%). When comparing a similar method, it was determined that the increased J_freq_ was not due to the amplification and sequencing methods, but rather a biological difference either from a compromised immune status or a prolonged viral infection. These data additionally imply that a prolonged DVG presence/production may associate with a prolonged viral infection and a longer length of illness. Indeed, DVGs have been shown to promote viral persistence for various viruses, such as influenza A (De and Nayak 1980), dengue (Juárez-Martínez, Vega-Almeida et al. 2013), Japanese encephalitis virus (Park, Choi et al. 2013), mumps (Andzhaparidze, Bogomolova et al. 1983), rabies (Kawai, Matsumoto et al. 1975), Sendai (Roux and Waldvogel 1981), measles (Baczko, Liebert et al. 1986); additionally, worse disease outcome was found to be associated with prolonged DVG detection in RSV (Felt, Sun et al. 2021). More longitudinal studies are needed to elucidate the relationship between DVGs and prolonged viral infection especially in immunosuppressed COVID-19 patients.

Determining the generation (recombination) and function of DVGs during SARS-CoV-2 infection would facilitate reducing the viral recombination events, which greatly contribute to newly emerging CoVs, and elucidate another point of mitigating disease severity from those infected. We showed here that the recombination sites of SARS-CoV-2 DVGs are clustered in several genomic regions, which are likely to be determined by RNA secondary structures formed in between. Furthermore, our studies provide the evidence that DVGs play vital roles in IFN stimulation, prolonged viral replication, and symptom development during SARS-CoV-2 infection, urging for more investigations to further determine the mechanism of DVG generation and their impact on SARS-CoV-2 pathogenesis.

## Materials and Methods

### Virus and cell preparation

The following reagents were deposited by the Centers for Disease Control and Prevention and obtained through BEI Resources, NIAID, NIH: SARS-Related Coronavirus 2, Isolate USA-WA1/2020, NR-52281. SARS-CoV-2 was propagated and titered using African green monkey kidney epithelial Vero E6 cells (American Type Culture Collection, CRL-1586) in Eagle’s Minimum Essential Medium (Lonza, 12-125Q) supplemented with 2% fetal bovine serum (FBS) (Atlanta Biologicals), 2 mM l-glutamine (Lonza, BE17-605E), and 1% penicillin (100 U/ml) and streptomycin (100 μg/ml). Viral stocks were stored at − 80°C. All work involving infectious SARS-CoV-2 was performed in the Biosafety Level 3 (BSL-3) core facility of the University of Rochester, under institutional biosafety committee (IBC) oversight.

### PHLE culture on air-liquid interface and SARS-CoV-2 infection

Primary human lung epithelial (PHLE) cells were cultured on an air-liquid interface as previously described (Wang, Bhattacharya et al. 2020, Anderson, Chirkova et al. 2021). Briefly, lung tissue issues were digested with a protease cocktail and cells were then cultured on a collagen-coated transwell plate (Corning, 3470) until each well reaches a transepithelial electrical resistance (TEER) measurement of >300 ohms. Cells were then placed on an air-liquid interface (ALI) by removing media from the apical layer of the transwell chamber and continuing to feed cells on the basolateral layer as they differentiate. Cells were differentiated for 4-5 weeks at ALI before use in experiments. The apical layer of primary lung cells that had been cultured on an air-liquid interface for about 4-5 weeks were inoculated with SARS-CoV-2 (BEI, NR-52281, hCoV-19/USA-WA1/2020) at a MOI of 5 (titered in VeroE6 cells) in phosphate-buffered saline containing calcium and magnesium (PBS++; Gibco, 14040-133), and incubated at 37°C for 1.5 hours. The infectious solution was then removed and the apical layer washed with PBS++. Cells were then incubated for 24, 48, or 72 hours.

### SARS-CoV-2 inactivation and sample preparation

Cells that were harvested at 24 and 72 hours post infections were lysed with SDS lysis buffer (50mM Tris pH8.0, 10mM EDTA, 1% SDS) and collected with a wide-bore pipette tip. Cells that were harvested at 48 hours were first washed by dispensing and aspirating 37°C HEPES buffered saline solution (Lonza, CC-5022), and then trypsinized with 0.025% Trypsin/EDTA (Lonza, CC-5012) for 10 min at 37°C. Dissociated cells were aspirated using a wide-bore pipette tip and to a tube containing ice-cold Trypsin Neutralization Solution (Lonza, CC5002); this was repeated to maximize cell collection. Cells were then pelleted by centrifugation, resuspended in chilled HEPES, and centrifugally pelleted once more before being resuspended in SDS lysis buffer. All samples were physically lysed with QIAshredder homegenizers (Qiagen, 79656) and stored at - 80°C. Homogenized SDS lysates were diluted 1:1 with RNA lysis buffer (Agilent) and RNA was extracted using the Absolutely RNA Microprep Kit (Agilent) according to the manufacturer’s protocol, including on-column DNase treatment.

### Bulk RNA-sequencing of infected PHLE cells

RNA concentration was determined with the NanopDrop 1000 spectrophotometer (NanoDrop, Wilmington, DE) and RNA quality assessed with the Agilent Bioanalyzer 2100 (Agilent, Santa Clara, CA). 1 ng of total RNA was pre-amplified with the SMARTer Ultra Low Input kit v4 (Clontech, Mountain View, CA) per manufacturer’s recommendations. The quantity and quality of the subsequent cDNA was determined using the Qubit Flourometer (Life Technologies, Carlsbad, CA) and the Agilent Bioanalyzer 2100 (Agilent, Santa Clara, CA). 150 pg of cDNA was used to generate Illumina compatible sequencing libraries with the NexteraXT library preparation kit (Illumina, San Diego, CA) per manufacturer’s protocols. The amplified libraries were hybridized to the Illumina flow cell and sequenced using the NovaSeq6000 sequencer (Illumina, San Diego, CA). Single end reads of 100nt were generated for each sample.

### Bulk RNA-seq data processing and DVG identification

The datasets used for bulk RNA-Seq analyses in Fig. 1 and Fig. 2 were publicly available. Their detailed information was listed in Table S1. The RNA-seq used in Fig. 4 were from our own infection following the protocol as demonstrated earlier. For each sample, we first used Bowtie2 (v. 2.2.9, (Langmead and Salzberg 2012)) to align the reads to the GRCh38 human reference genome. The unmapped reads were then applied to ViReMa (Viral-Recombination-Mapper v. 0.21) to identify recombination junction sites and their corresponding read counts using SARS-CoV-2 reference genome (GenBank ID MT020881.1). A custom filtering script was developed in R to identify junction reads that met our criteria (R v4.1.0 and RStudio v1.4.17, script in Fig. S6). We required the positions of both sites (break and rejoin) of junction reads larger than 85, as TRS-L is reported to be located with the first 85 nts of the SARS-CoV-2 genome. Additionally, we required deletions longer than 100 nts to ensure that the truncated viral RNAs are deficient in replication. We also included all deletions that had one or more reads as identified by ViReMa. The number of viral reads in each bulk RNA-Seq sample was quantified using the RSubread Bioconductor package. The junction frequency (J_freq_) was calculated as shown below for each sample.

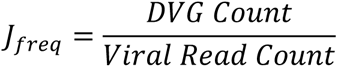

For host transcriptome analysis, raw fastq files were mapped to the human transcriptome (cDNA; Ensembl release 86) using Kallisto (Bray, Pimentel et al. 2016) with 60 bootstraps per sample. Annotation and summarization of transcripts to genes was carried out in R, using the TxImport package (Soneson, Love et al. 2015). Differentially expressed genes (≥twofold and ≤ 1% false discovery rate) were identified by linear modeling and Bayesian statistics using the VOOM function in the Limma package (Ritchie, Phipson et al. 2015). Gene Ontology (GO) was performed using the Database for Annotation, Visualization and Integrated Discovery (DAVID) (Dennis, Sherman et al. 2003).

### DVG identification from scRNA-seq dataset

We used the publicly available dataset from Ravindra et al. 2021 accessed through the NCBI database (GSE166766). This study consisted of single cell RNA-Seq (scRNA-Seq) data from human bronchial epithelial cells (NHBEs) infected with SARS-CoV-2 that were harvested 1 day post infection (dpi), 2 dpi, and 3 dpi. We first used Cell Ranger (Zheng, Terry et al. 2017) to construct gene expression matrices for each sample. To identify the number of viral transcripts, the SARS-CoV-2 reference sequence was concatenated to the end of the human genome reference as one additional gene. The gene expression matrices were then loaded into the Seurat package in R (Satija, Farrell et al. 2015), followed by principal component analysis and cell clustering were performed. Cells were then clustered and annotated based on the gene makers used in the original publication of this dataset. To identify DVGs, we first used UMI-tools (Smith, Heger et al. 2017) to associate the cell barcodes and UMIs with each corresponding read name. Similar to the bulk RNA-Seq analysis, we used Bowtie2 (Langmead and Salzberg 2012), ViReMa, and a custom R filtering script for DVG identification (details in Fig. S6). We then used the filtered ViReMa output to re-quantify DVG count based on the UMIs associated with each cell barcode, which is considered as DVG count per cell. We also calculated J_freq_ for each cell by using DVG UMI/viral UMI per cell barcode. The numbers of DVG UMIs and J_freq_ of each cell barcode was then added to the gene expression matrix created by Cell Ranger. The J_freq_ values were multiplied by 10^3^ so that they would not be left out during the cell clustering and type identification steps. Cells with more than one DVG UMI (virus positive cells) were grouped as DVG+ and DVG− based on the presence or absence of DVG UMI, respectively.

### Differentially expressed genes between DVG+ and DVG− in scRNA-seq analysis

The list of differentially expressed genes between the DVG+ group and DVG− group was generated with the Seurat function FindMarkers, after normalizing and scaling the data with the Seurat function SCTransform. Three different types of tests were used to create three differential gene expression (DGE) lists for both 2 dpi and 3 dpi: Mast, DESeq2, and the Wilcoxon rank sum test (default) using the criteria of percentage of cells where the gene was detected (pct) > 0.1, adj_pval < 0.01, and log fold change > 0.25. The final DGE list was determined based on common genes that were found in two of the three methods. To identify the pathways enriched in the DGE list, we first divided the DGE list based on their upregulation and downregulation in DVG+ group. GO analysis was performed for the upregulated genes and downregulated genes separately through DAVID functional annotation clustering tool and graphed in R using the code in Fig. S6. We then specifically focused on interferon responses between DVG+ and DVG− groups. Low, medium and high groups were further categorized based on their amount of viral UMI within virus positive cells and the expression of selected IFN related genes were specifically compared and graphed between DVG+ and DVG− cells within each viral groups in R (code in Fig. S6).

### DVG identification from the tiled-PCR deep sequencing

The protocol for identifying DVGs in three publicly available datasets that utilize PCR tiling from ARTIC LoCost (V1 or V3) (https://artic.network) primer sets followed bulk sequencing data processing for DVG identification. The first dataset was used to study DVG generation during longitudinal COVID-19 persistence in one immunosuppressed patient (ENA: ERP132087, NCBI SRA: PRJEB4778) and the second one was served as the control cohort containing 16 regular COVID-19 patients using the same way to prepare the library (PRJNA707211). The third one is to study DVGs in a cohort of both asymptomatic and symptomatic COVID-19 patients (NCBI SRA: PRJNA690577). This method of amplification produced overlapping 400 bp amplicons that are then used to construct respective sequencing libraries from which data processing and subsequent analysis can occur. For the longitudinal study, the ARTIC V3 amplicons were sequenced as paired-end 300 bp reads on Illumina Miseq. The ARTIC V3 amplicons of the symptomatic cohort study was PCR amplified by five cycles and also sequenced identically.

### Secondary structure analysis of DVG junction positions

Our definition of structural distance follows (Clote, Ponty et al. 2012). For a given primary sequence and a corresponding secondary structure, we first convert them to a graph where each nucleotide i is a node. We add an edge (i, i+1) between any two adjacent nucleotides i and i+1 (gray bonds in Fig. 3B), and an edge (i, j) between any paired bases i and j (black bonds in Fig. 3B) as reported by Ziv et al. from their COMRADES mapping (Ziv, Gabryelska et al. 2018). This graph can model alternative base pairs. For example, if nucleotide i has possible pairs with nucleotides j, k, and l, then node i will connect five edges (i, i-1), (i, i+1), (i, j), (i, k), and (i, l). Based on the connected graph, the structural distance between two nucleotides is formalized as the number of edges in the shortest path between them (red solid path in Fig. 3B), which can be solved by classical graph algorithms (Cormen, Leiserson et al. 2022).

The chimeric reads detected by COMRADES from (Ziv, Price et al. 2020) consist of only left- and right-side sequences without base-pairing information. For short-range interactions, they extracted a (continuous) subsequence between the 5’ end of the left side and the 3’ end of the right side and used RNAfold (Lorenz, Bernhart et al. 2011) to predict structures for that subsequence. For long-range interactions, they utilized RNAduplex (Lorenz, Bernhart et al. 2011) to predict interactions between the two (distant) segments, which does not model any intra-segmental base pairs for either segment. Note that alternative base pairs exist in the data. Therefore, we built the graph based on the predicted base pairs in Ziv et al.’s data and calculated the structural distance between any two positions using the method described above. Additionally, we chose a cutoff value of 50 for the number of chimeric reads, which leads to a balanced precision and sensitivity evaluated on the known structure (Ziv, Gabryelska et al. 2018).

### Statistical analysis

Pearson’s correlation was performed to identify the association between virus and DVG counts and virus and J_freq_ in the bulk RNA-Seq datasets. For the scRNA-Seq dataset, unpaired two-sided Wilcoxon rank sum tests were performed to identify the differences in viral load, DVG counts, and J_freq_ among mock, 1 dpi, 2 dpi, and 3 dpi samples. We first log transformed viral UMI counts and expression level of selected IFN related genes and then compared between DVG− and DVG+ cells for each time point using unpaired two-sided Wilcoxon rank sum tests.

### Data availability

Source data for the publicly available NGS datasets described in this manuscript is available as Supplementary Table S1. All NGS datasets were retrieved with NCBI and ENA accession numbers GSE147507 (Daamen, Bachali et al. 2021), GSE148729 (Wyler, Mösbauer et al. 2021), BioProject PRJNA628043 (Ogando, Dalebout et al. 2020), GSE166766 (Ravindra, Alfajaro et al. 2021), GSE150316 (Desai, Neyaz et al. 2020), BioProject PRJNA707211 (Jaworski, Langsjoen et al. 2021), and BioProject PRJNA690577 (Wong, Ngan et al. 2021); ERP132087-BioProject PRJEB47786 (Weigang, Fuchs et al. 2021), respectively. Dataset used in Fig. 4 are available upon request and the raw data of all infected samples are under submission to GEO.

## Acknowledgments

We would like to thank the lab group of Dr. Andrew Routh from UTMB for assistance in setting up ViReMa for our analysis of DVG generation. We would like to thank Dr. Xing Qiu from University of Rochester for statistical advice. Publicly available datasets provided by the following lab groups listed are especially recognized: Chandam Deshpande, Landthaler, Lipsky, and Wilen. The authors want to acknowledge the contributions of Gloria S. Pryhuber, M.D., University of Rochester Center for Advanced Research Technologies, the University of Rochester Genomics Research Center (GRC), the Biosafety Level 3 program, the University of Rochester Biosafety Level 3 (BSL3) core facility, and the University of Rochester’s Institutional Biosafety Committee (IBC). We thank Sara Ali, University of Rochester, for help in discussion and correlations between −sense and +sense DVGs.

## Author Contributions

PHLE infection and bulk RNA-sequencing: R.M.O., C.S.A.

Secondary structure analysis: S.L., D.H.M., L.H.

DVG analysis and graphing for all bulk RNA-seq: T.Z., Y.S.

DVG analysis and graphing for single cell RNA-seq: T.Z., S.S., Y.S.

DVG analysis and graphing for tiled-PCR sequencing: N.J.G.

Manuscript writing: N.J.G, S.L., Y.S.

Funding support: T.J.M., J.T., S.D., Y.S.

Supervision: T.J.M., J.T., S.D., L.H., Y.S.

## Supplementary tables and figures

**Table S1.** Summary of all samples from published datasets.

| Cells/Tissues/Sample Types | Infection Type | MOI | Time Points | Dataset# (Sample#) | Sequence Method | Paired or Single end |
| --- | --- | --- | --- | --- | --- | --- |
| A549-ACE2<br>Fig. 1B, D, & E<br>Fig. 2 | In vitro<br>(infected cells) | 2 | 24h | GSE147507<br>(GSM4486160,<br>4486161, 4486162) | Bulk | Single |
| NHBE<br>Fig. 1B, D, & E<br>Fig. 2 | In vitro<br>(infected cells) | 2 | 24h | GSE147507<br>(GSM4432381,<br>4432382, 4432382) | Bulk | Single |
| Calu3_total RNA<br>Fig. 1B, D, & E<br>Fig. 2 | In vitro<br>(infected cells) | 0.3 | 24h | GSE148729<br>(GSM4477962,<br>4477963) | Bulk | Paired |
| Calu3_polyA<br>Fig. 1B, D, & E<br>Fig. 2 | In vitro<br>(infected cells) | 0.3 | 24h | GSE148729<br>(GSM4477910,<br>4477911) | Bulk | Single |
| Caco2<br>Fig. 1B, D, & E<br>Fig. 2 | In vitro<br>(infected cells) | 0.3 | 24h | GSE148729<br>(GSM4477888,<br>4477889) | Bulk | Single |
| H1299<br>Fig. 1B, D, & E<br>Fig. 2 | In vitro<br>(infected cells) | 0.3 | 24h | GSE148729<br>(GSM4477868,<br>4477868) | Bulk | Single |
| Vero E6_S<br>Fig. 1B, D, & E<br>Fig. 2 | In vitro<br>(supernatants) | 3 | 48h/<br>passage | PRJNA628043:<br>SRP258466 | Bulk | Paired |
| NHBE<br>Fig. 4<br>Fig. 5 | In vitro<br>(infected cells) | ~0.01 | 24h, 48h,<br>72h | GSE166766 | scRNA-seq | Paired |
| Case1<br>Fig. 1C, D, & E<br>Fig. 2 | Autopsy lung<br>tissue | - | - | GSE150316<br>(GSM4546576,<br>4546577, 4546578,<br>4546579) | Bulk | Paired |
| Case8<br>Fig. 1C, D, & E<br>Fig. 2 | Autopsy lung tissue | - | - | GSE150316<br>(GSM4698544, 4698545, 4698546, 4698547, 4698548) | Bulk | Paired |
| Case9<br>Fig. 1C, D, & E<br>Fig. 2 | Autopsy lung tissue | - | - | GSE150316<br>(GSM4698549, 4698550, 4698551, 4698552, 4698553) | Bulk | Paired |
| Case11<br>Fig. 1C, D, & E<br>Fig. 2 | Autopsy lung tissue | - | - | GSE150316<br>(GSM4698526, 4698527, 4698528) | Bulk | Paired |
| CaseC<br>Fig. 1C, D, & E<br>Fig. 2 | Autopsy lung tissue | - | - | GSE150316<br>(GSM4698556) | Bulk | Paired |
| CaseD<br>Fig. 1C, D, & E<br>Fig. 2 | Autopsy lung tissue | - | - | GSE150316<br>(GSM4698557) | Bulk | Paired |
| CaseE<br>Fig. 1C, D, & E<br>Fig. 2 | Autopsy lung tissue | - | - | GSE150316<br>(GSM4698558) | Bulk | Paired |
| Longitudinal samples<br>Fig. 6A, B, & C | Nasal | - | - | ENA: ERP132087,<br>NCBI SRA:<br>PRJEB47786 | ARTICv3-Bulk | Paired |
| ARTIC samples<br>Fig. 6D | Nasal | - | - | PRJNA707211 | ARTIC<br>v1&v3-Bulk | Paired |
| Asymptomatic and Symptomatic samples<br>Fig. 7 | Nasal | - | - | PRJNA690577 | ARTICv3-Bulk | Paired |

**Table S2.** Common DVGs identified from *in vitro* infections.

| GSE147507 |  |  |  |  |
| --- | --- | --- | --- | --- |
| Break Point | Rejoin Point | Counts | Strand | ID |
| 28691 | 1620 | 109 | - | A549-ACE2_r1 |
| 28691 | 1620 | 49 | - | A549-ACE2_r3 |
| 29173 | 27800 | 8 | - | NHBE_r2 |
| 29173 | 27800 | 104 | - | A549-ACE2_r1 |
| 29173 | 27800 | 92 | - | A549-ACE2_r2 |
| 29173 | 27800 | 63 | - | A549-ACE2_r3 |
| 29307 | 731 | 129 | - | A549-ACE2_r1 |
| 29307 | 731 | 115 | - | A549-ACE2_r2 |
| 29307 | 731 | 96 | - | A549-ACE2_r3 |
| 29308 | 733 | 52 | - | A549-ACE2_r1 |
| 29308 | 733 | 57 | - | A549-ACE2_r2 |
| 29308 | 733 | 46 | - | A549-ACE2_r3 |
| 29310 | 747 | 76 | - | A549-ACE2_r1 |
| 29310 | 747 | 109 | - | A549-ACE2_r2 |
| 29310 | 747 | 48 | - | A549-ACE2_r3 |
| 29310 | 755 | 55 | - | A549-ACE2_r1 |
| 29310 | 755 | 56 | - | A549-ACE2_r2 |
| 29310 | 755 | 54 | - | A549-ACE2_r3 |
| 29310 | 827 | 89 | - | A549-ACE2_r1 |
| 29310 | 827 | 99 | - | A549-ACE2_r2 |
| 29310 | 827 | 49 | - | A549-ACE2_r3 |
| 29350 | 824 | 66 | - | A549-ACE2_r1 |
| 29350 | 824 | 110 | - | A549-ACE2_r2 |
| 29350 | 824 | 92 | - | A549-ACE2_r3 |
| 29353 | 734 | 13 | - | NHBE_r2 |
| 29353 | 734 | 104 | - | A549-ACE2_r1 |
| 29353 | 734 | 52 | - | A549-ACE2_r3 |
| 29353 | 735 | 64 | - | A549-ACE2_r1 |
| 29353 | 735 | 68 | - | A549-ACE2_r3 |
| 29477 | 730 | 54 | - | A549-ACE2_r2 |
| 29477 | 730 | 68 | - | A549-ACE2_r3 |
| **GSE148729** |  |  |  |  |
| 27234 | 27344 | 14 | + | calu3_totalRNA_AR2 |
| 27234 | 27344 | 13 | + | calu3_totalRNA_BR2 |
| 27341 | 27231 | 11 | - | calu3_polyA_A |
| 27341 | 27231 | 9 | - | calu3_polyA_B |
| 27341 | 27231 | 25 | - | calu3_totalRNA_AR1 |
| 27341 | 27231 | 25 | - | calu3_totalRNA_BR1 |
| 27341 | 27231 | 11 | - | caco2_polyA_A |
| 27341 | 27231 | 24 | - | caco2_polyA_B |
| 27794 | 29175 | 19 | + | calu3_totalRNA_AR2 |
| 27794 | 29175 | 10 | + | calu3_totalRNA_BR2 |
| 27794 | 29176 | 12 | + | calu3_totalRNA_BR2 |
| 27795 | 29175 | 12 | + | calu3_totalRNA_AR2 |
| 27796 | 29195 | 8 | + | calu3_totalRNA_BR2 |
| 27802 | 29175 | 29 | + | calu3_totalRNA_AR2 |
| 27802 | 29175 | 28 | + | calu3_totalRNA_BR2 |
| 27965 | 27231 | 3 | - | calu3_polyA_A |
| 27965 | 27231 | 9 | - | calu3_totalRNA_AR1 |
| 27965 | 27231 | 9 | - | caco2_polyA_B |
| 28318 | 29123 | 8 | + | calu3_totalRNA_AR2 |
| 28318 | 29123 | 13 | + | calu3_totalRNA_BR2 |
| 28319 | 29017 | 12 | + | calu3_totalRNA_AR2 |
| 28319 | 29017 | 7 | + | calu3_totalRNA_BR2 |
| 28408 | 29017 | 11 | + | calu3_totalRNA_AR2 |
| 28408 | 29017 | 9 | + | calu3_totalRNA_BR2 |
| 28673 | 28505 | 9 | - | calu3_totalRNA_BR1 |
| 28673 | 28505 | 5 | - | caco2_polyA_A |
| 28729 | 28464 | 13 | - | caco2_polyA_A |
| 28729 | 28464 | 6 | - | caco2_polyA_B |
| 28731 | 28465 | 13 | - | caco2_polyA_A |
| 28731 | 28465 | 8 | - | caco2_polyA_B |
| 28731 | 28495 | 8 | - | caco2_polyA_A |
| 28731 | 28495 | 5 | - | caco2_polyA_B |
| 29084 | 28318 | 12 | - | calu3_totalRNA_AR1 |
| 29084 | 28318 | 11 | - | calu3_totalRNA_BR1 |
| 29084 | 28318 | 6 | - | caco2_polyA_A |
| 29084 | 28318 | 8 | - | caco2_polyA_B |
| 29164 | 27800 | 4 | - | calu3_polyA_A |
| 29164 | 27800 | 6 | - | caco2_polyA_A |
| 29173 | 27792 | 8 | - | calu3_polyA_B |
| 29173 | 27793 | 6 | - | calu3_polyA_B |
| 29173 | 27793 | 6 | - | caco2_polyA_A |
| 29173 | 27800 | 16 | - | calu3_polyA_A |
| 29173 | 27800 | 15 | - | calu3_polyA_B |
| 29173 | 27800 | 16 | - | calu3_totalRNA_AR1 |
| 29173 | 27800 | 14 | - | calu3_totalRNA_BR1 |
| 29173 | 27800 | 25 | - | caco2_polyA_A |
| 29173 | 27800 | 12 | - | caco2_polyA_B |
| 29173 | 27801 | 3 | - | calu3_polyA_A |
| 29173 | 27801 | 6 | - | caco2_polyA_A |
| 29343 | 6653 | 3 | - | calu3_polyA_A |
| 29343 | 6655 | 5 | - | caco2_polyA_B |
| 29345 | 6635 | 3 | - | calu3_polyA_A |
| 29353 | 6603 | 3 | - | calu3_polyA_B |
| 29353 | 6653 | 7 | - | calu3_totalRNA_AR1 |
| 29481 | 6683 | 3 | - | calu3_polyA_B |

| 29481 | 6683 | 9 | - | calu3_totalRNA_BR1 |
| --- | --- | --- | --- | --- |
| 29493 | 6653 | 3 | - | calu3_polyA_A |
| 29494 | 6653 | 4 | - | calu3_polyA_A |
| 29494 | 6653 | 3 | - | calu3_polyA_B |
| 29495 | 6655 | 10 | - | calu3_totalRNA_BR1 |
| 29520 | 6883 | 3 | - | calu3_polyA_B |
| 29520 | 6883 | 11 | - | calu3_totalRNA_BR1 |
| 29805 | 29686 | 8 | - | caco2_polyA_A |
| 29805 | 29686 | 9 | - | caco2_polyA_B |
| **SRP258466** |  |  |  |  |
| Break Point | Rejoin Point | Counts | Strand | ID |
| 5981 | 6566 | 8 | + | veroE6_L8 |
| 5981 | 6566 | 9 | + | veroE6_s5p2 |
| 5982 | 6566 | 8 | + | veroE6_L8 |
| 5982 | 6566 | 8 | + | veroE6_s5p1 |
| 5982 | 6566 | 7 | + | veroE6_s5p3 |
| 5982 | 6573 | 7 | + | veroE6_s5p1 |
| 6044 | 6525 | 8 | + | veroE6_s5p1 |
| 6045 | 6525 | 8 | + | veroE6_L8 |
| 20272 | 20387 | 8 | + | veroE6_L8 |
| 20272 | 20387 | 9 | + | veroE6_s5p3 |
| 27386 | 29472 | 7 | + | veroE6_s5p3 |
| 27386 | 29473 | 13 | + | veroE6_L8 |
| 27386 | 29473 | 17 | + | veroE6_s5p2 |
| 27785 | 29195 | 9 | + | veroE6_s5p3 |
| 27788 | 29196 | 23 | + | veroE6_L8 |
| 27788 | 29196 | 8 | + | veroE6_s5p1 |
| 27788 | 29196 | 26 | + | veroE6_s5p3 |
| 27793 | 29163 | 11 | + | veroE6_s5p1 |
| 27794 | 29166 | 6 | + | veroE6_s5p3 |
| 27794 | 29175 | 7 | + | veroE6_s5p1 |
| 27794 | 29175 | 9 | + | veroE6_s5p3 |
| 27794 | 29195 | 7 | + | veroE6_L8 |
| 27802 | 29175 | 14 | + | veroE6_L8 |
| 27802 | 29175 | 11 | + | veroE6_s5p3 |
| 28508 | 28676 | 22 | + | veroE6_L8 |
| 28508 | 28676 | 7 | + | veroE6_s5p1 |
| 28508 | 28676 | 11 | + | veroE6_s5p2 |
| 28508 | 28676 | 22 | + | veroE6_s5p3 |
| **PHLE cells in vitro infections (own infection)** |  |  |  |  |
| Break Point | Rejoin Point | Counts | Strand | ID |
| 1363 | 29345 | 6 | + | D231_I_72hr_R1 |
| 1363 | 29353 | 4 | + | D231_I_72hr_R1 |
| 1369 | 29353 | 1 | + | D283_I_72hr_R1 |
| 1416 | 29449 | 3 | + | D231_I_72hr_R1 |
| 1425 | 29444 | 2 | + | D283_I_72hr_R1 |
| 1624 | 29337 | 15 | + | D231_I_72hr_R1 |
| 1624 | 29339 | 1 | + | D231_I_72hr_R1 |
| 27382 | 29472 | 10 | + | D231_I_72hr_R1 |
| 27382 | 29473 | 7 | + | D231_I_72hr_R1 |
| 27382 | 29483 | 1 | + | D231_I_72hr_R1 |
| 27385 | 29473 | 14 | + | D231_I_48hr_R1 |
| 27385 | 29479 | 9 | + | D231_I_72hr_R1 |
| 27385 | 29472 | 7 | + | D231_I_72hr_R1 |
| 27385 | 29473 | 3 | + | D231_I_72hr_R1 |
| 27385 | 29473 | 2 | + | D239_I_48hr_R1 |
| 27386 | 29473 | 12 | + | D231_I_72hr_R1 |
| 27386 | 29476 | 4 | + | D231_I_72hr_R1 |
| 27386 | 29474 | 1 | + | D231_I_72hr_R1 |
| 27386 | 29473 | 9 | + | D239_I_48hr_R1 |
| 27386 | 29468 | 4 | + | D239_I_48hr_R1 |
| 27793 | 29166 | 4 | + | D231_I_48hr_R1 |
| 27793 | 29176 | 1 | + | D231_I_72hr_R1 |
| 27794 | 29175 | 5 | + | D231_I_72hr_R1 |
| 27794 | 29167 | 3 | + | D231_I_72hr_R1 |
| 27794 | 29166 | 2 | + | D231_I_72hr_R1 |
| 27795 | 29166 | 11 | + | D231_I_72hr_R1 |
| 27795 | 29195 | 3 | + | D231_I_72hr_R1 |
| 27795 | 29176 | 2 | + | D231_I_72hr_R1 |
| 27795 | 29175 | 2 | + | D231_I_72hr_R1 |
| 27795 | 29175 | 22 | + | D239_I_72hr_R1 |
| 27796 | 29186 | 10 | + | D231_I_48hr_R1 |
| 27797 | 29167 | 4 | + | D231_I_72hr_R1 |
| 27798 | 29176 | 7 | + | D231_I_72hr_R1 |
| 27800 | 29174 | 1 | + | D239_I_48hr_R1 |
| 27801 | 29175 | 4 | + | D231_I_72hr_R1 |
| 27802 | 29175 | 21 | + | D231_I_48hr_R1 |
| 27802 | 29166 | 12 | + | D231_I_72hr_R1 |
| 27802 | 29175 | 9 | + | D231_I_72hr_R1 |
| 27802 | 29176 | 3 | + | D231_I_72hr_R1 |
| 27802 | 29175 | 28 | + | D239_I_48hr_R1 |
| 27802 | 29175 | 1 | + | D283_I_72hr_R1 |
| 27803 | 29174 | 1 | + | D203_I_72hr_R1 |
| 27803 | 29175 | 10 | + | D231_I_72hr_R1 |
| 27803 | 29172 | 3 | + | D231_I_72hr_R1 |
| 29172 | 27803 | 3 | - | D231_I_72hr_R1 |
| 29173 | 27802 | 1 | - | D203_I_72hr_R1 |
| 29173 | 27800 | 10 | - | D231_I_48hr_R1 |
| 29173 | 27801 | 10 | - | D231_I_72hr_R1 |
| 29173 | 27800 | 7 | - | D231_I_72hr_R1 |
| 29173 | 27793 | 4 | - | D231_I_72hr_R1 |
| 29173 | 27792 | 3 | - | D231_I_72hr_R1 |
| 29173 | 27800 | 12 | - | D239_I_48hr_R1 |
| 29173 | 27800 | 4 | - | D283_I_72hr_R1 |
| 29174 | 27800 | 2 | - | D231_I_72hr_R1 |
| 29175 | 27797 | 2 | - | D231_I_72hr_R1 |
| 29176 | 27802 | 2 | - | D231_I_72hr_R1 |
| 29184 | 27794 | 29 | - | D231_I_48hr_R1 |
| 29443 | 1424 | 1 | - | D283_I_72hr_R1 |
| 29448 | 1415 | 5 | - | D231_I_72hr_R1 |
| 29468 | 27381 | 3 | - | D231_I_72hr_R1 |
| 29471 | 27383 | 1 | - | D231_I_48hr_R1 |
| 29472 | 27382 | 1 | - | D231_I_72hr_R1 |
| 29473 | 27386 | 3 | - | D231_I_72hr_R1 |
| 29473 | 27386 | 1 | - | D239_I_48hr_R1 |
| 29474 | 27389 | 1 | - | D198_I_72hr_R1 |
| 29474 | 27386 | 1 | - | D231_I_72hr_R1 |
| 29475 | 27385 | 1 | - | D231_I_72hr_R1 |
| 29685 | 29808 | 1 | + | D231_I_72hr_R1 |
| 29687 | 29813 | 1 | + | D231_I_72hr_R1 |
| 29689 | 29812 | 1 | + | D231_I_48hr_R1 |
| 29690 | 29813 | 2 | + | D231_I_72hr_R1 |
| 29690 | 29828 | 1 | + | D231_I_72hr_R1 |
| 29691 | 29818 | 3 | + | D231_I_72hr_R1 |
| 29695 | 29814 | 8 | + | D231_I_72hr_R1 |
| 29695 | 29823 | 1 | + | D231_I_72hr_R1 |
| 29695 | 29814 | 4 | + | D239_I_48hr_R1 |
| 29791 | 29681 | 1 | - | D239_I_48hr_R1 |
| 29793 | 29690 | 1 | - | D231_I_72hr_R1 |
| 29805 | 29686 | 10 | - | D231_I_72hr_R1 |
| 29807 | 29688 | 3 | - | D203_I_48hr_R1 |
| 29807 | 29688 | 6 | - | D231_I_48hr_R1 |
| 29807 | 29688 | 7 | - | D231_I_72hr_R1 |
| 29807 | 29688 | 16 | - | D239_I_48hr_R1 |
| 29807 | 29688 | 1 | - | D283_I_72hr_R1 |
| 29810 | 29687 | 13 | - | D231_I_48hr_R1 |
| 29812 | 29686 | 1 | - | D231_I_72hr_R1 |
| 29813 | 29690 | 4 | - | D231_I_72hr_R1 |
| 29813 | 29690 | 15 | - | D239_I_48hr_R1 |
| 29814 | 29693 | 1 | - | D231_I_72hr_R1 |
| 29816 | 29680 | 4 | - | D231_I_72hr_R1 |
| 29817 | 29690 | 1 | - | D231_I_72hr_R1 |
| 29818 | 29686 | 4 | - | D231_I_48hr_R1 |

**Figure S1.**
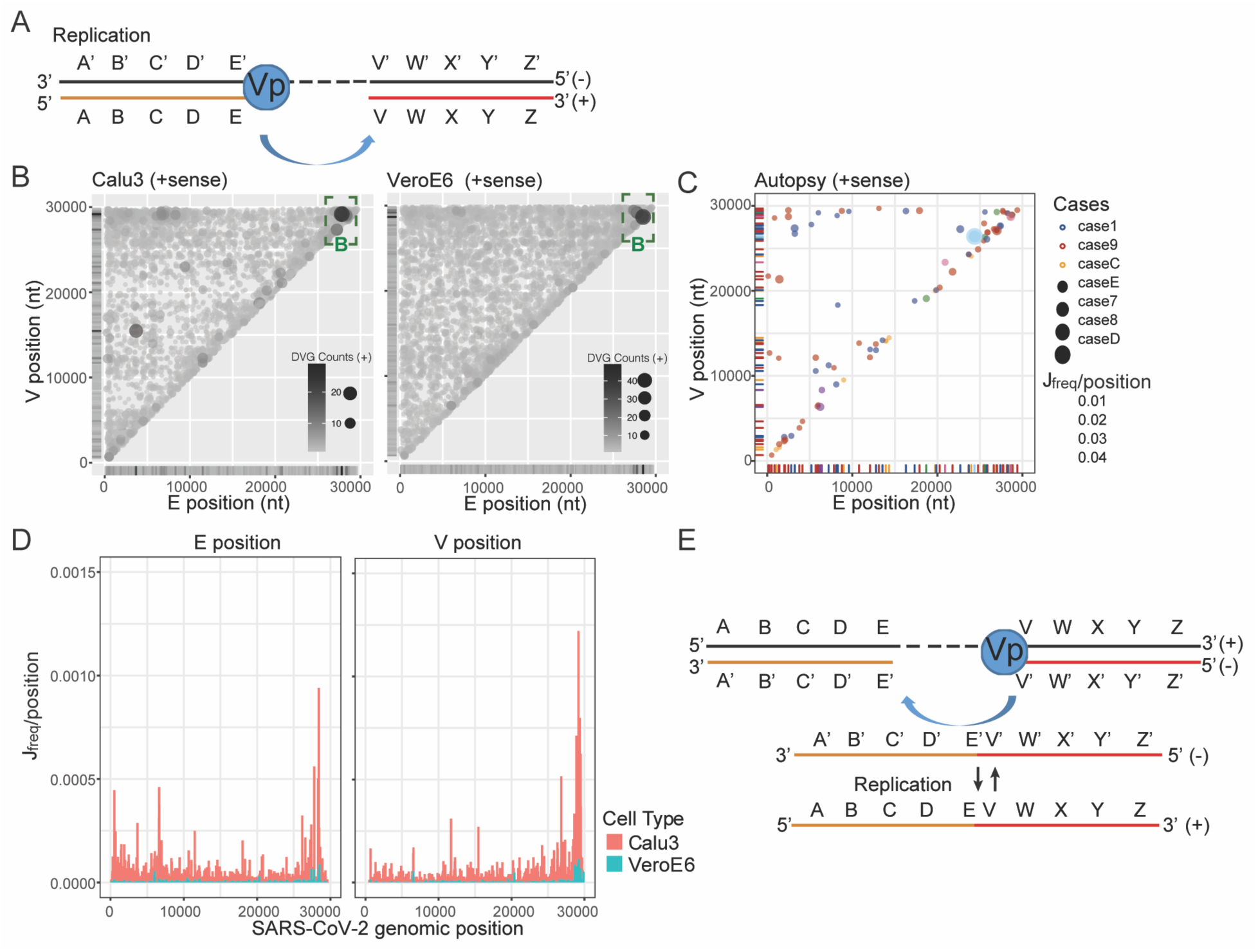
Positive sense DVG generation in SARS-CoV-2 in vitro and autopsy samples. **(A)** scheme for +sense DVGs as they were generated from −sense genomic template. V and E position distributions for +sense DVG from in vitro infected samples **(B),** where circle size and color intensity indicated DVG counts, and autopsy samples **(C)**, where circle size indicated the Jfreq at that position and circle color indicates sample case. The green dashed boxes represented genomic hotspots for DVG junctions. **(D)** V and E position distributions by Jfreq per position for +sense DVGs. Graph showed two in vitro infected samples with more than half of the DVGs are positive sense. The width of each bar represents 300 nucleotides. **(E)** Schematic representation of how - sense and +sense DVGs replicate from each other, leading to the observation that V position of *+sense DVGs shared the same hotspots with V’ position of −sense DVGs.*

**Figure S2:**
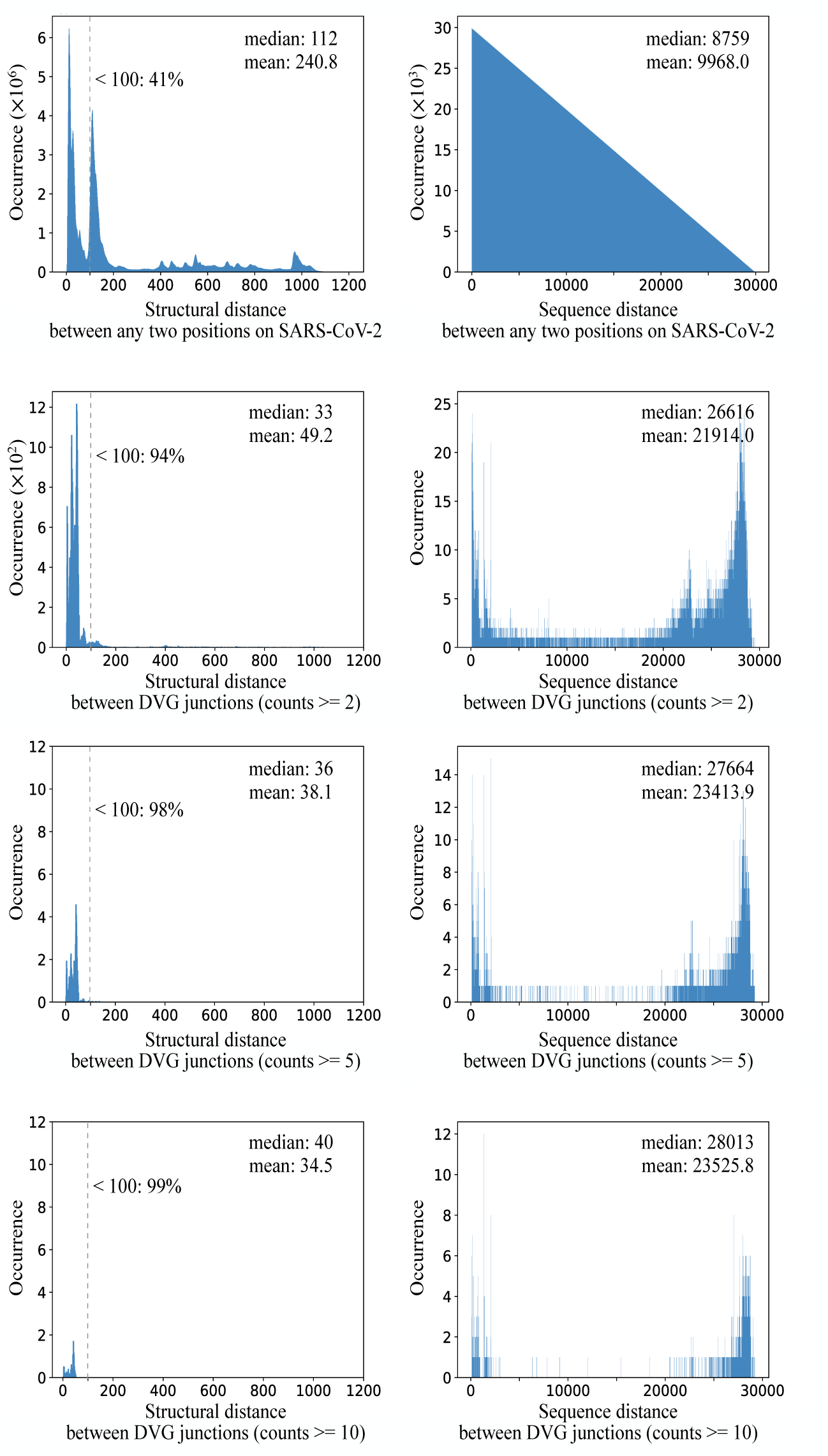
More comparisons between structural distance (left) and sequence distance (right). The first row showed the distributions over all pairs of positions, and the next rows represented distributions over DVG junctions with different cutoff values for counts (2, 5, 10). As the cutoff value increased, a greater proportion of distances are under 100, and the mean values get smaller.

**Figure S3.**
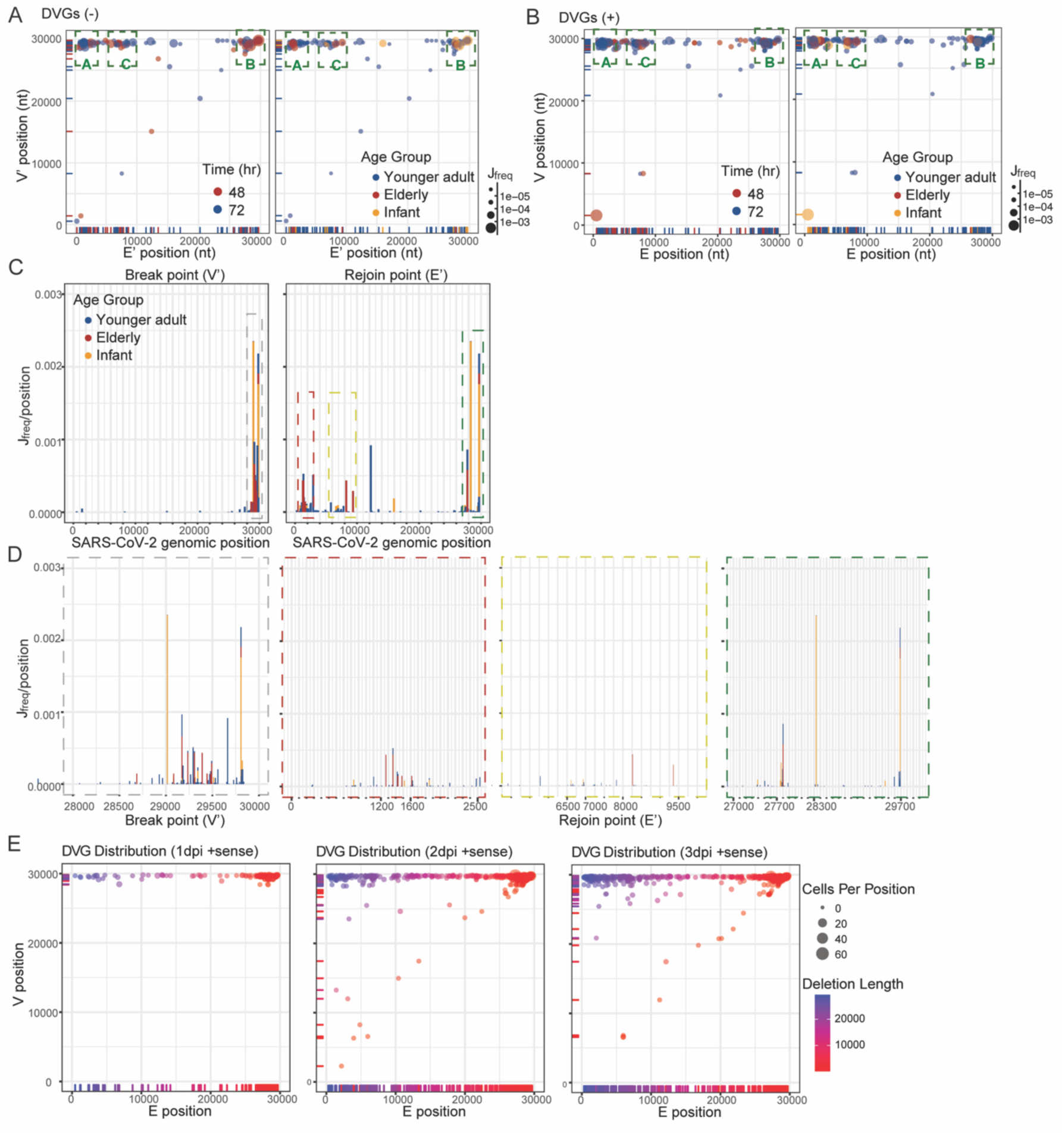
Junction distribution of DVGs identified in bulk RNA-seq and scRNA-seq using infected NHBE cells. **(A-D)** graphed DVGs of NGS used in Fig. 3. Junction distributions for identified −sense **(A)** and +sense **(B)** DVGs from infected NHBE cells of different age groups were graphed as scatterplot. Circle color represented harvest time post infection or patient age group. **(C)** The location distribution of Break point and Rejoin point of −sense DVGs were plotted separately as bar graph. The dashed boxes indicated hotspots with high concentrations of break or rejoin points. The width of each bar represented 300 nucleotides. **(D)** Detailed positions of identified hotspots clustered with −sense DVG break and rejoin points. The color of the dashed outline around each graph indicated the corresponding hotspot with the same color in (C). The width of each bar represented 10 nucleotides. **(E)** represented scRNA-seq used in Fig.4 and 5. Break point (E) and rejoin point (V) distributions of +sense DVGs were graphed at different time points post infection. Circle size represented cell count per position and circle color represents length of deletions in DVGs.

**Figure S4.**
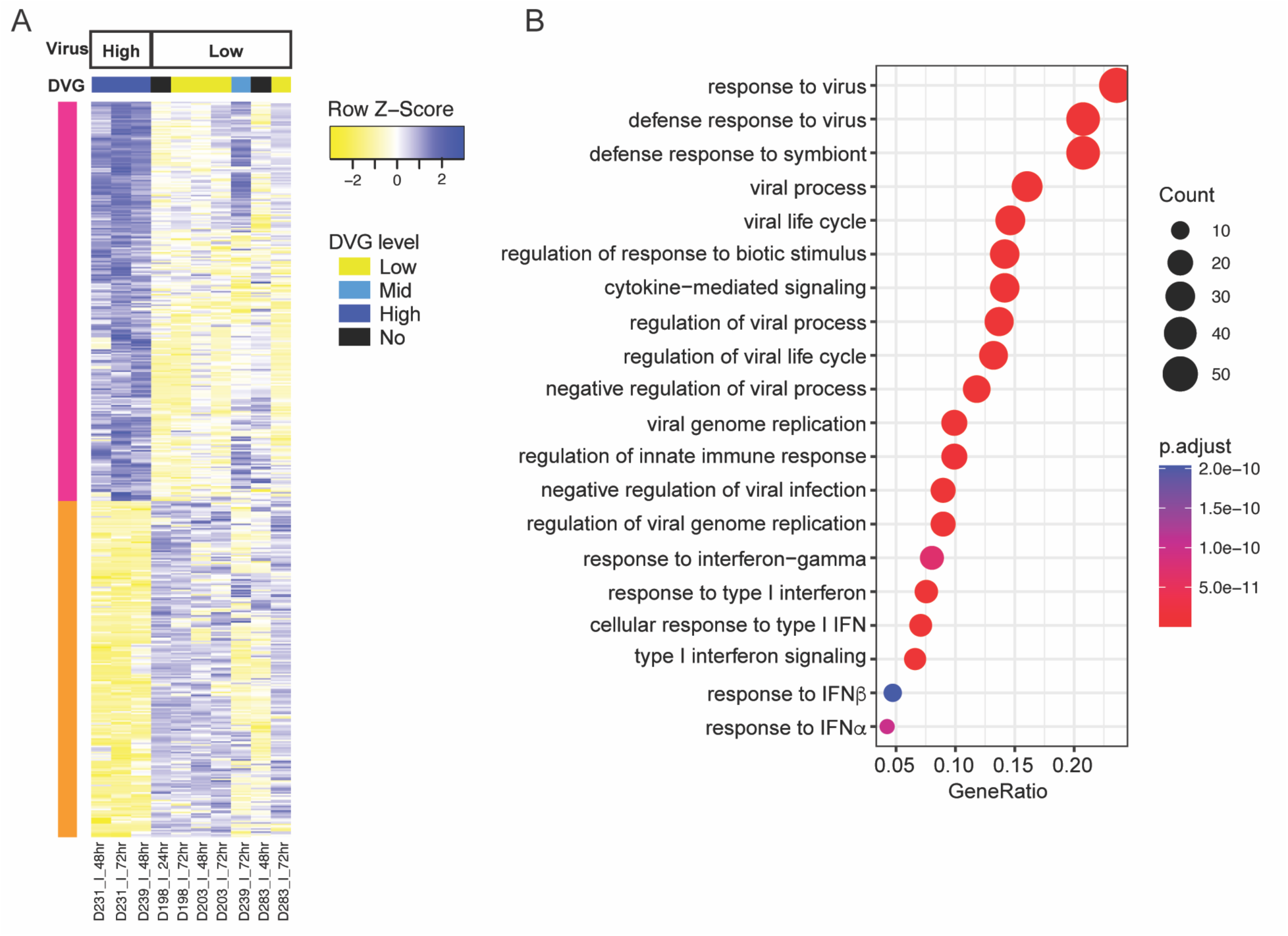
Antiviral (type I IFN) responses was upregulated in samples with high viral counts and DVG counts. **(A)** Differential expressed genes between high virus and low virus groups were graphed as heatmap for all samples. Pink cluster was the genes upregulated in high virus group and orange cluster was the gens downregulated in high virus group. The virus group and the DVG level of each sample were both indicated on top of the heatmap. **(B)** Gene ontology analysis of genes that were upregulated in high virus group with high DVG level (pink cluster) were graphed in R (GOplot). Circle size represented number of genes in each pathway. Gene ratio represented the ratio of number of genes in that pathway to the number of genes in the entire cluster.

**Figure S5.**
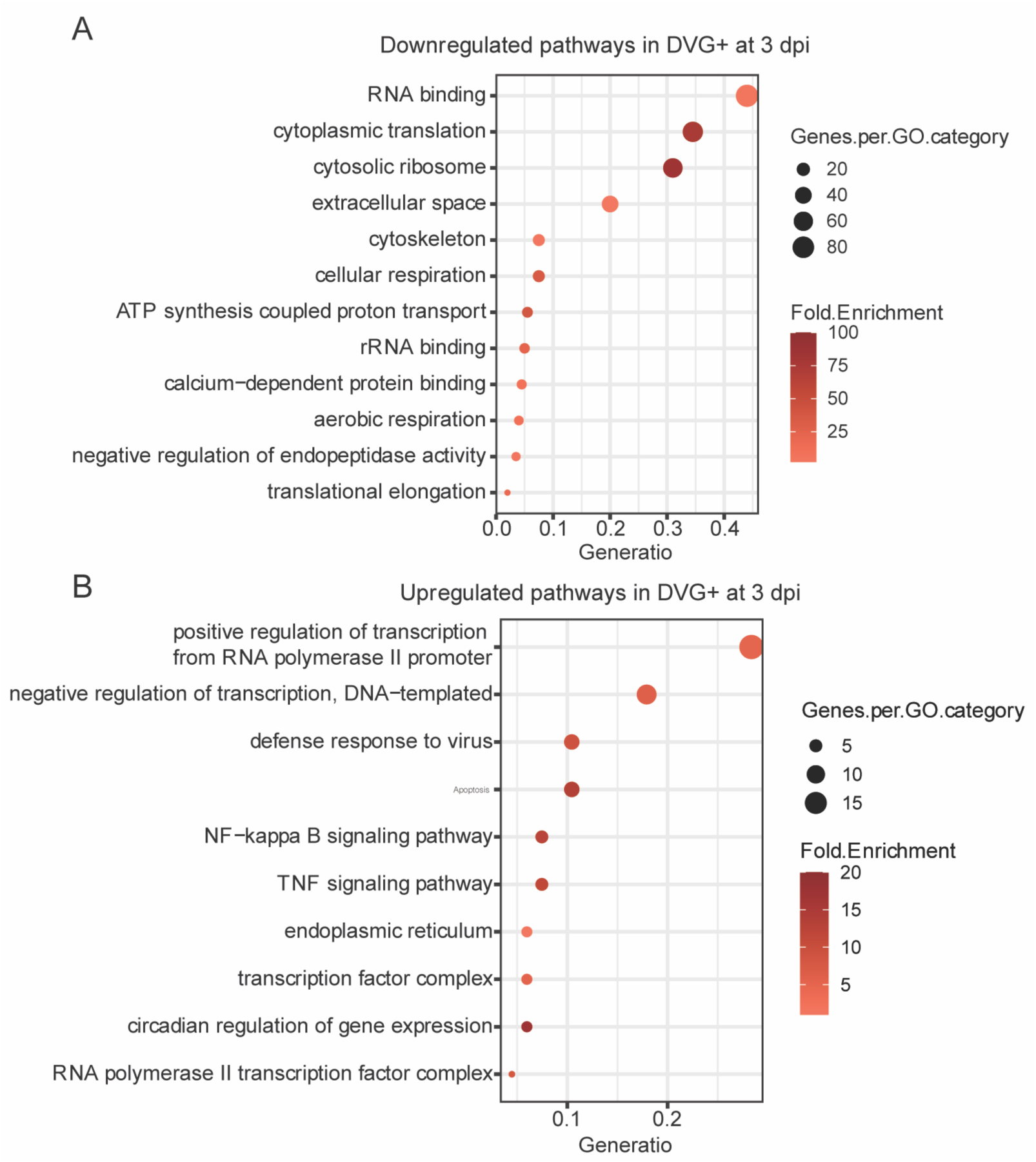
Gene ontology analysis of differential expressed genes between DVG+ and DVG− groups at 3 dpi. Gene ontology analysis of genes that were downregulated **(A)** and upregulated **(B)** in DVG+ cells relative to DVG− cells at 3 dpi. Circle size represented number of genes in each pathway. Gene ratio represented the ratio of number of genes in that pathway to the number of genes in the entire cluster.

**Figure S6.** ViReMa, Cellranger and Seurat Pipelines

### Introduction

The purpose of this standard operating procedure is to outline the pipeline used for ViReMa. This document describes the steps needed to identify and analyze defective viral genomes (DVGs) from bulk RNA-Seq and single cell RNA-Seq (scRNA-Seq) data in SARS-CoV-2.

The Bowtie2 and ViReMa scripts were both run on the BlueHive Linux computing cluster supported by the Center for Integrated Research Computing at the University of Rochester.

We used version 2.2.9 of Bowtie2 to map our samples to the human genome. We used the GRCh38 (hg38) human reference genome. We also used UMI-Tools version b1 for our single cell RNA-Seq analysis.

We used version 0.21 of ViReMa to identify the DVG recombinant events and their corresponding counts. Version 0.21 of ViReMa uses version 0.12.9 of Bowtie and Python3 to map each sample to the reference viral genome. We used the SARS-CoV-2 reference genome with GenBank ID MT020881.1.

For the rest of our analysis, we used version 4.1.0 of R and version 1.4.1717 of RStudio. Our analysis used the following packages:

- Rsubread
- tidyverse
- ggplot2
- plotly
- openxlsx
- data.table

### DVG analysis from bulk RNA-Seq dataset

The pipeline to identify DVGs from bulk RNA-Seq analysis was as follows:

1. Bowtie2
2. ViReMa
3. Subread
4. R filtering

#### Bowtie2

We used Bowtie2 to align our sample to the human reference genome (GRCh38 (hg38)). The GRCh38 (hg38) index was downloaded from the Bowtie2 website. The unmapped output sequence served as the viral sequence to be used for ViReMa.

The SLURM script used to run Bowtie2 alignment for single end reads is shown below:

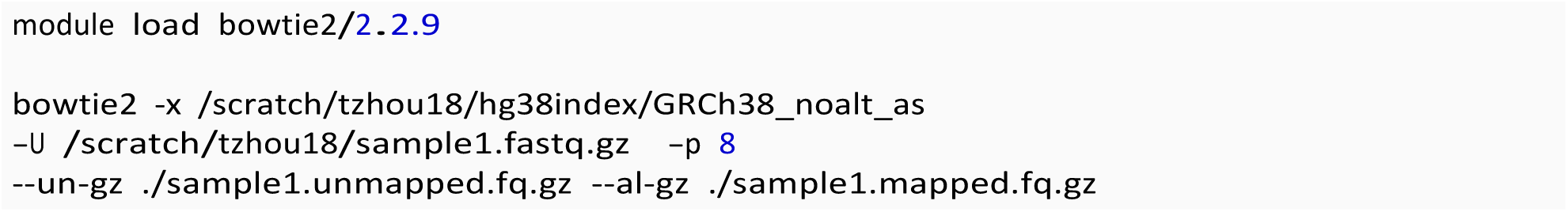

For paired end samples, properly paired read files were specified using the -1 and -2 options instead of the -U option used for single end reads.

#### ViReMa

We used ViReMa to identify viral recombinant events.

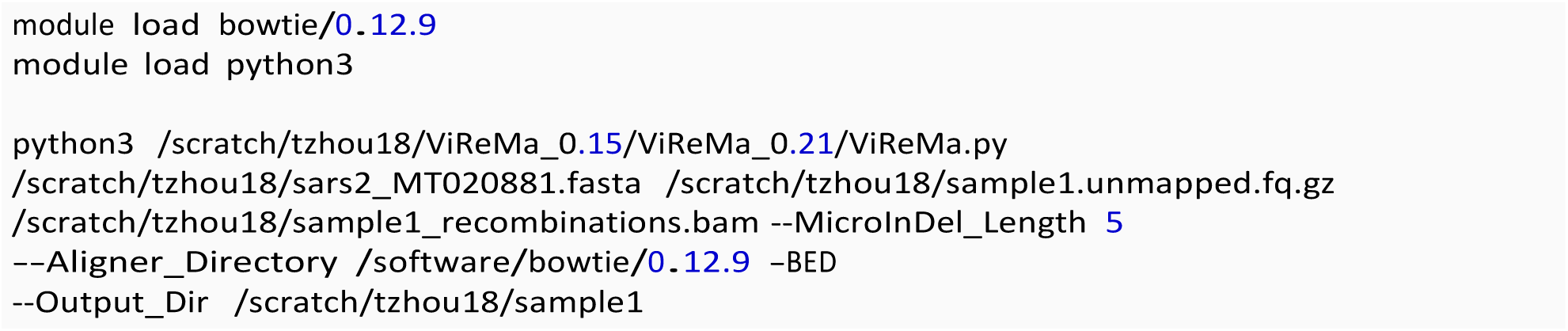

The Virus_Recombination_Results.bed file within the BED_Files folder and the recombinations.bam file were used for the downstream analysis.

#### Subread

Bioconductor R package Rsubread (v2.6.4) was used to align our RNA-seq data to the viral reference genome to identify the number of viral reads in each sample.

To import each sample into RStudio to run Subread, a tab-delimited file named study_design.txt was created to contain the file names and paths, as shown below:

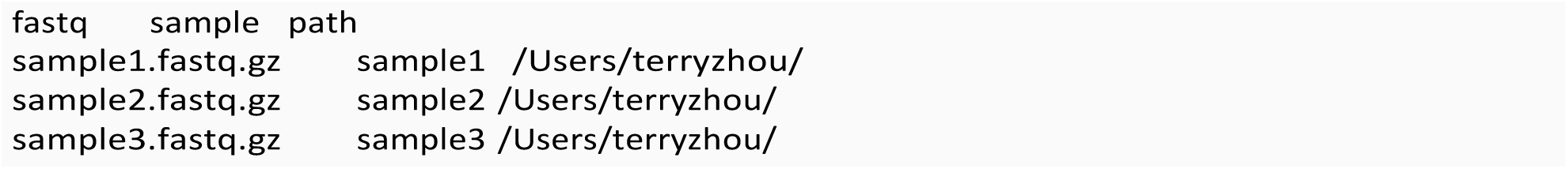

The following R script was used to run Rsubread in RStudio.

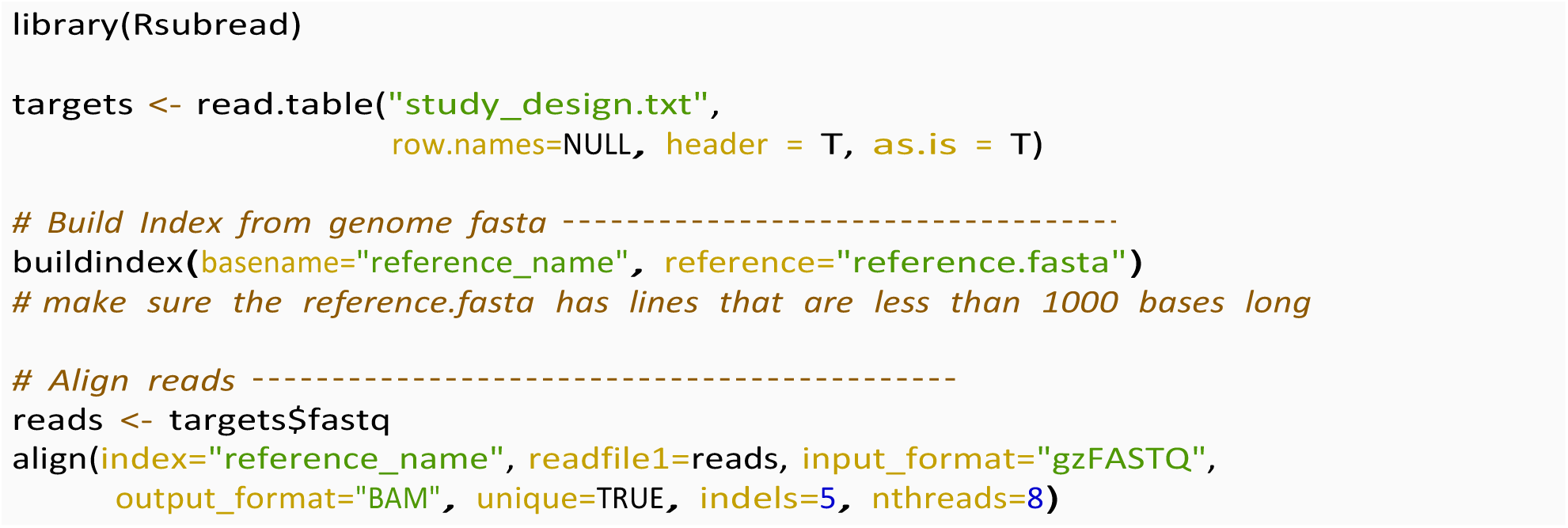

The number of viral reads printed out in the R console, as well as the subread.BAM.summary file, in the Uniquely_mapped_reads row were used as the total counts of viral reads (Fig1B_virus).

#### R Filtering Script

The following R script was used to filter out recombinations that are not deletions (i.e. insertions, duplications), deletions shorter than 100 nt, and those that had a break point before the 85 nt position. We also separated the identified DVGs into positive and negative sense and analyzed them separately.

Since viral load can affect DVG level, we used the junction frequency (*Jfreq*) as a standardized value to quantify DVG level. We calculated *Jfreq* by dividing the DVG count by the viral read count. The viral read count was identified via the previous Subread step.

The filtering script is shown below:

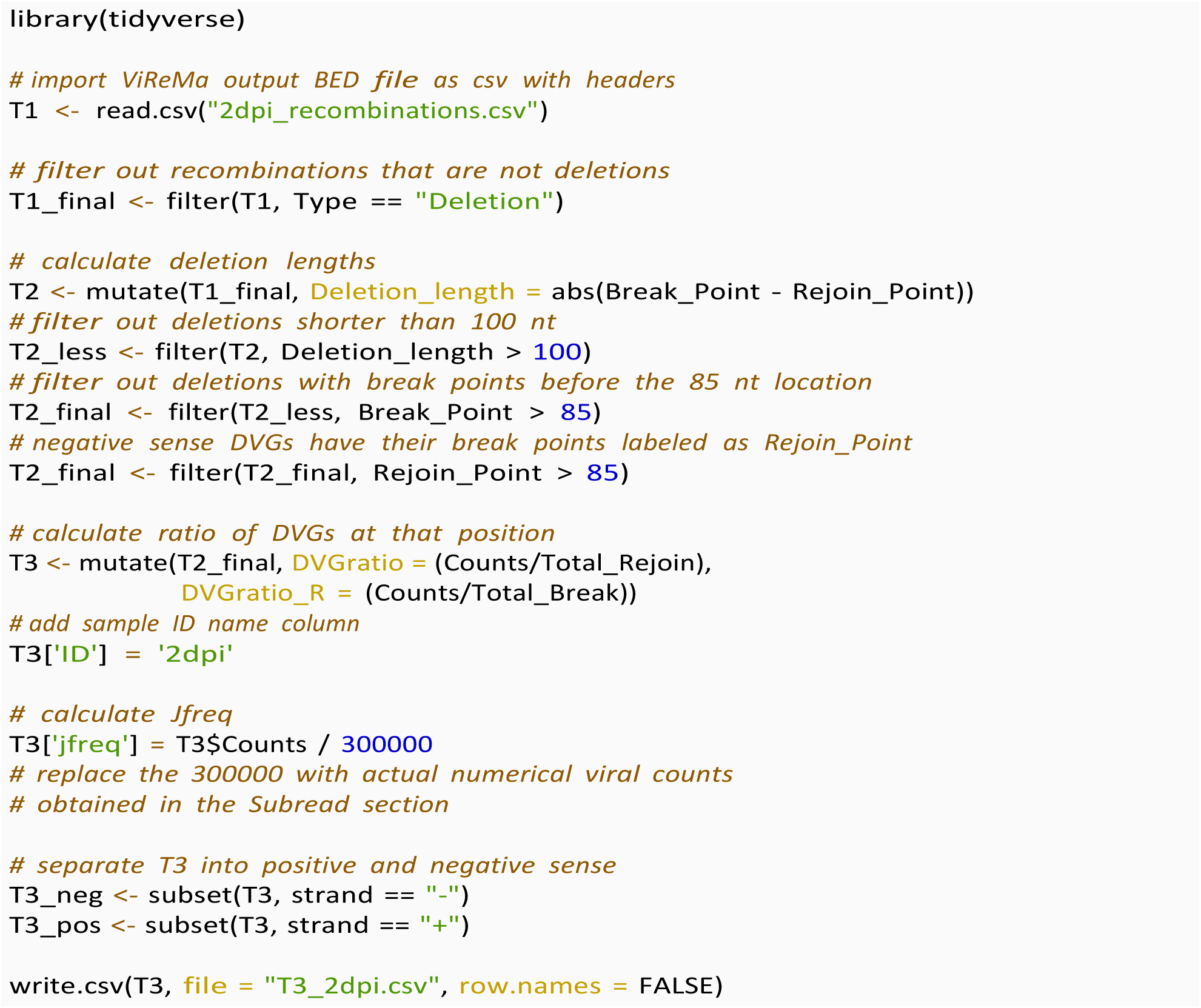

#### Making Plots

We used the following script to graph plots as shown in Fig. 2 and Fig. S1.

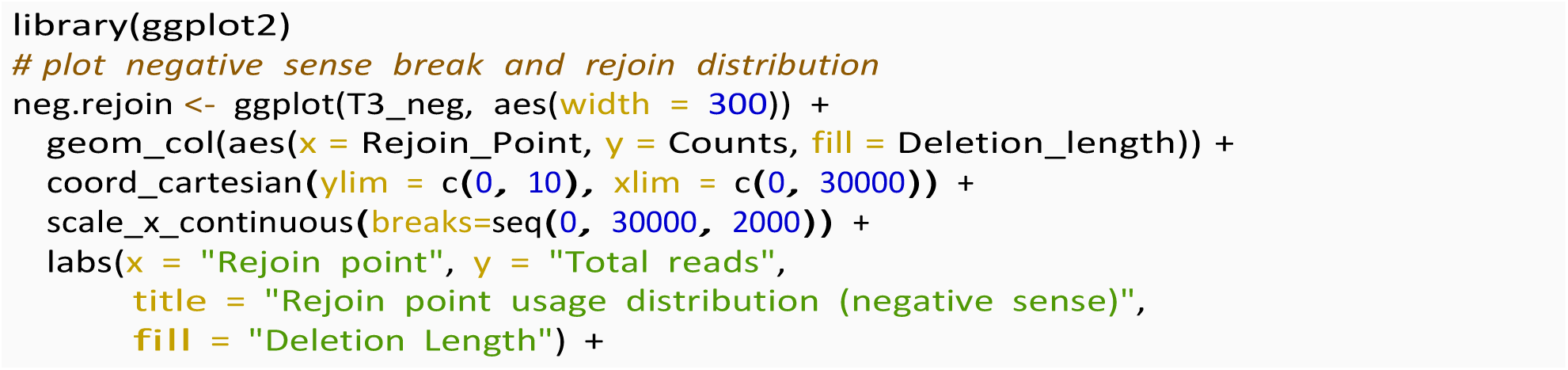
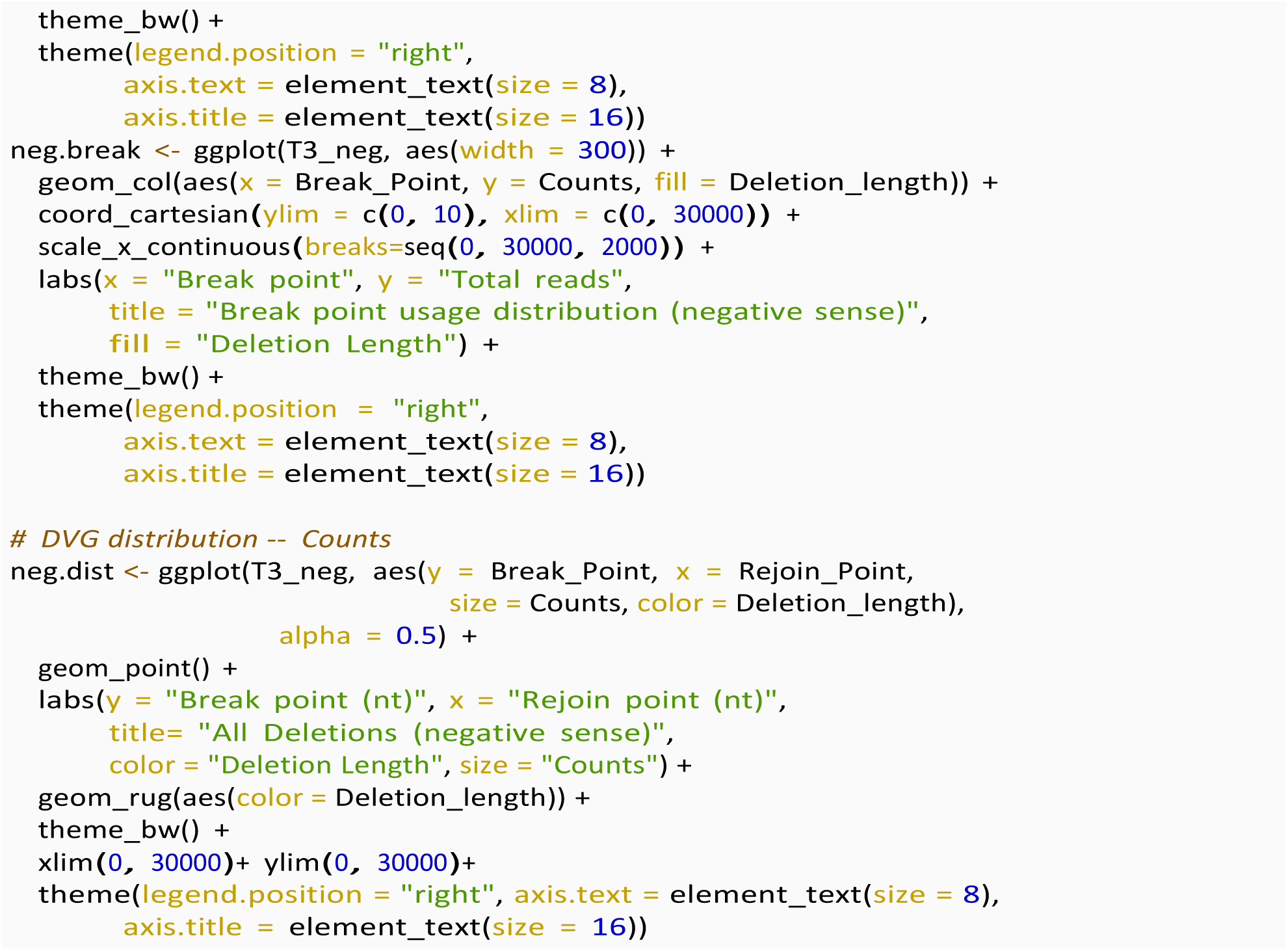

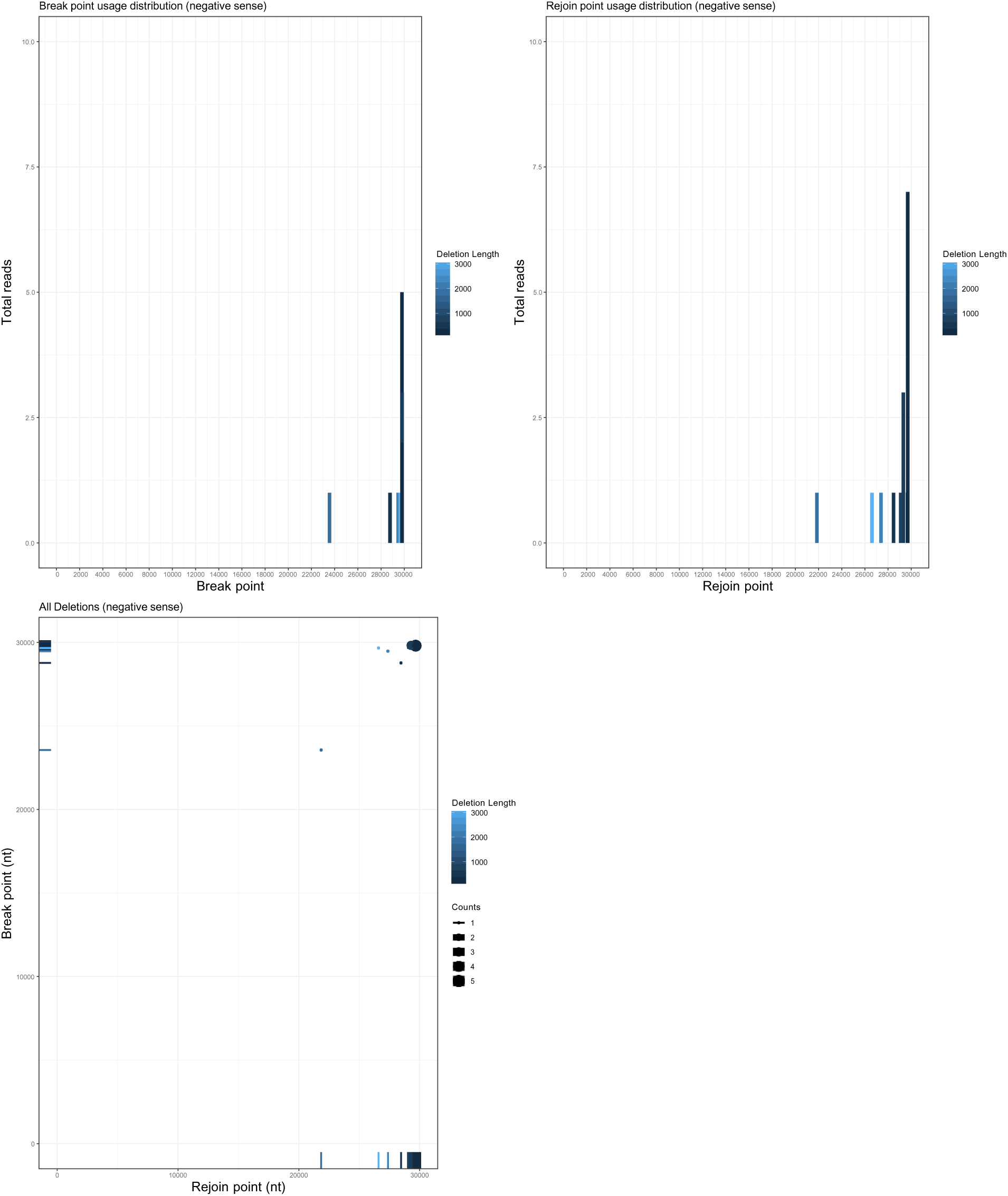

### DVG analysis from scRNA-Seq dataset

The pipeline to identify DVGs from scRNA-Seq analysis was as follows:

1. UMI-Tools
2. Bowtie2
3. ViReMa
4. R filtering

#### UMI-Tools

UMI-Tools was used to associate the cell barcodes and UMIs for each read to the sequence. The cell barcodes and UMIs from the R1 file were combined with the corresponding read in the R2 file.

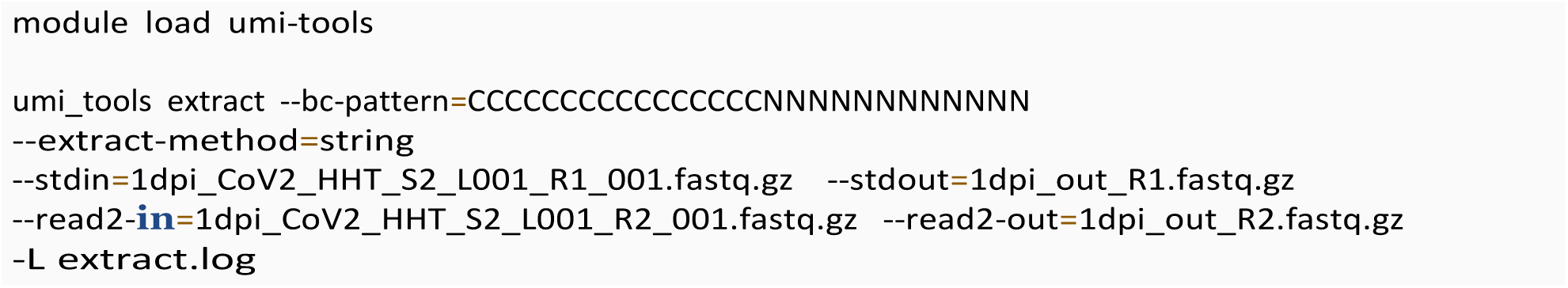

#### Bowtie2

As with the bulk RNA-Seq analysis, we are used Bowtie2 to align our sample to the human genome. The unmapped output file was used for ViReMa analysis.

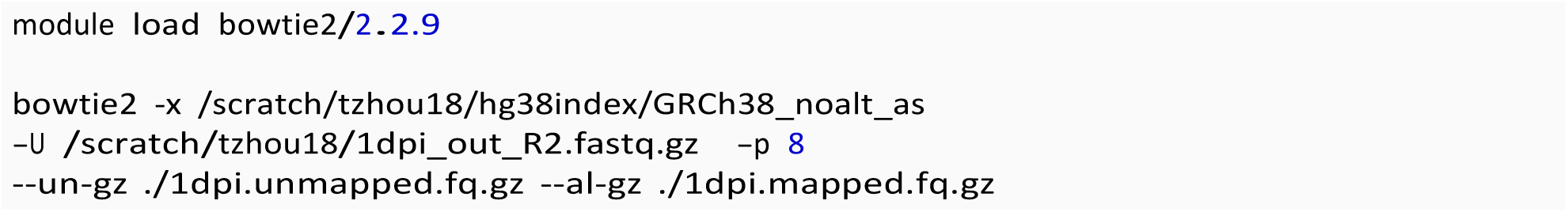

#### ViReMa

For scRNA-Seq analysis, ViReMa must be run twice in order to link the cell barcodes and UMIs to the identified DVGs. The first run was identical to running ViReMa for bulk RNA-Seq. The second run was as follows:

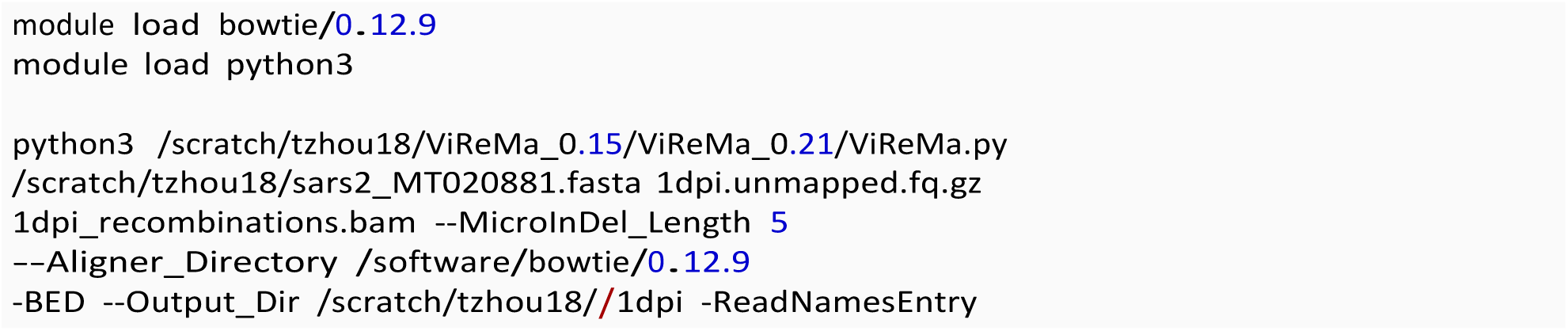

The Virus_Recombination_Results.bed file within the BED_Files folder and the recombinations.bam file from the first run and the Virus_Recombination_Results.txt file from the second run were used for the following R filtering.

#### R Filtering Script

The first section of the R filtering script was identical to bulk RNA-seq section.

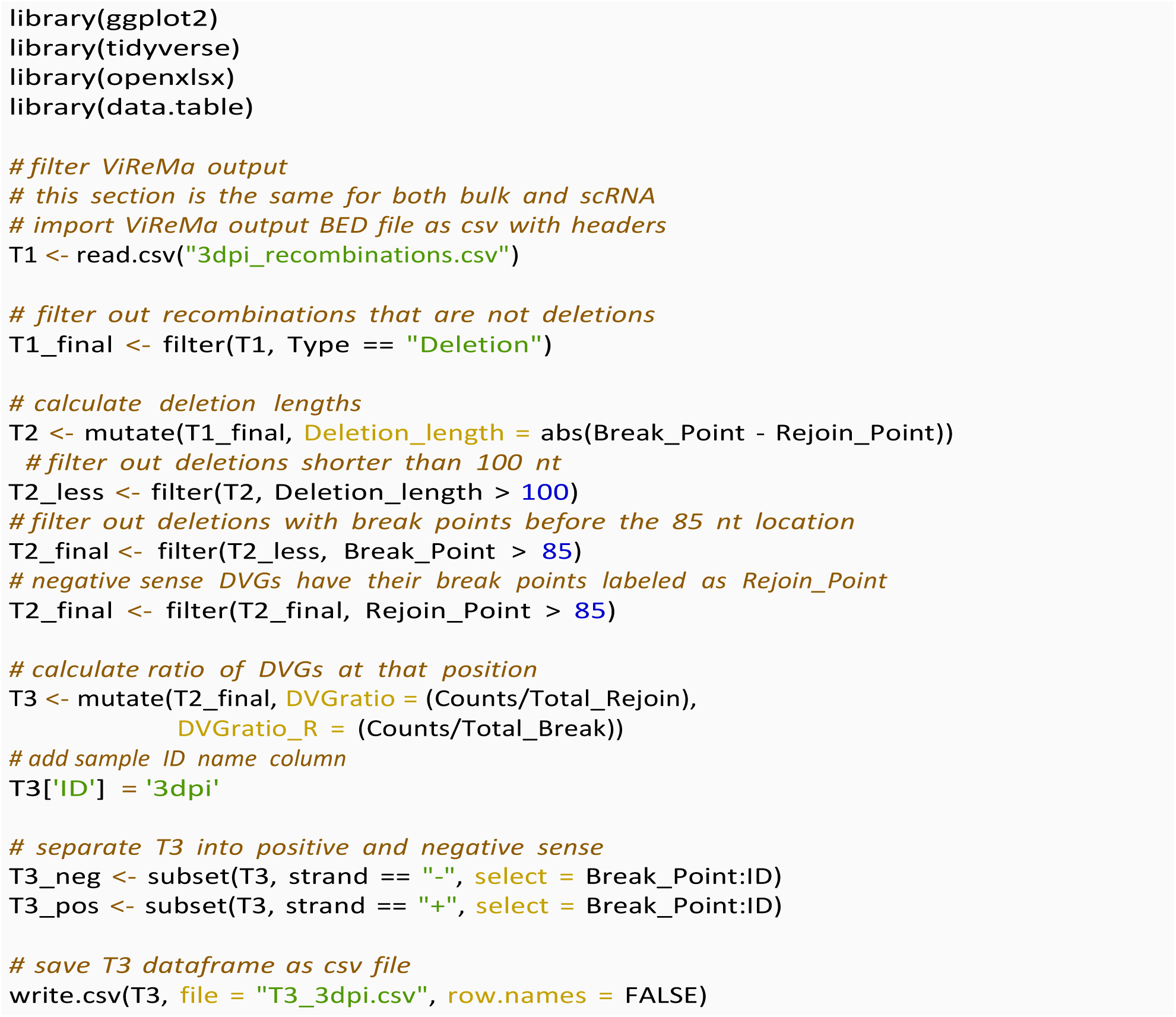

This next section of the R filtering was unique to scRNA-seq analysis. The input file for this section was the Recombination_Results.txt file from the second run of ViReMa, including all of the read counts, positions, and cell barcodes/UMIs for each DVG.

The viral.reads dataframe, including the number of viral UMIs for each cell and lists the cells in the order, was used for downstream Seurat analysis. This dataframe was retrieved by outputting only the viral reads row from the Cell Ranger gene matrix into a .csv file.

The final dataframes of interest from the following R filtering script were the df1, df2, and df3 dataframes, which included the number of DVG UMIs per cell, the number of DVG UMIs per position and the number of DVG UMIs per combination of cell and position, respectively. These datasets were used for downstream analyses. In addition, the bcmatrix dataframe was added to the matrix in Seurat as a “DVG gene.”

The following script only showed the filtering process for the positive sense DVGs, however the same script was used for the negative sense DVGs.

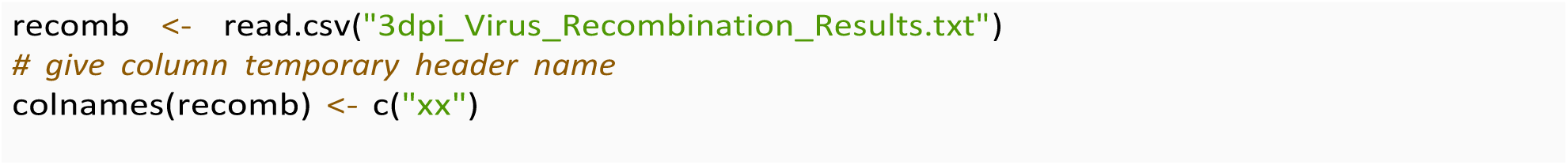
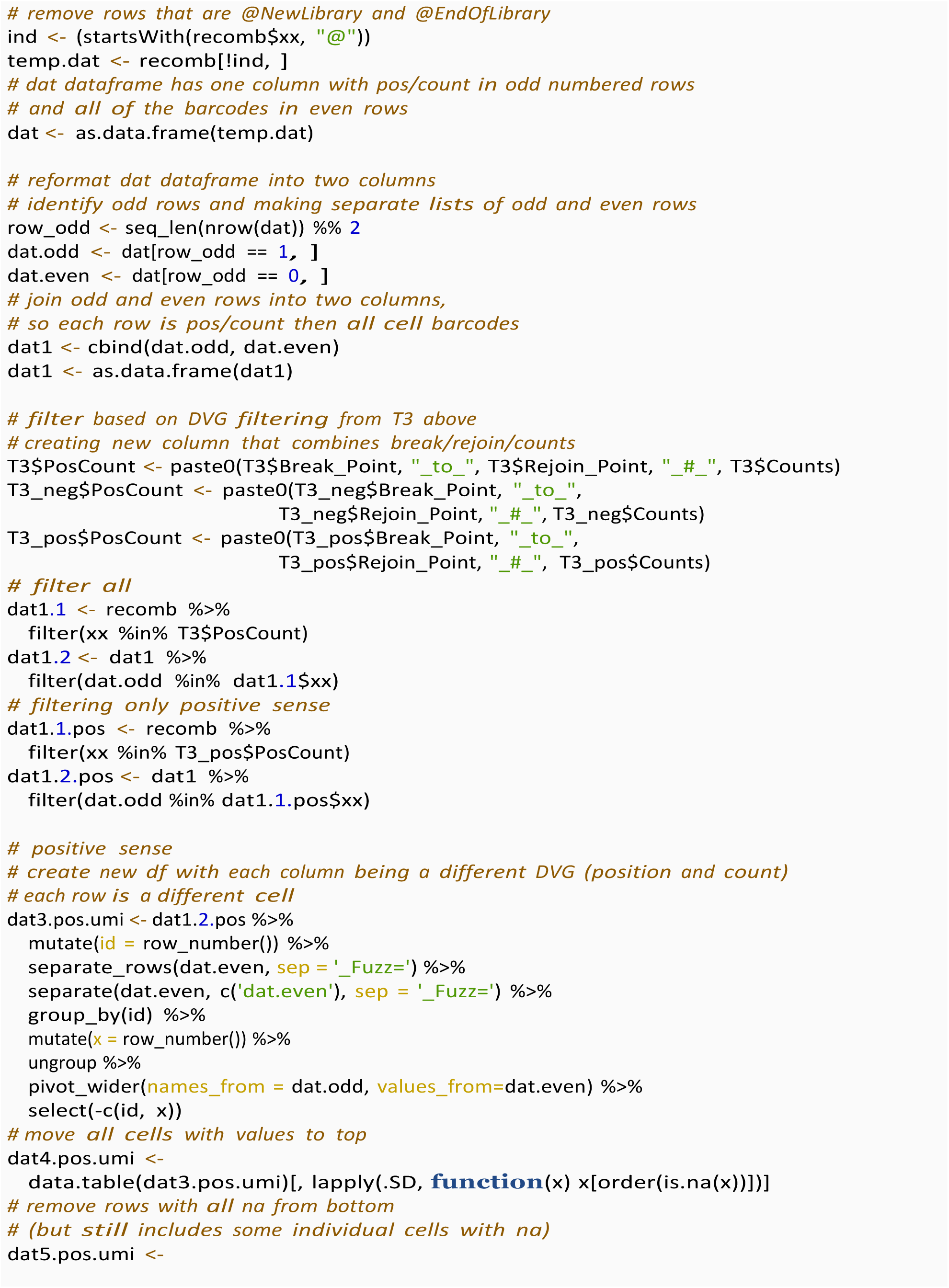
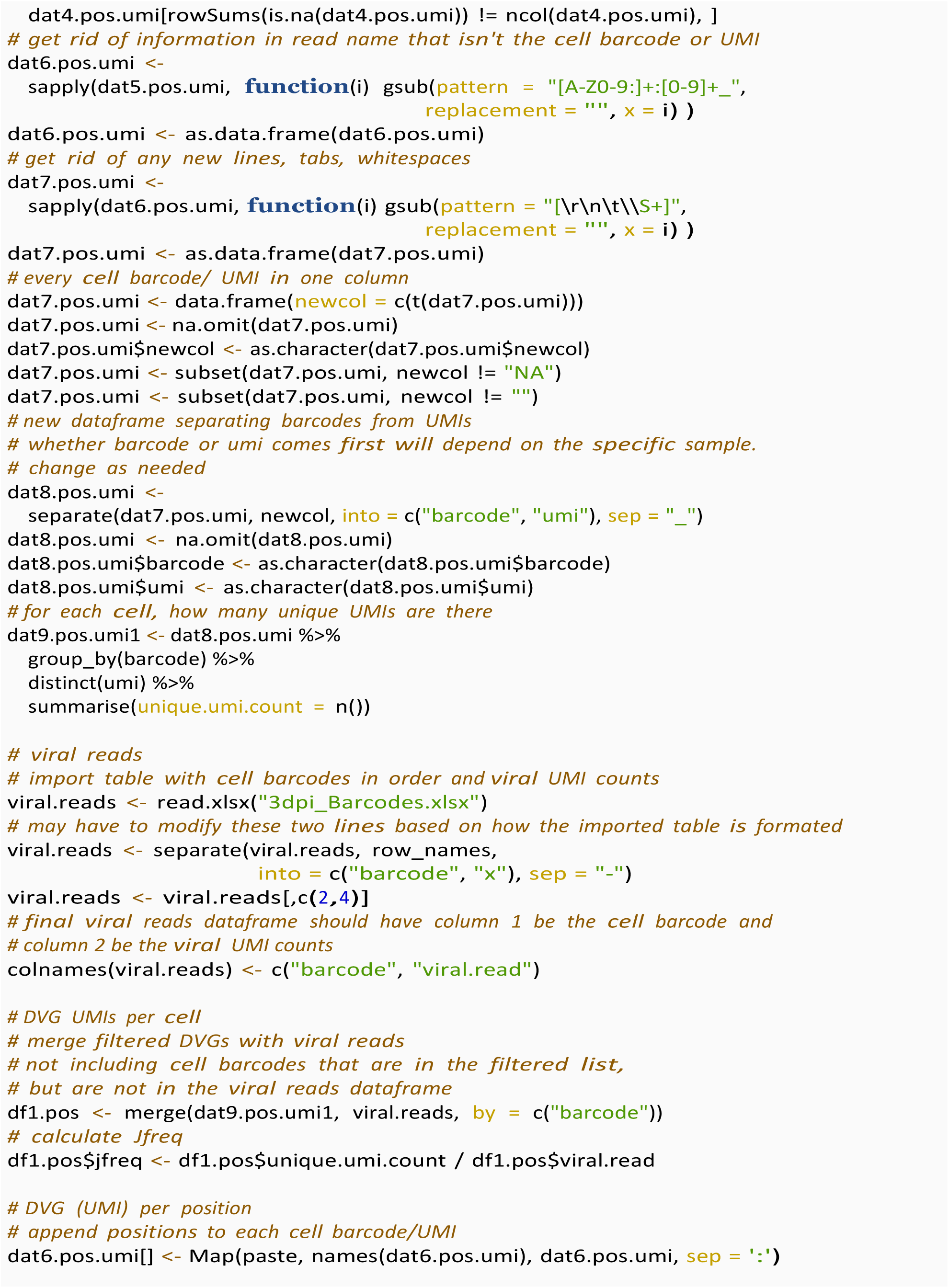
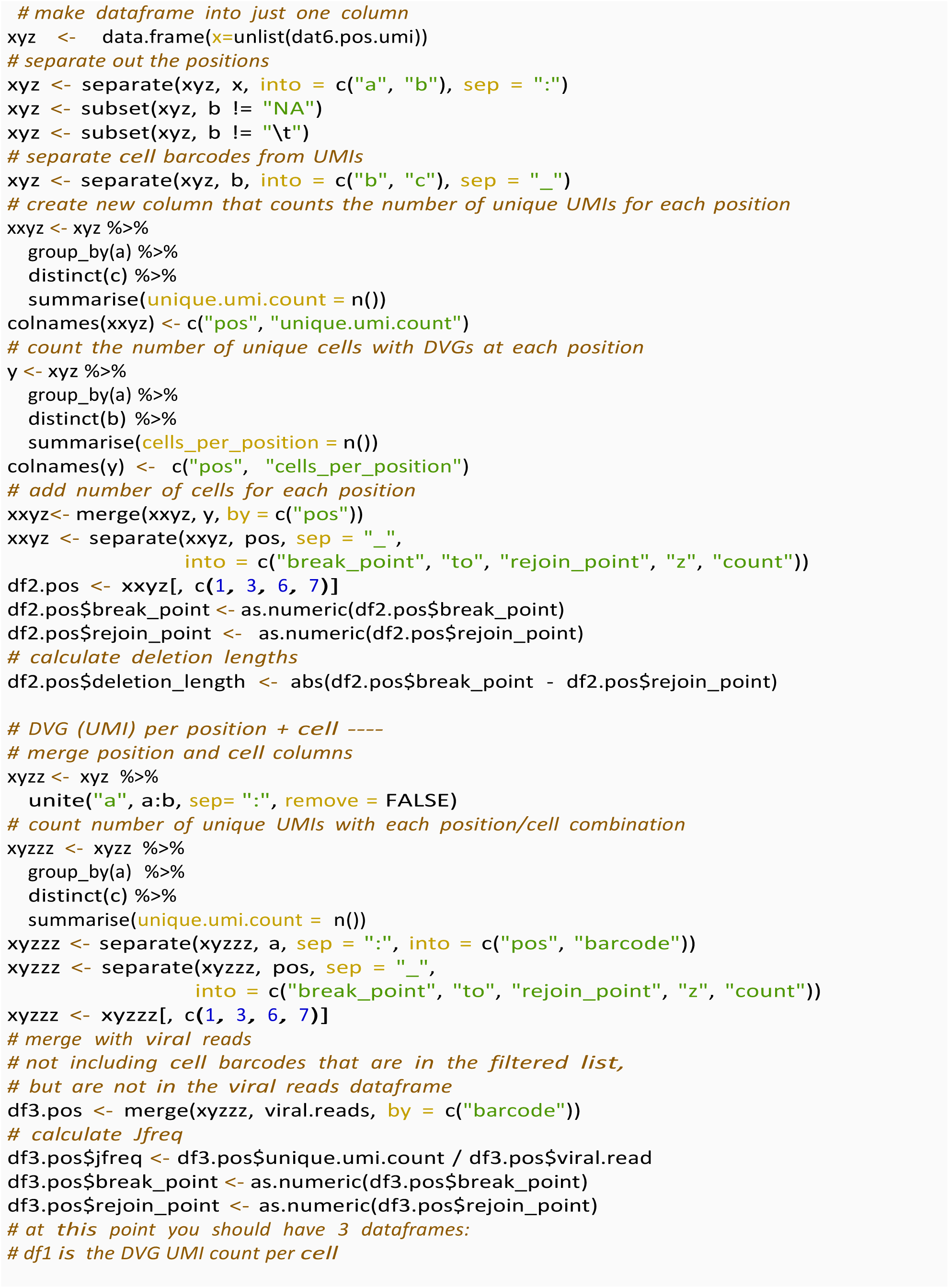
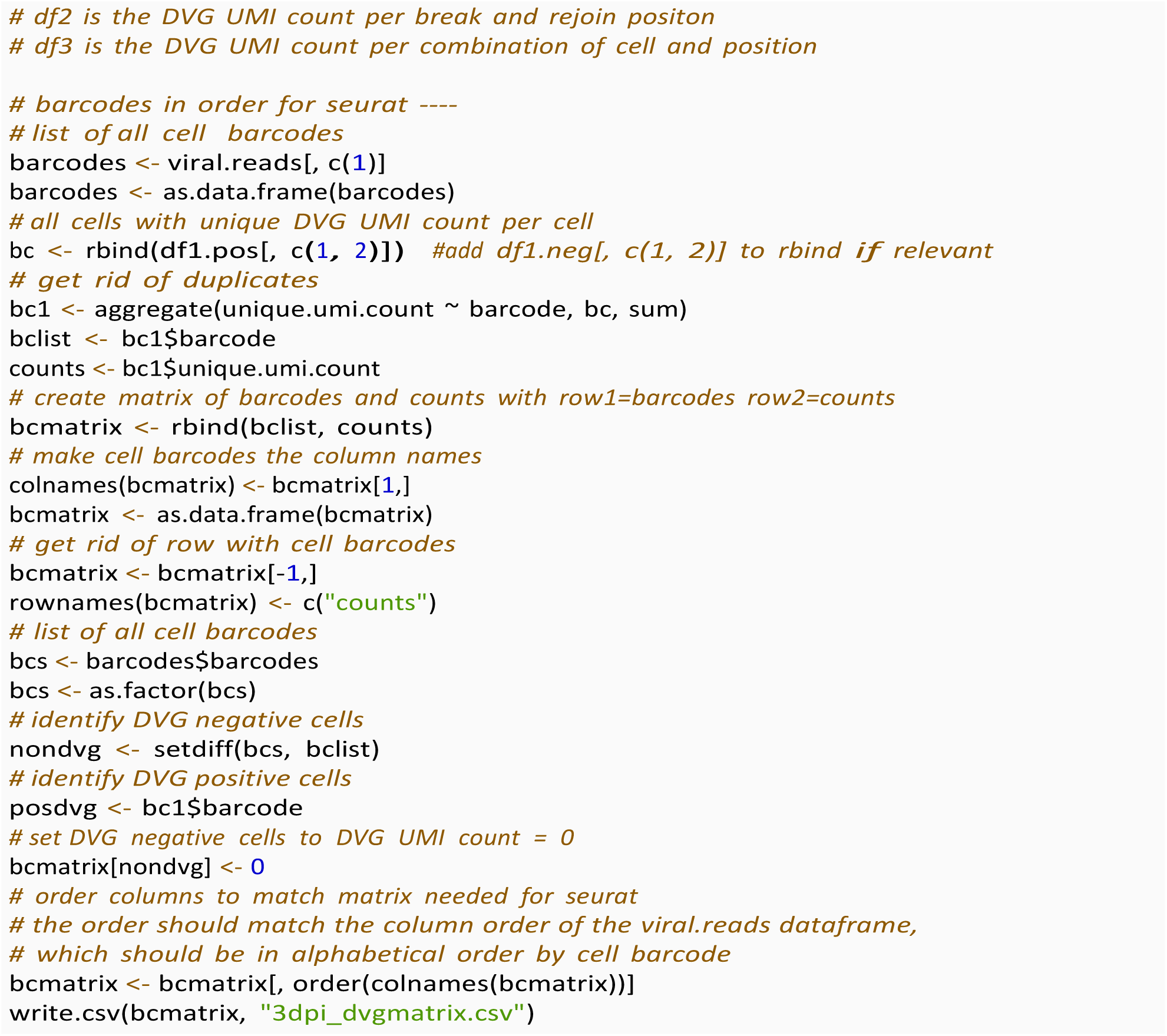

#### Making Plots

We used the following code to create exploratory data analysis plots of the positive sense DVGs, however the same code was used to visualize the negative sense DVGs.

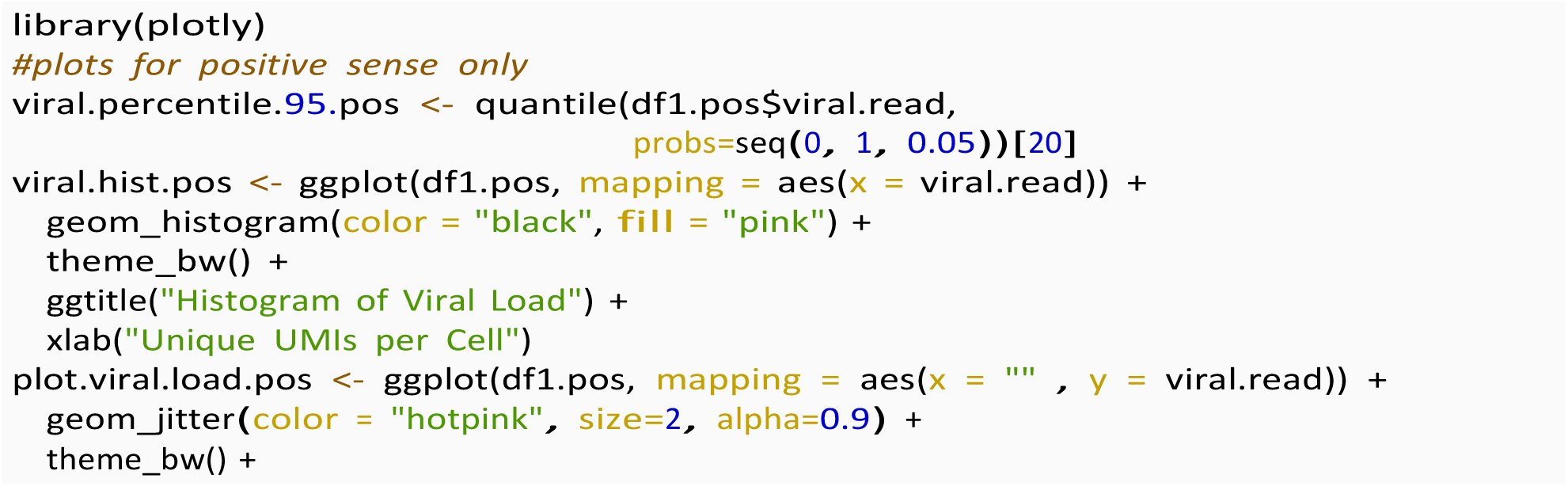
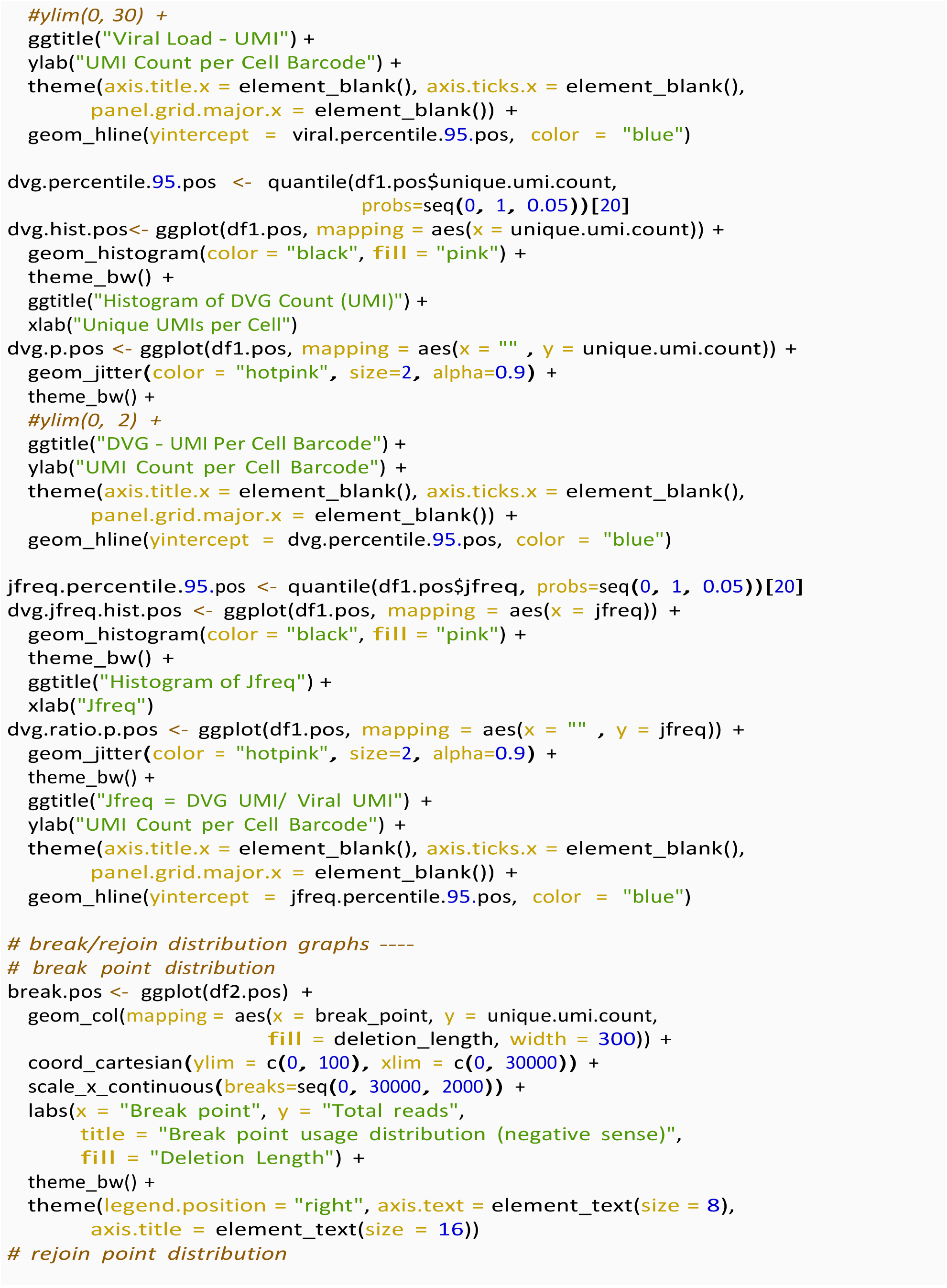
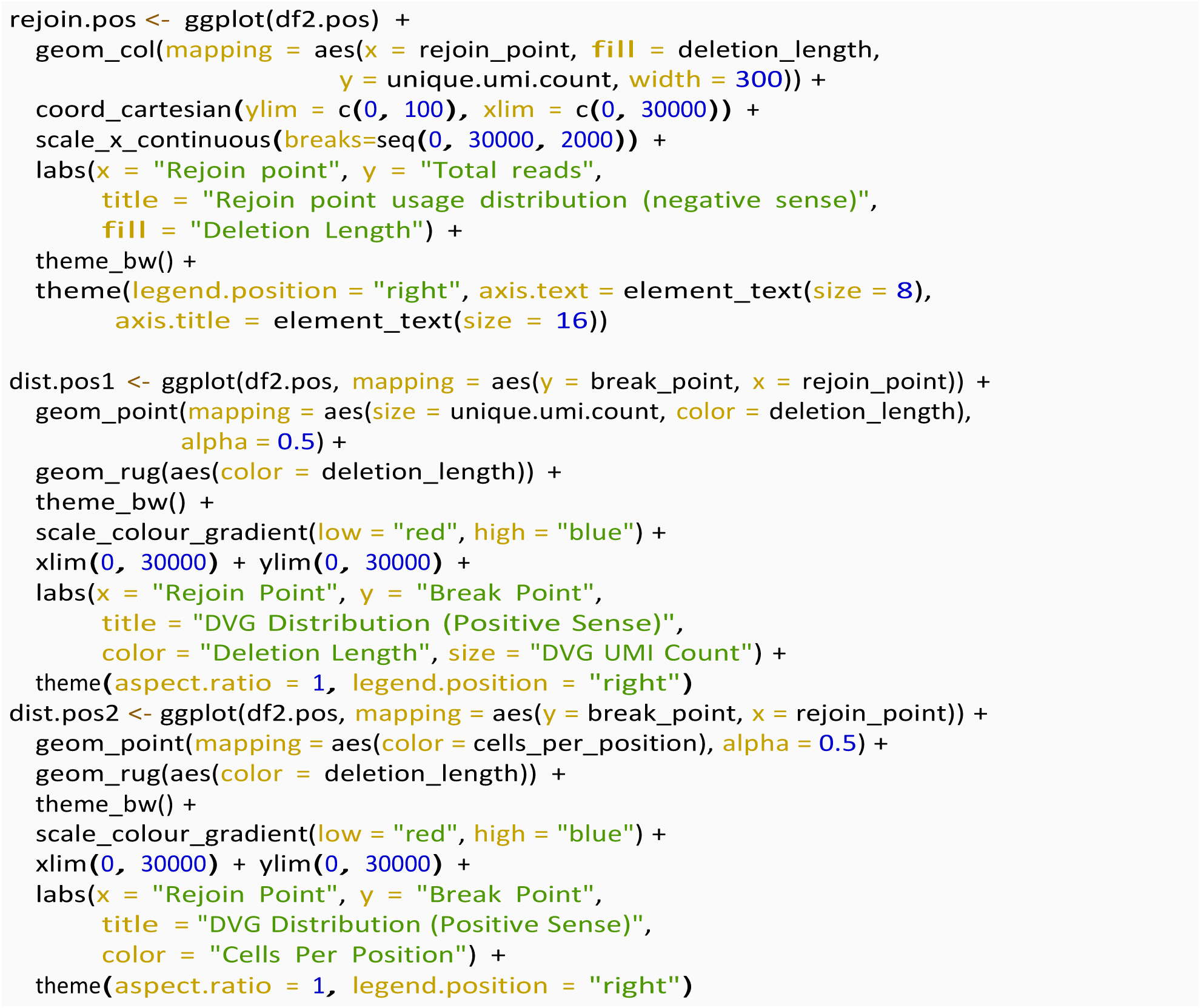

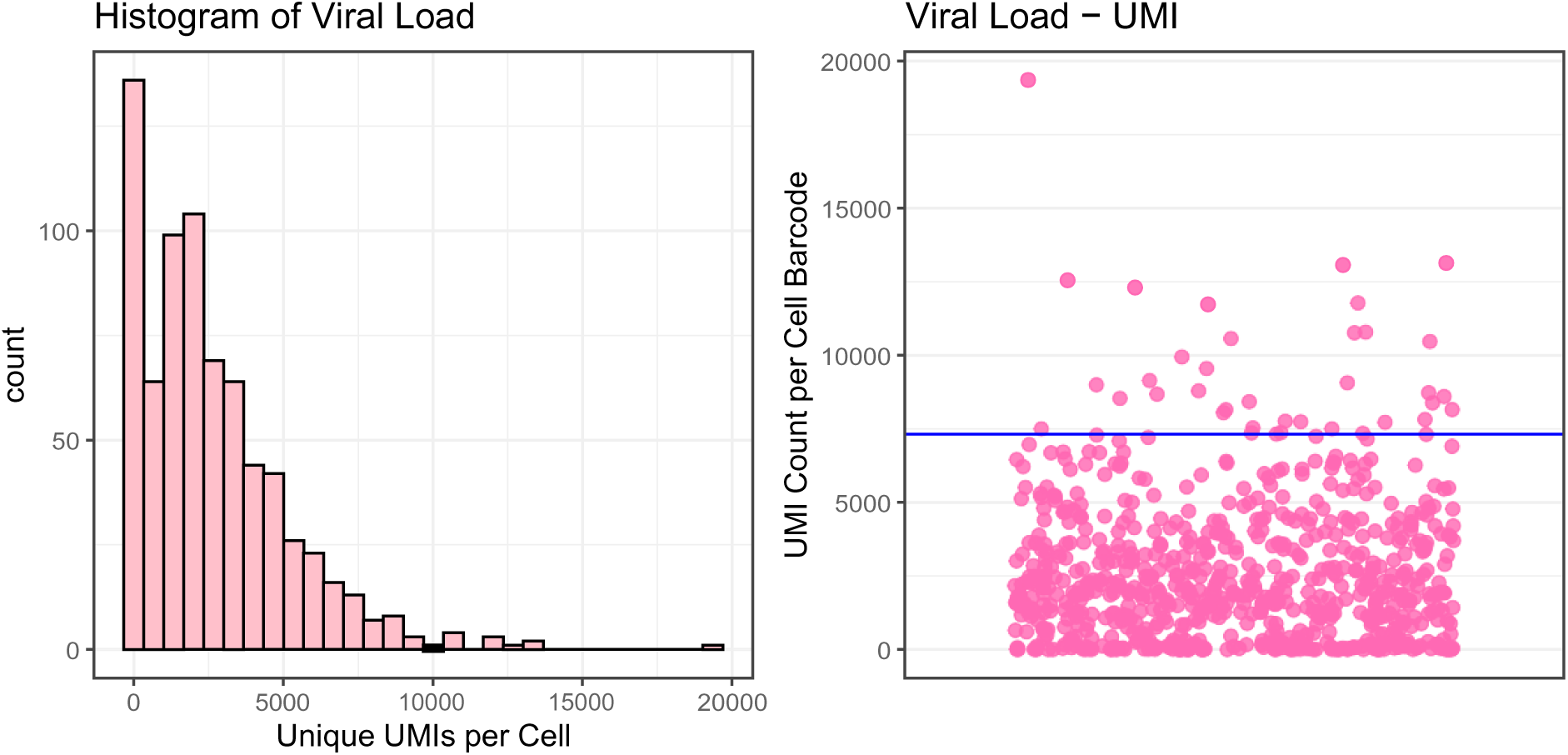
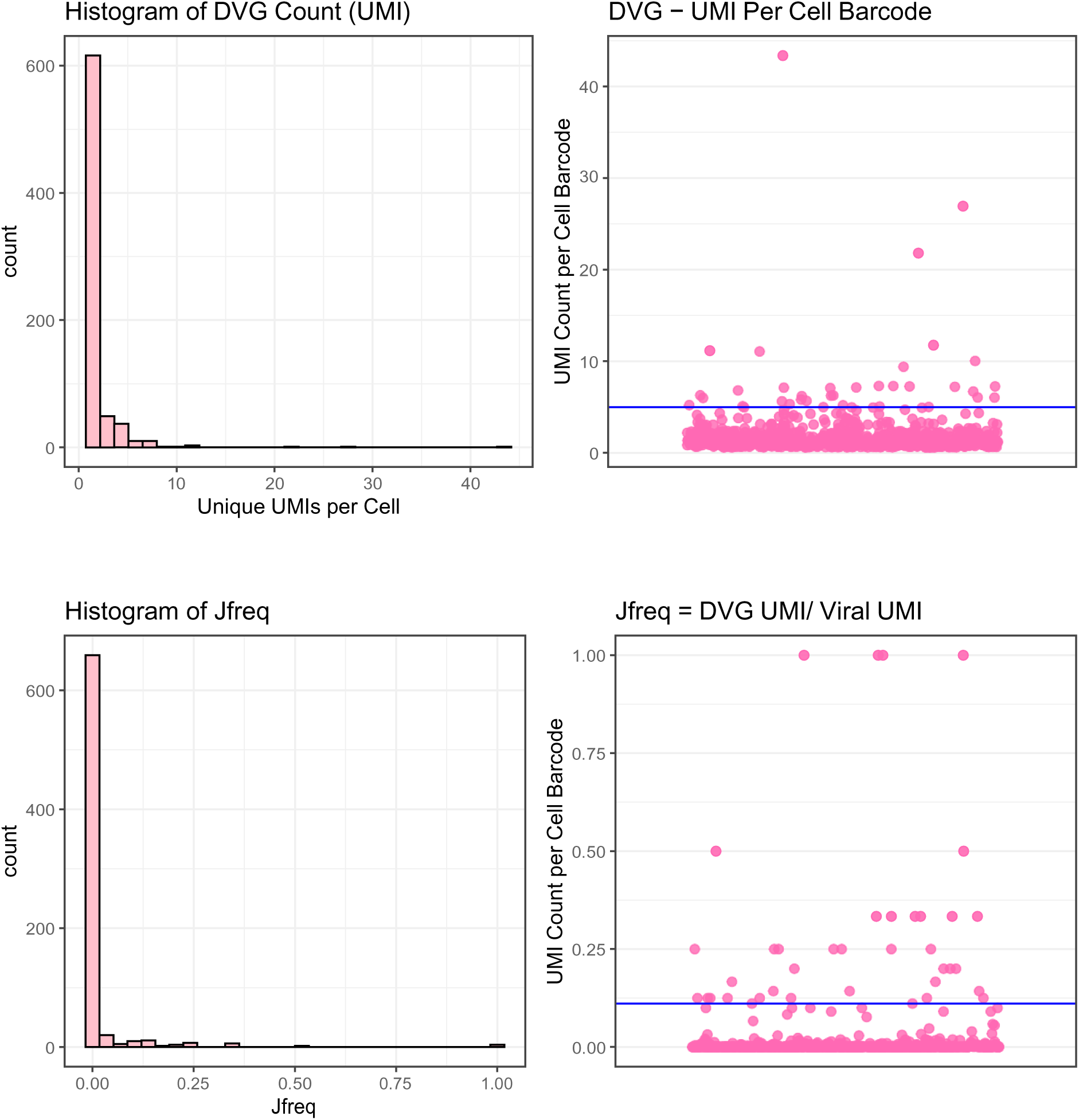
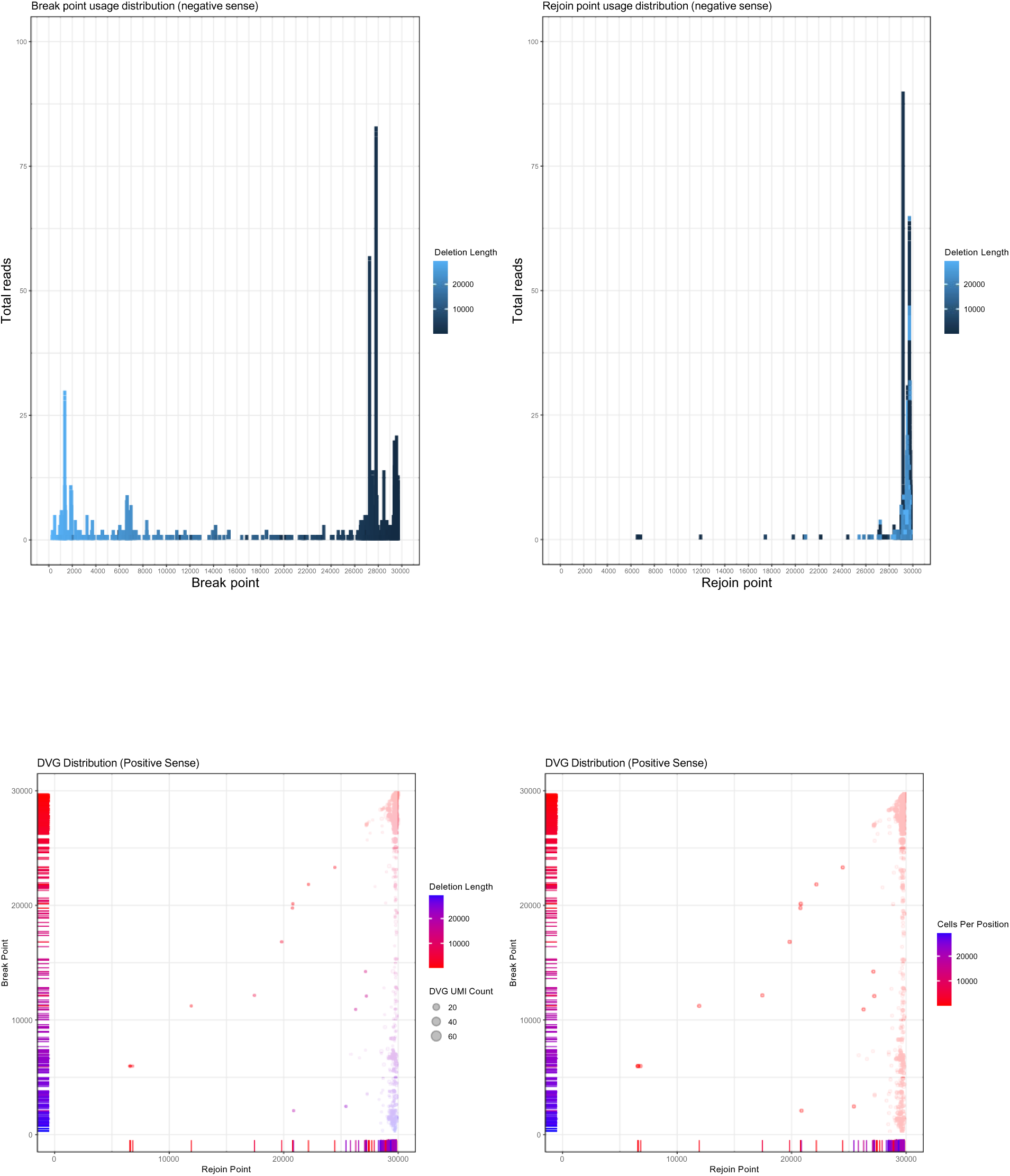

We used the following script to graph plots as shown in Fig. 4 and Fig. 5. The gene expression matrices were imported for each time point and the mock sample. The same script as Fig.4D was used to plot other time points and the same script as plots Fig.5A Fig.5B and Fig.5C was used to plot other time points and viral loads.

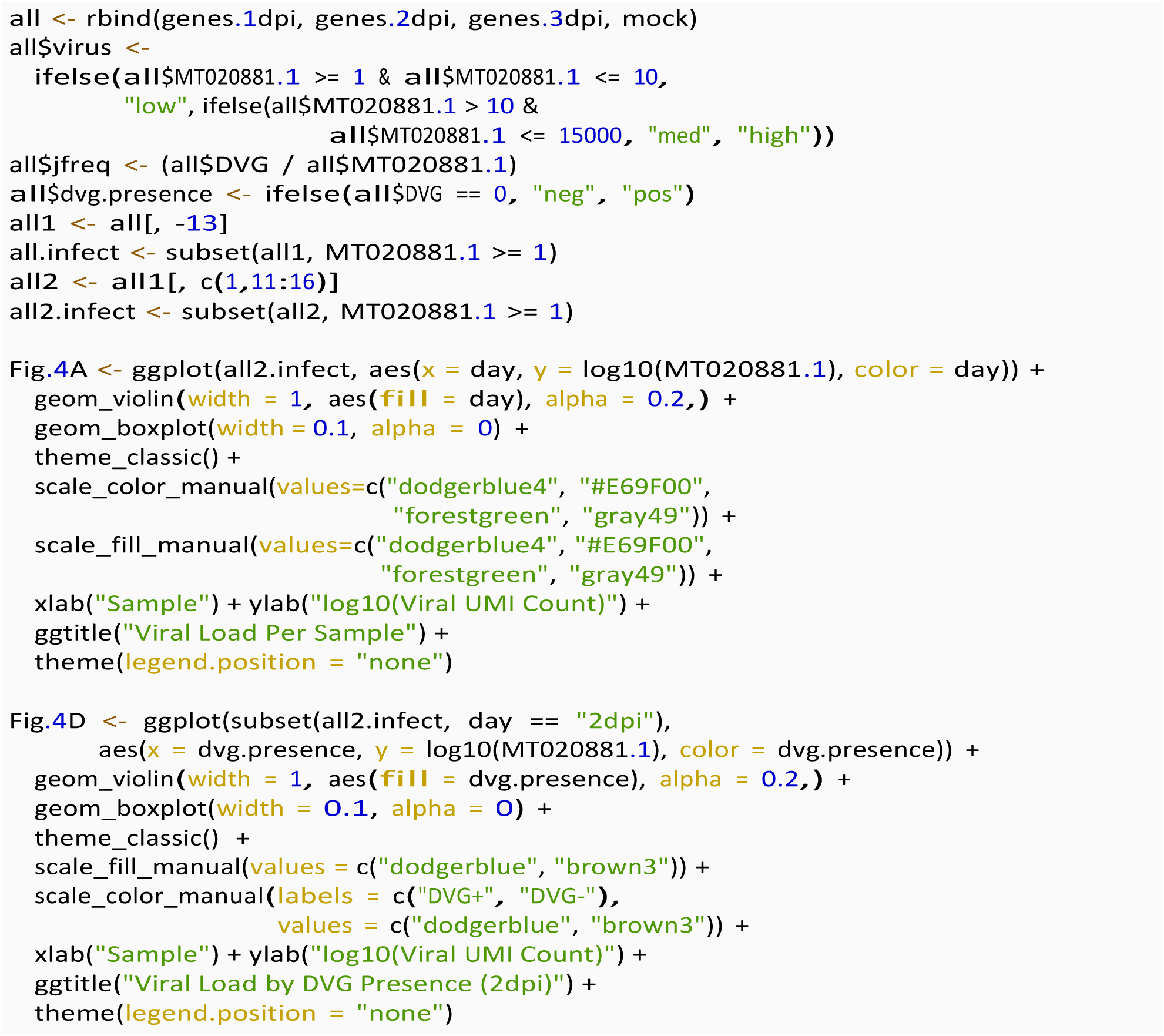

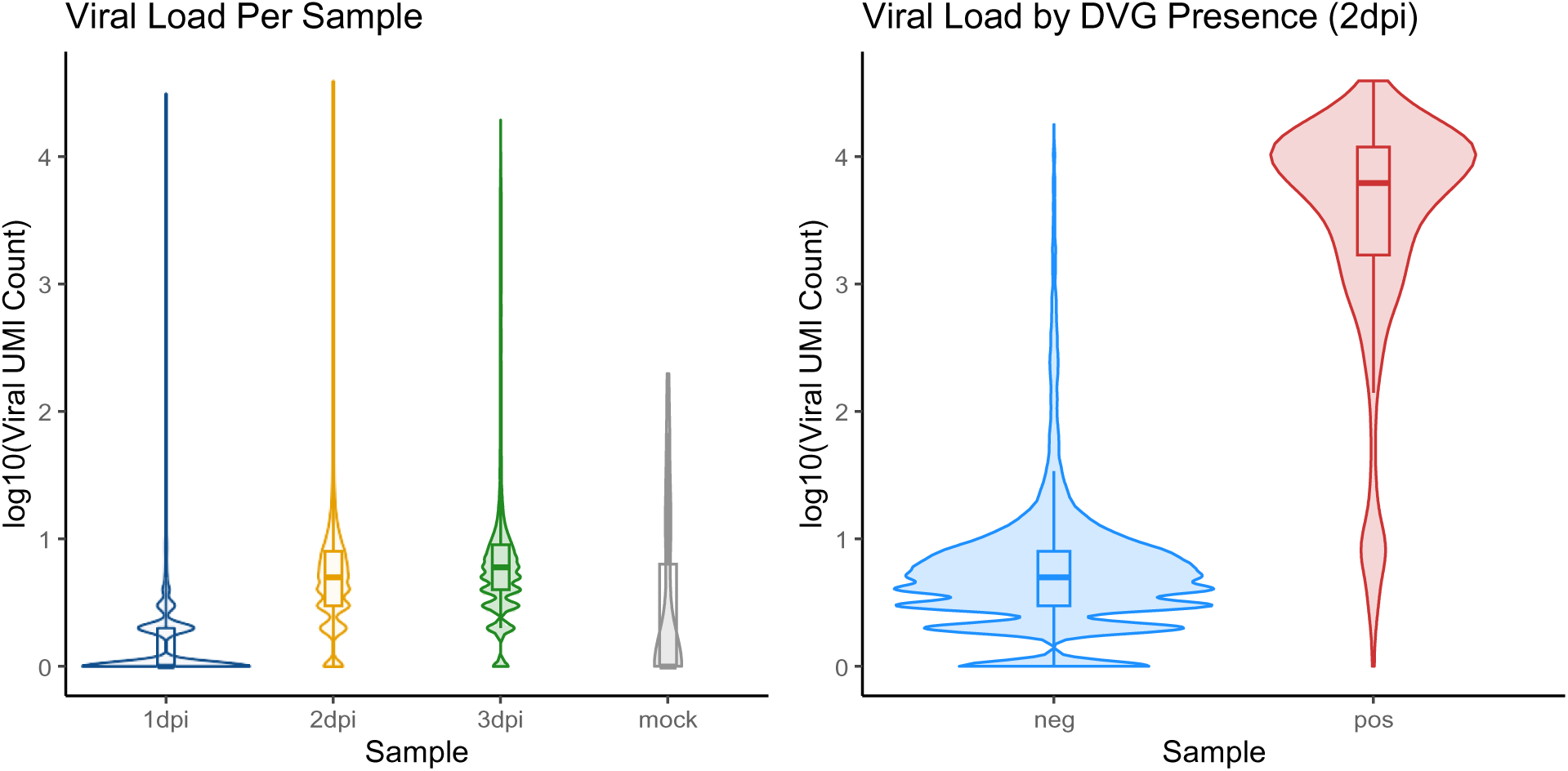

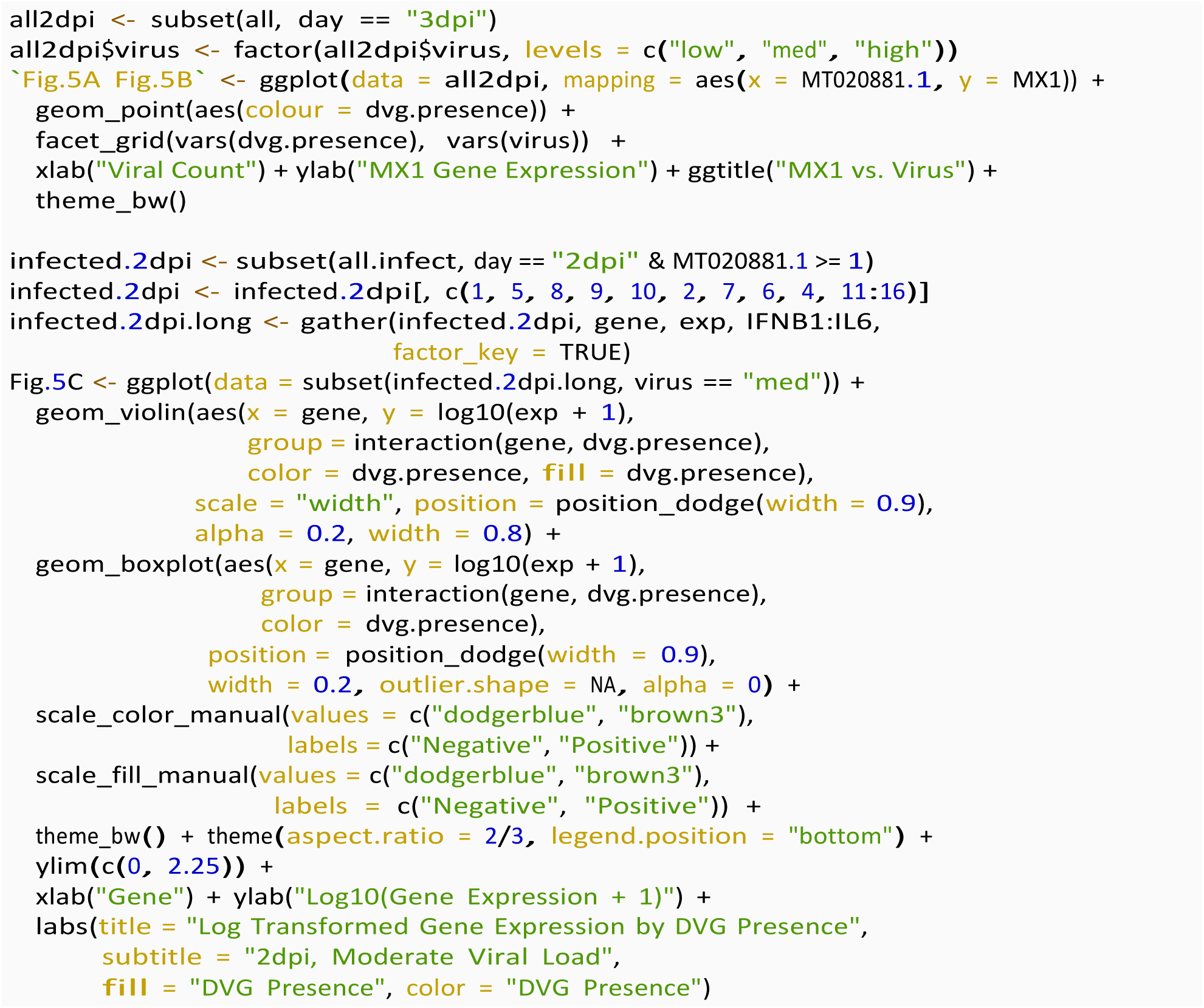
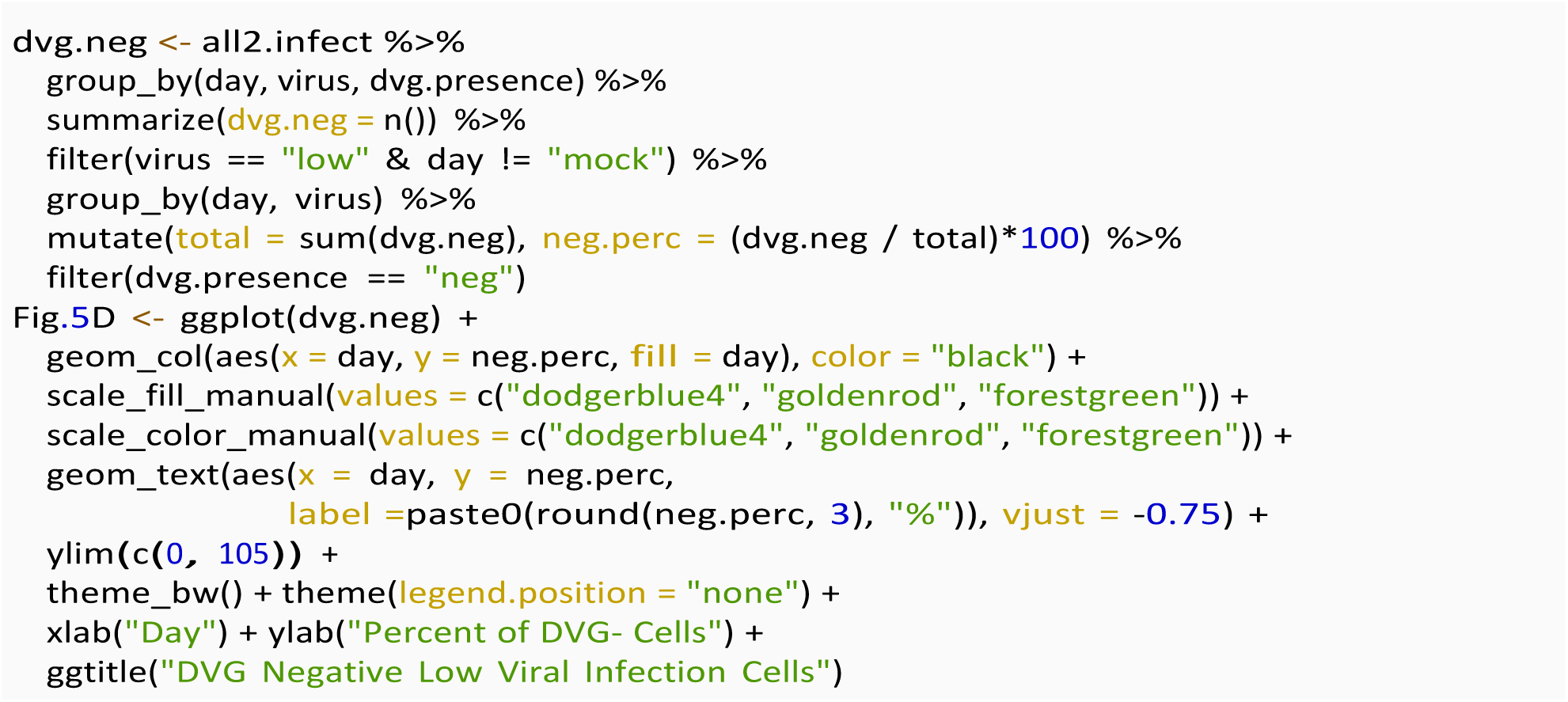

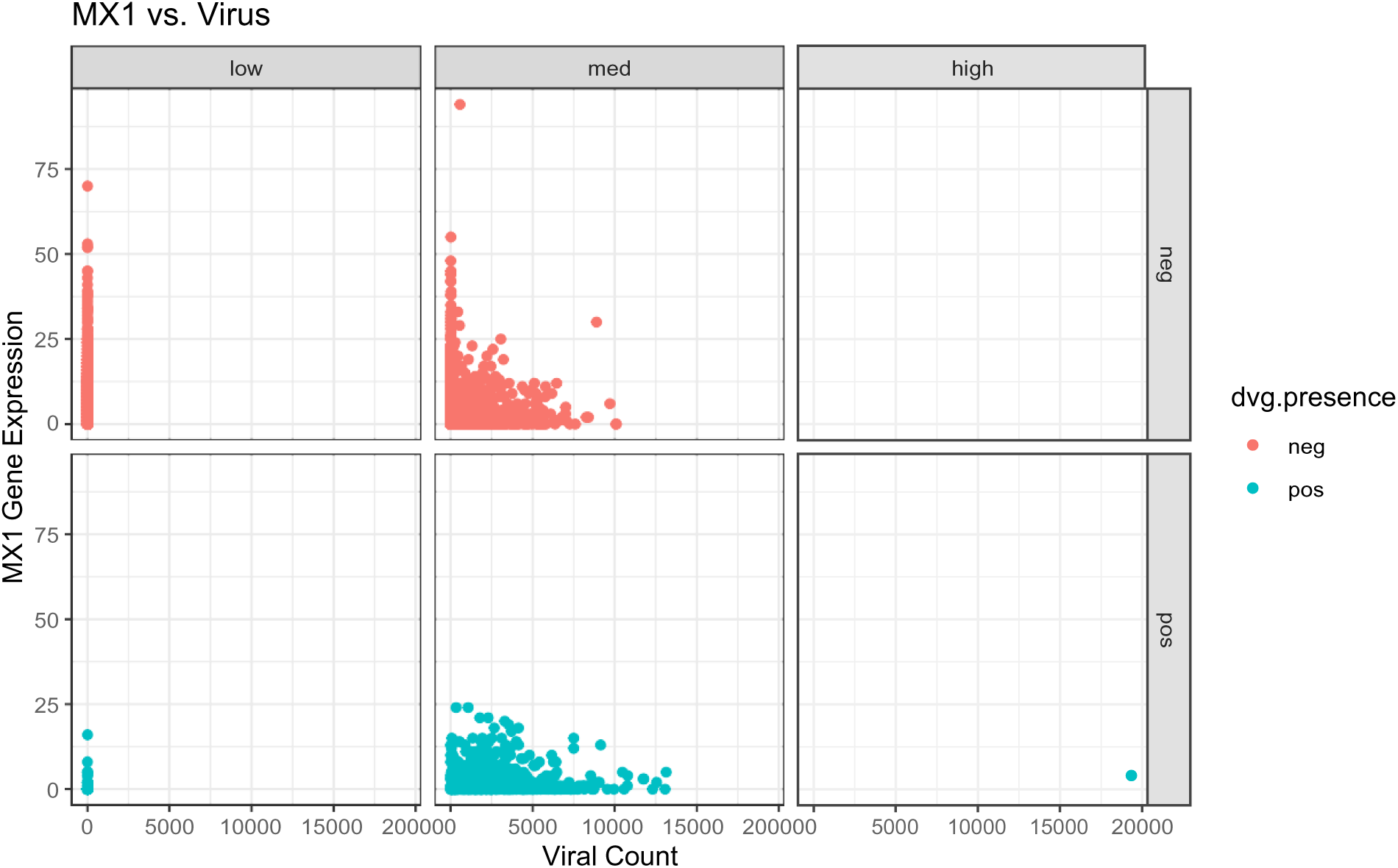

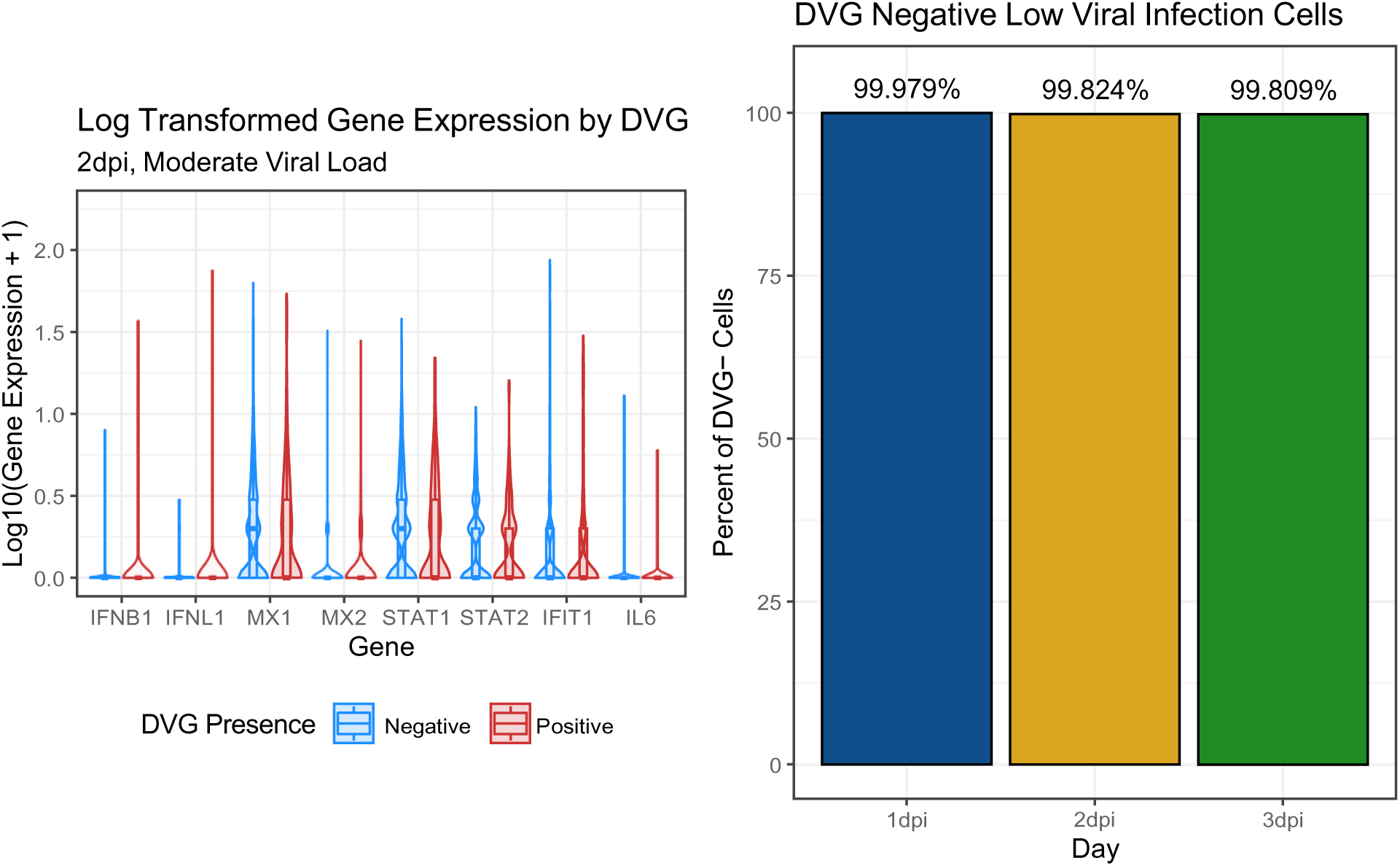

### Cellranger and Seurat

The purpose of this section of the standard operating procedure is to outline the pipeline used for Cellranger and the Seurat R package for DVG counting and their impact on host responses from single cell RNA-seq.

We used version 6.1.2 of Cellranger to generate the gene expression matrices for our single cell RNA-Seq analysis.

This part of the analysis used the following R packages:

- Seurat
- Matrix
- sctransform
- Mast
- DESeq2
- tidyverse
- ggplot2
- dplyr
- data.table

### Reference Genome

#### SARS-CoV2 FASTA and GTF

We dowloaded the SARS-CoV2 genome fasta file. For the MT020881.1 strain, it can be found in the following ncbi link. https://www.ncbi.nlm.nih.gov/nuccore/MT020881.1?report=fasta)

We made a custom GTF for the SARS-CoV2 genome such that it was labeled as a ‘gene’ in the human reference genome to which the covid genome was appended.

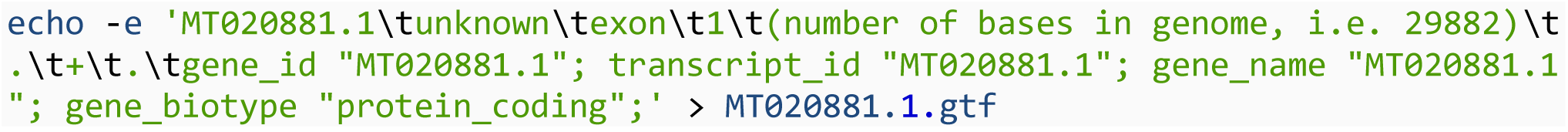

The resulting gtf file looked like the following with the ‘cat MT020881.1.gtf’ command.

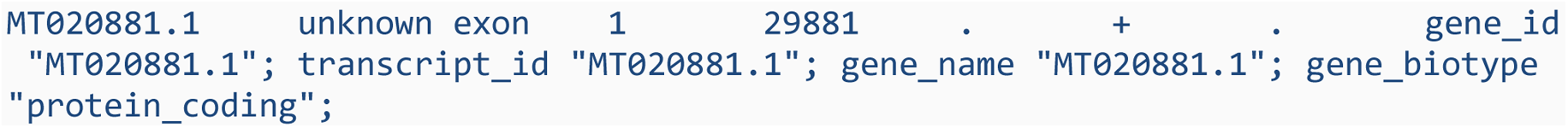

#### Creating reference package for Cellranger

We used the following shell script to run the mkref command in cellranger to create the reference package.

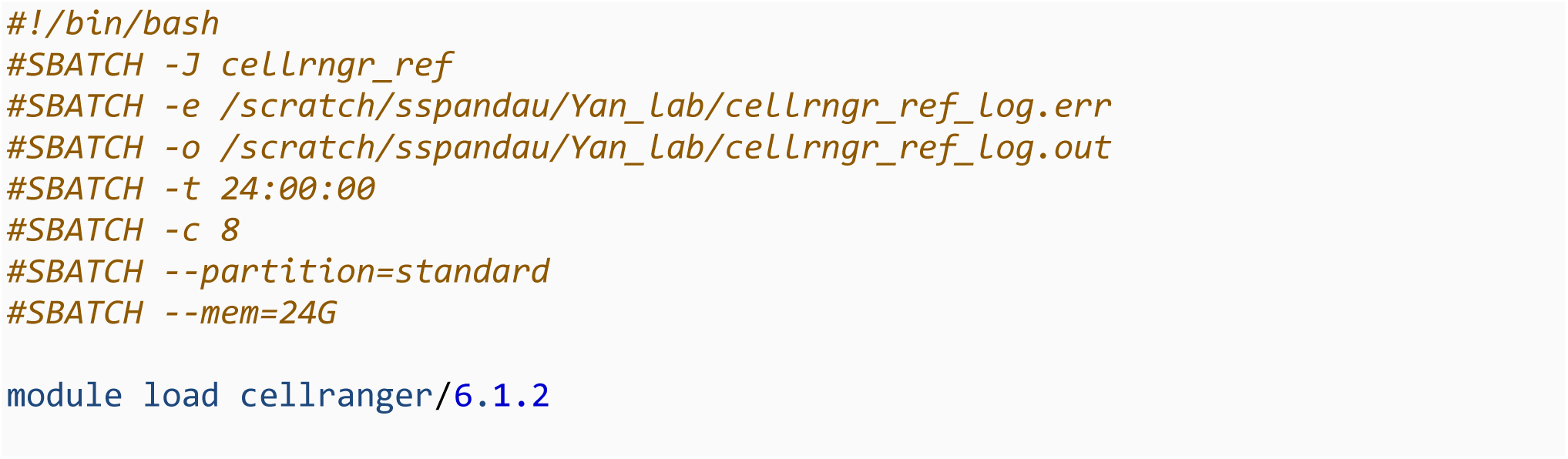
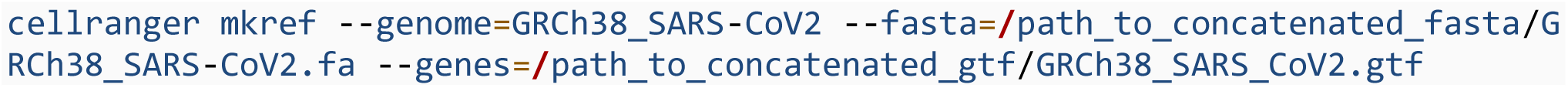

The ‘–genome=’ argument was for naming the resulting reference package. The ‘–fasta=’ was for inputing the reference fasta, and the ‘–genes=’ was for inputing the corresponding gtf file.

### Gene Expression Matrix

#### Cellranger count

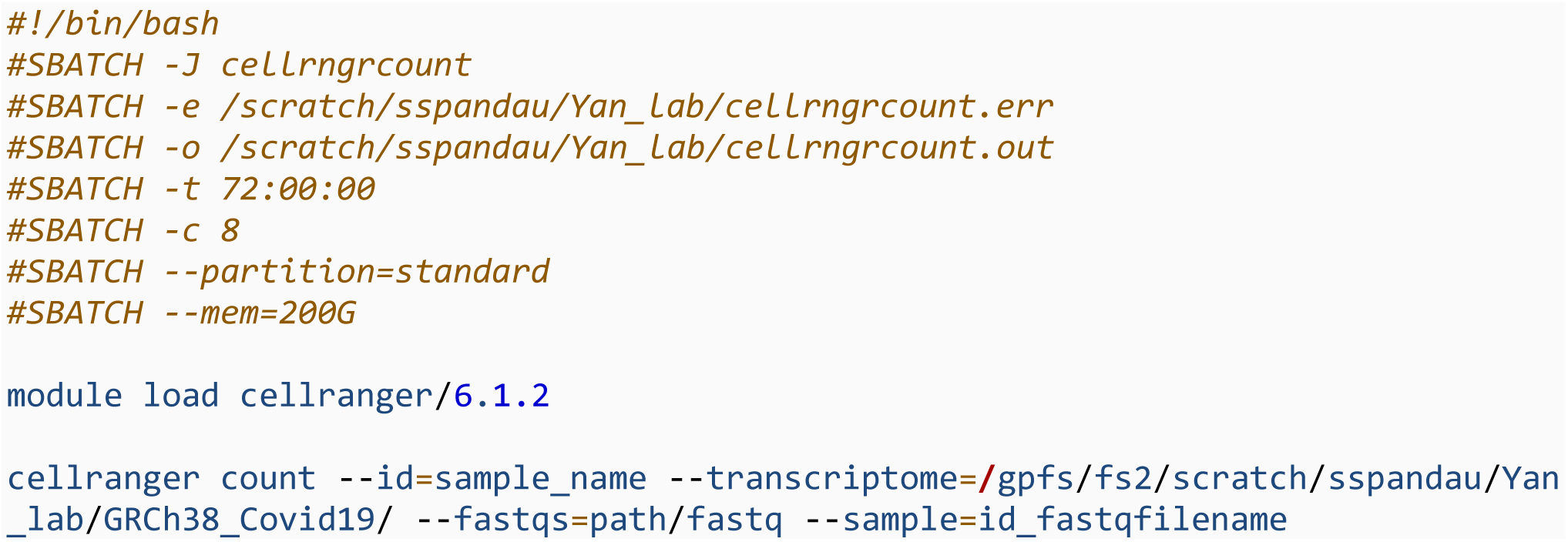

The ‘–id=’ argument was for naming the output folder which contains the gene expression matrix.

The ‘–transcriptome=’ argument was for inputing the path to the reference genome folder that was previously generated. The ‘–fastqs=’ argument was for input the path(s) to the R1 and R2 fastqs of the sample. The ‘–sample=’ argument was for input the sample id, which was the first few characters at the beginning of the R1 and R2 fastq file names.

#### Loading matrix into R and creating csv files

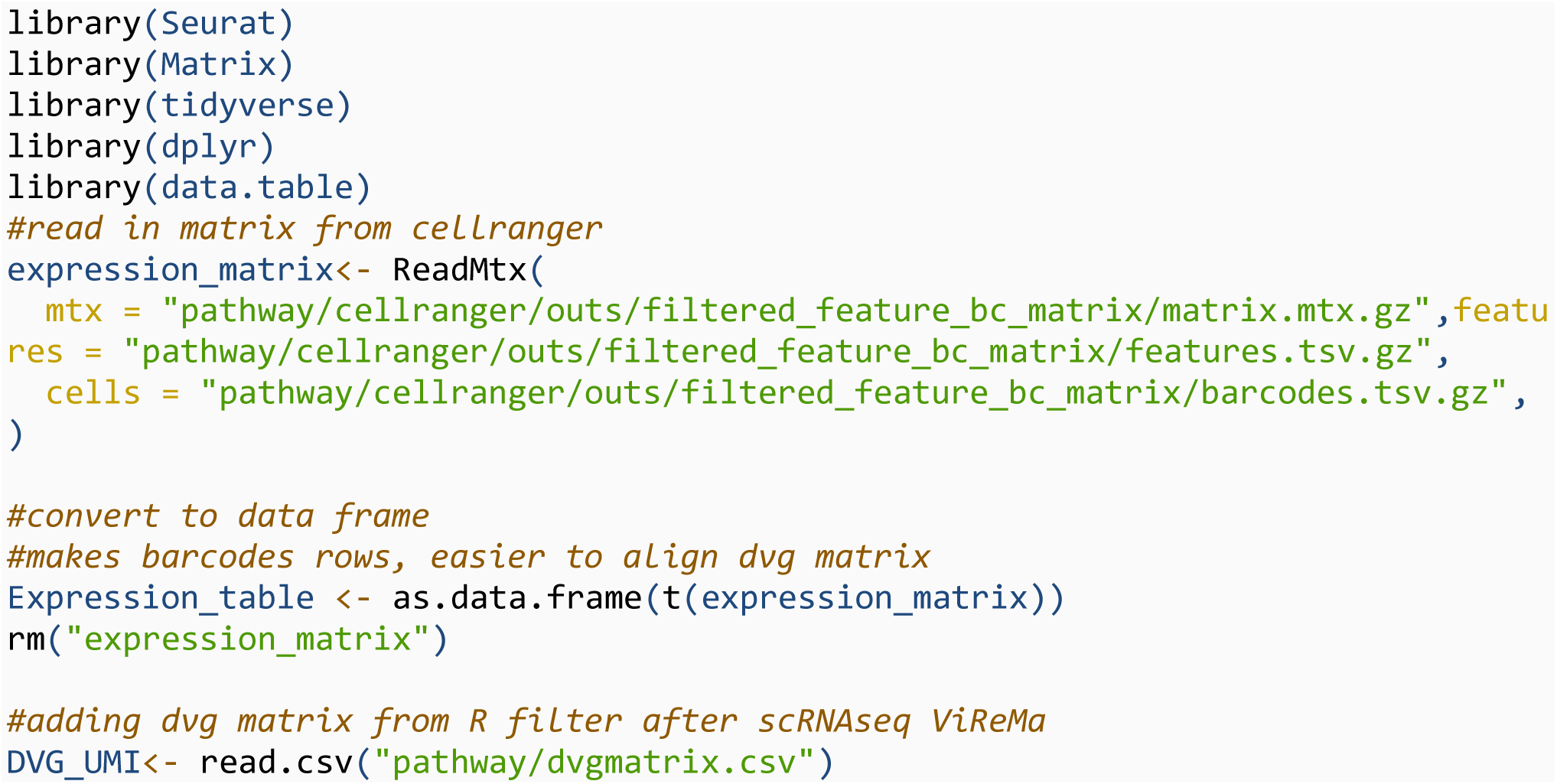
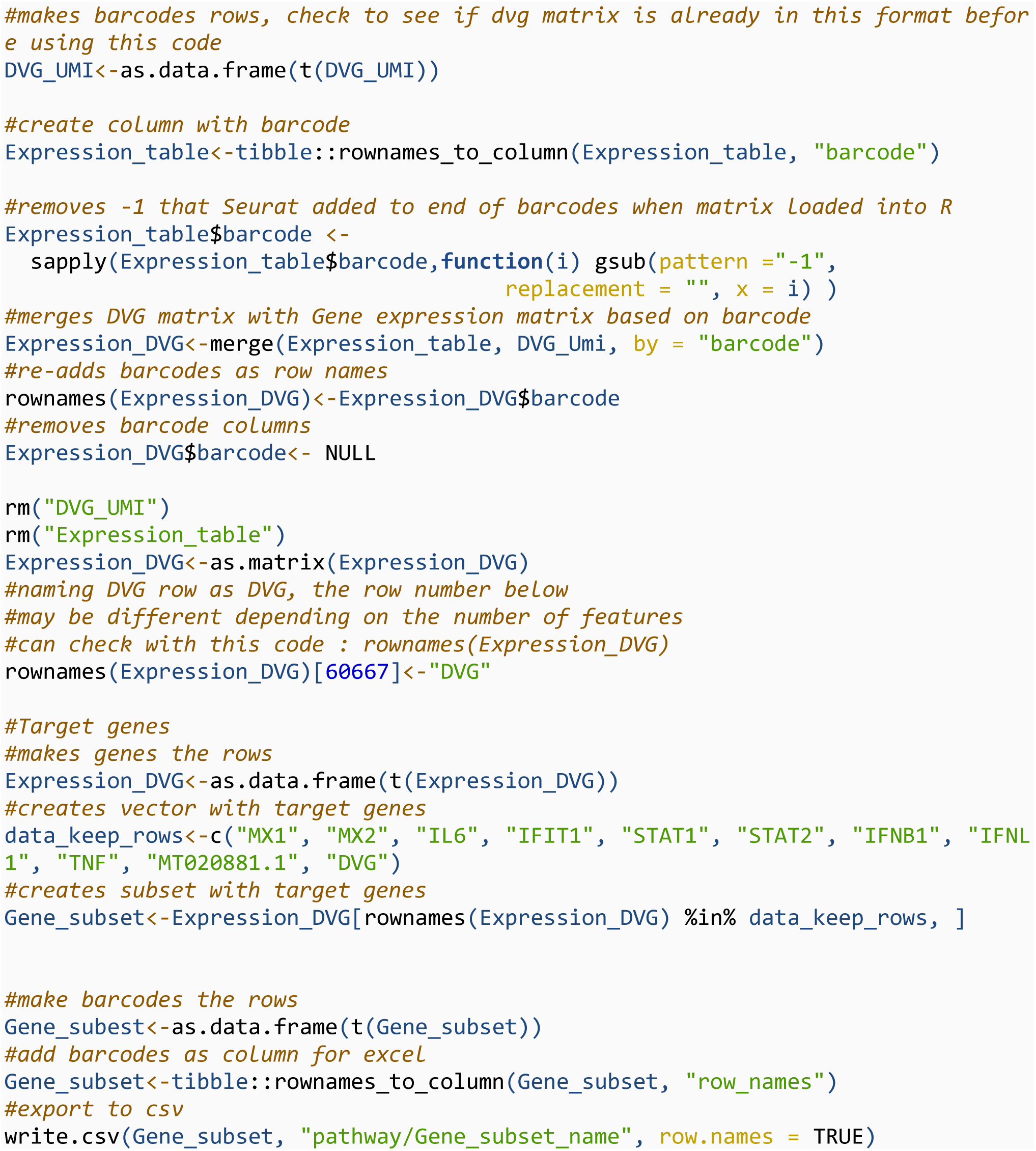

#### Celltype Identification

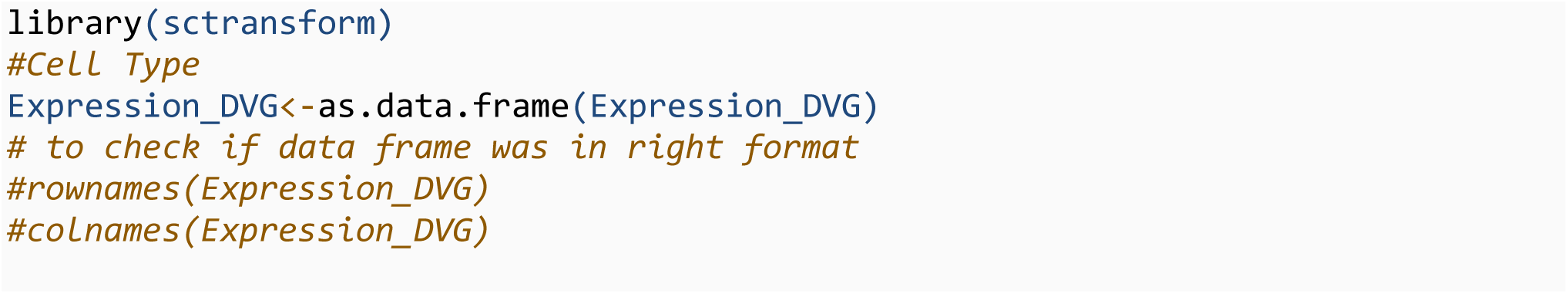
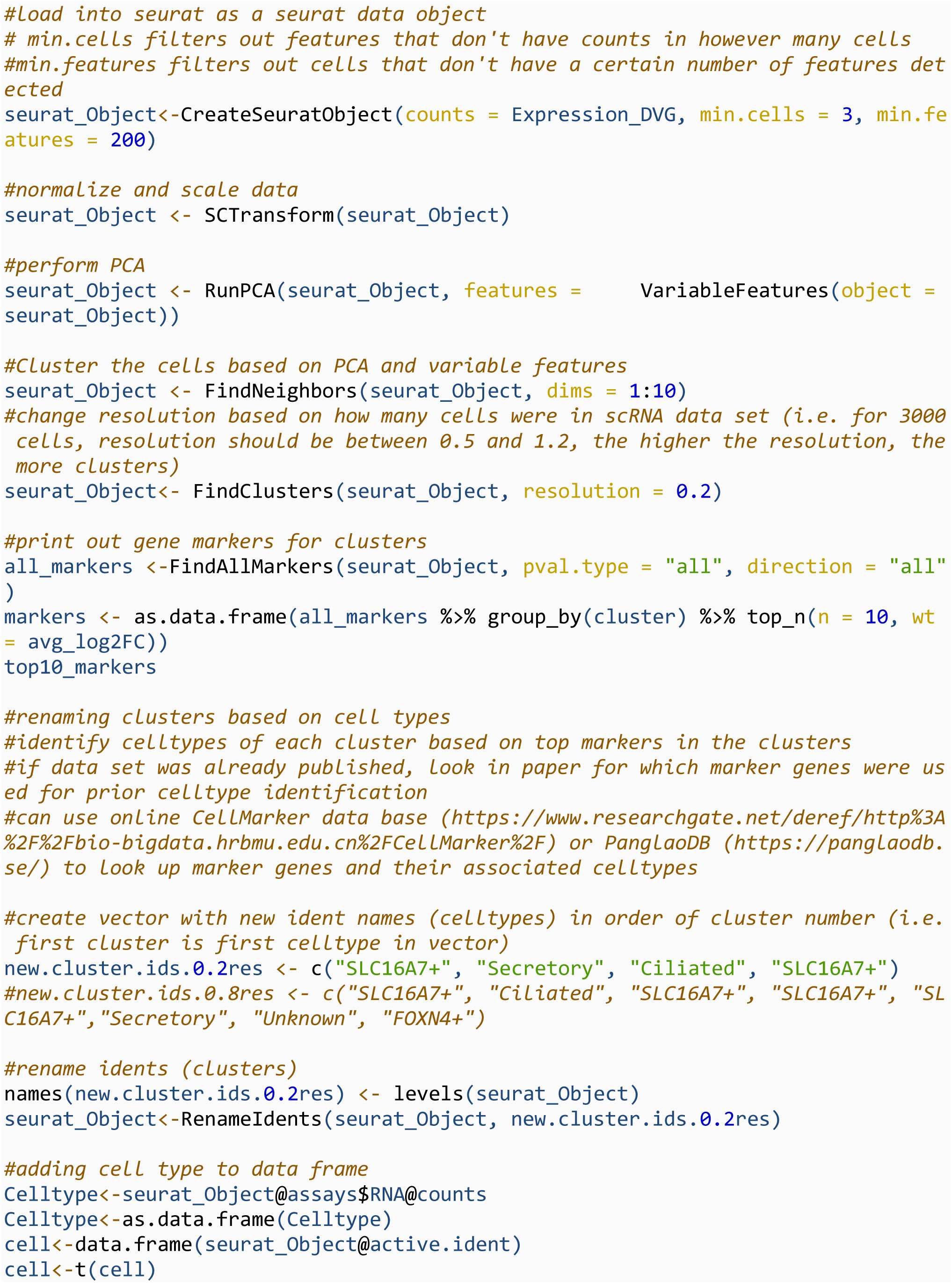
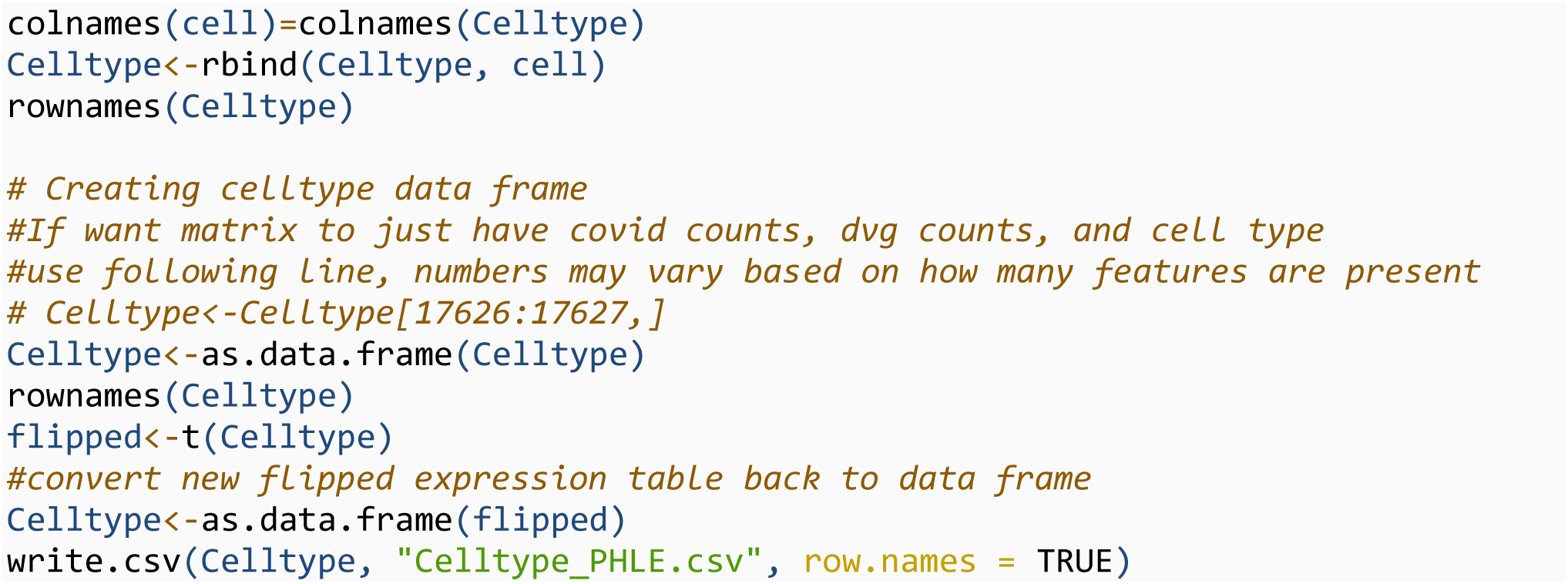

### Celltype Percents

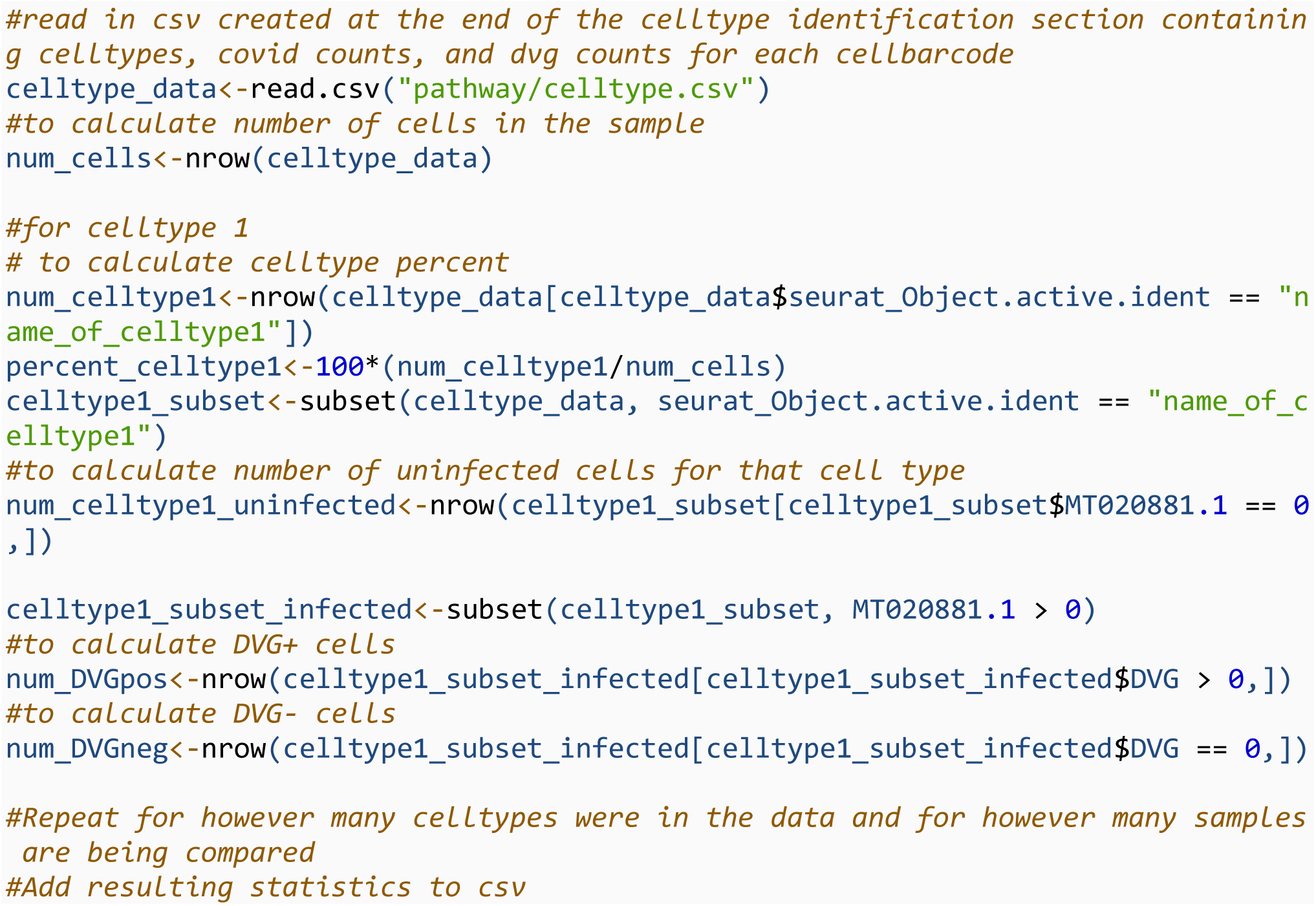

### Differential Gene Expression

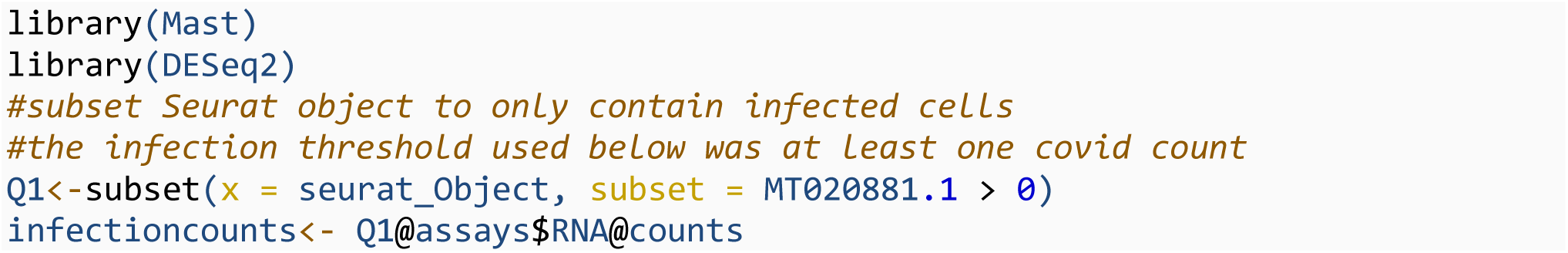
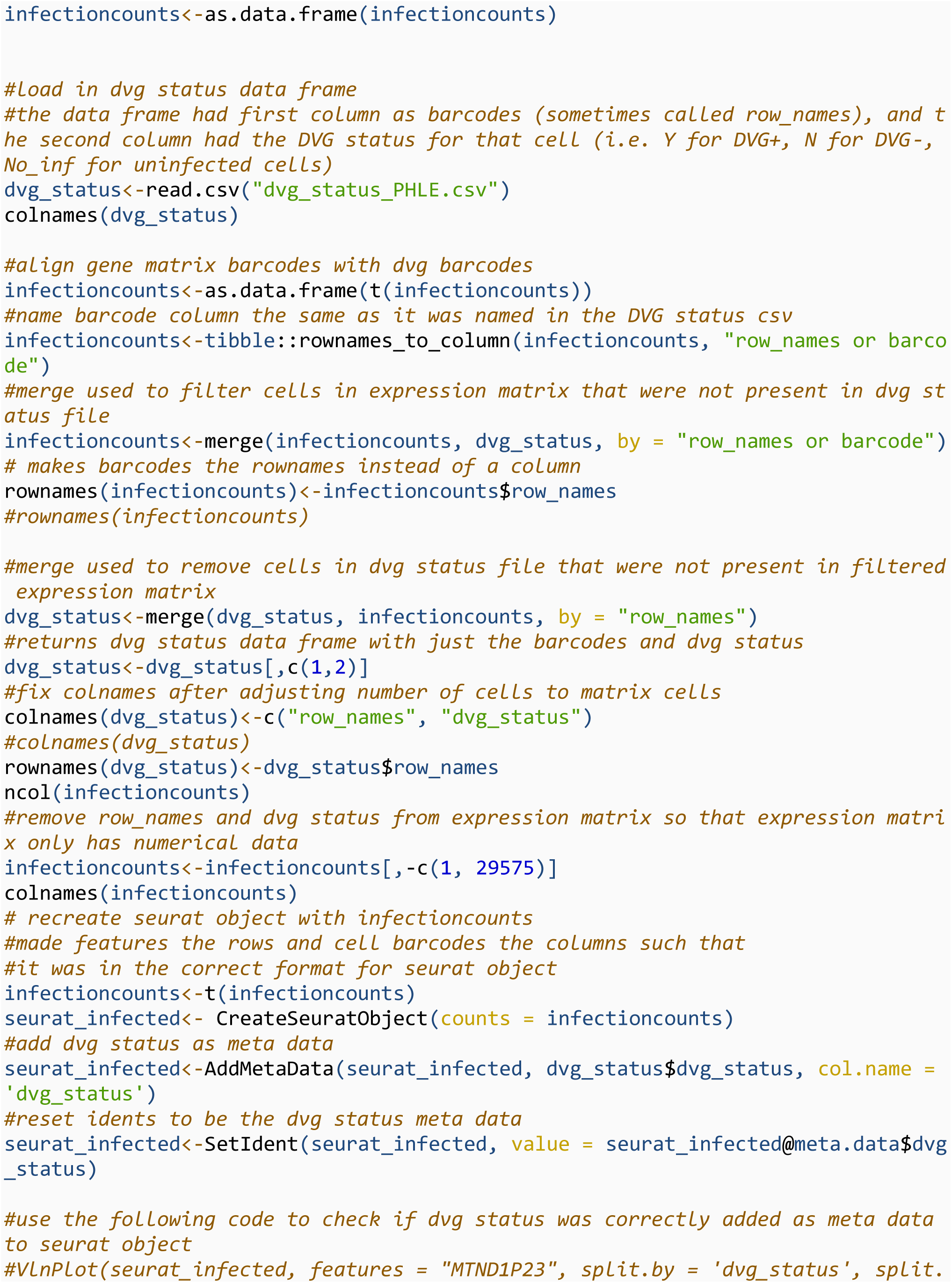
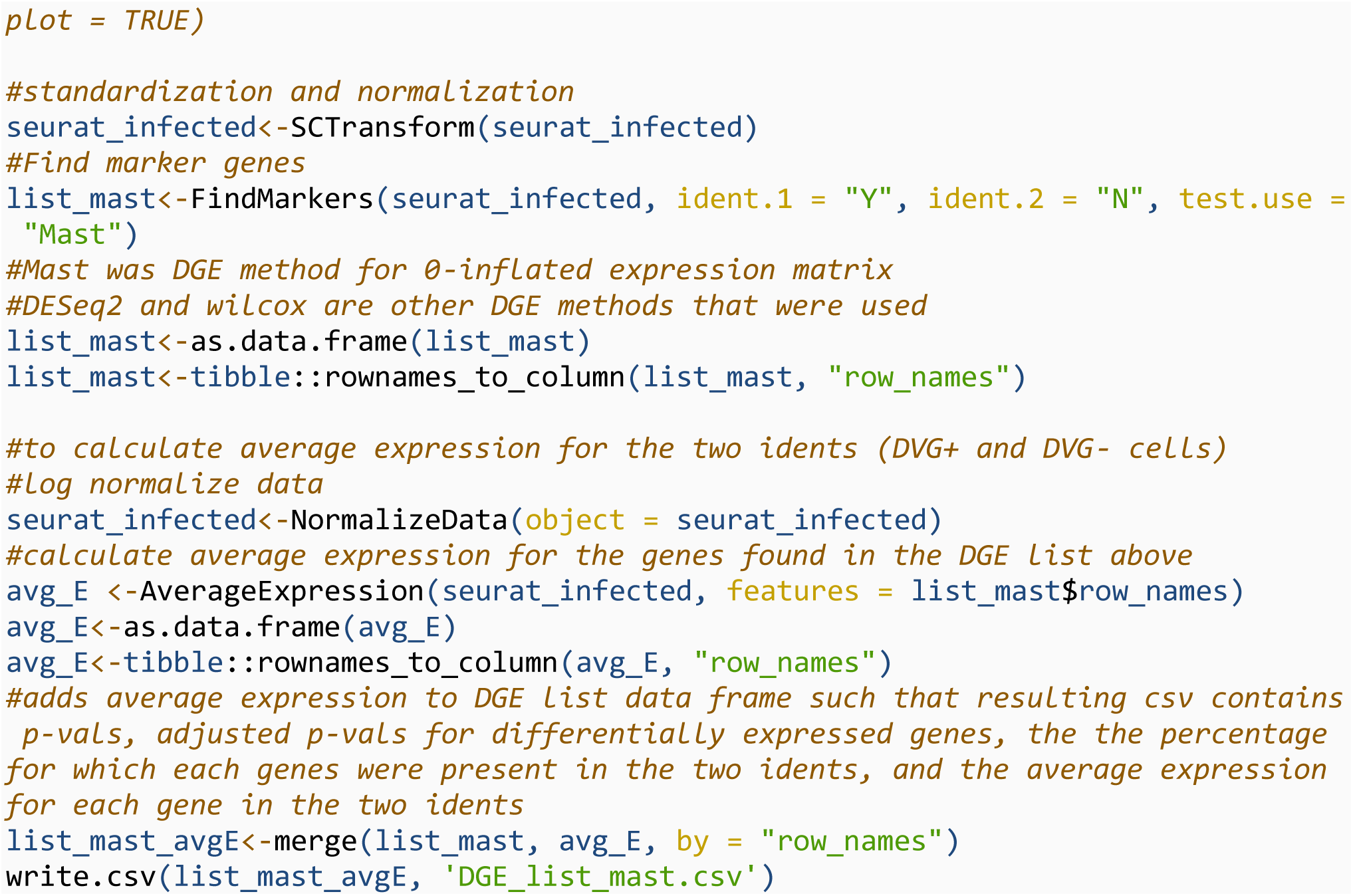

#### Fig 5 A: GO dotplots

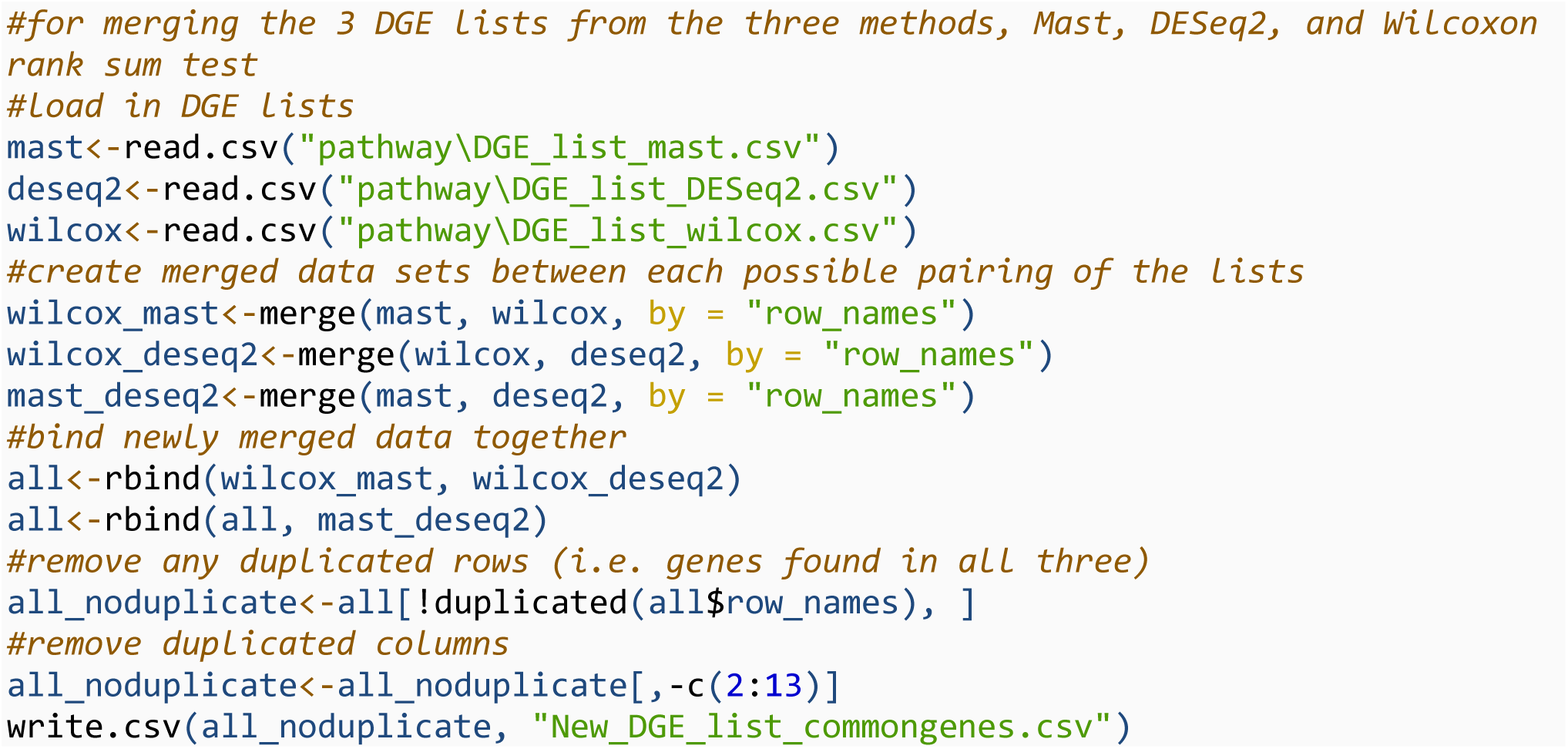

After submitting DGE list to DAVID functional annotation tool and selecting the top pathways found in each cluster of the DAVID results, we used the following script to generate the GO dotplots.

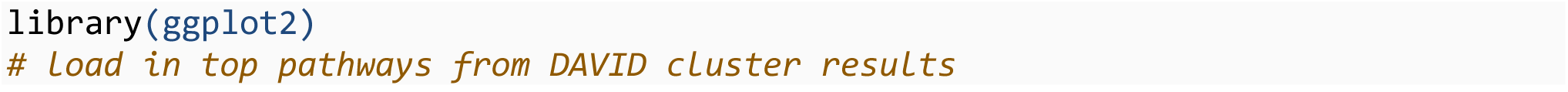
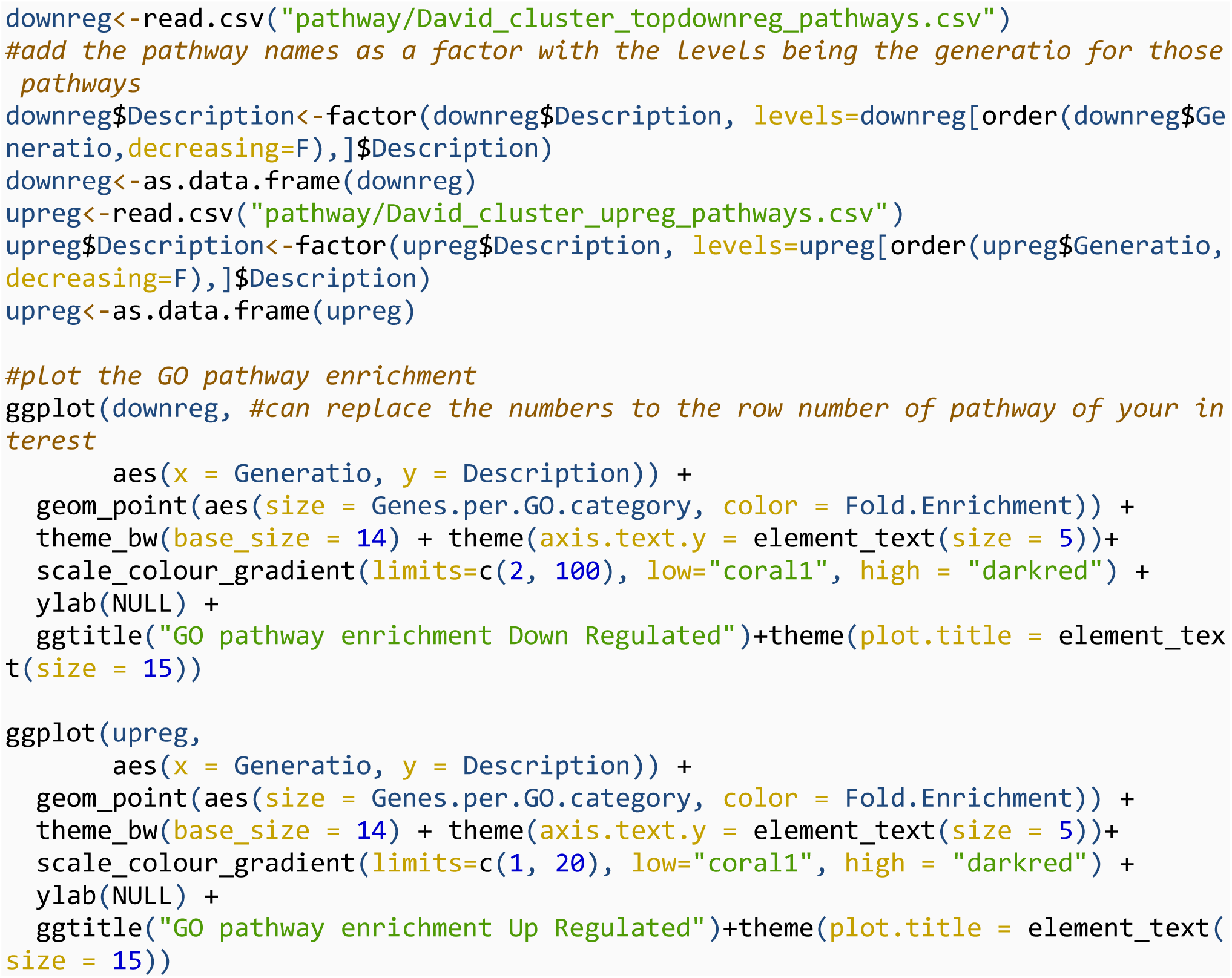

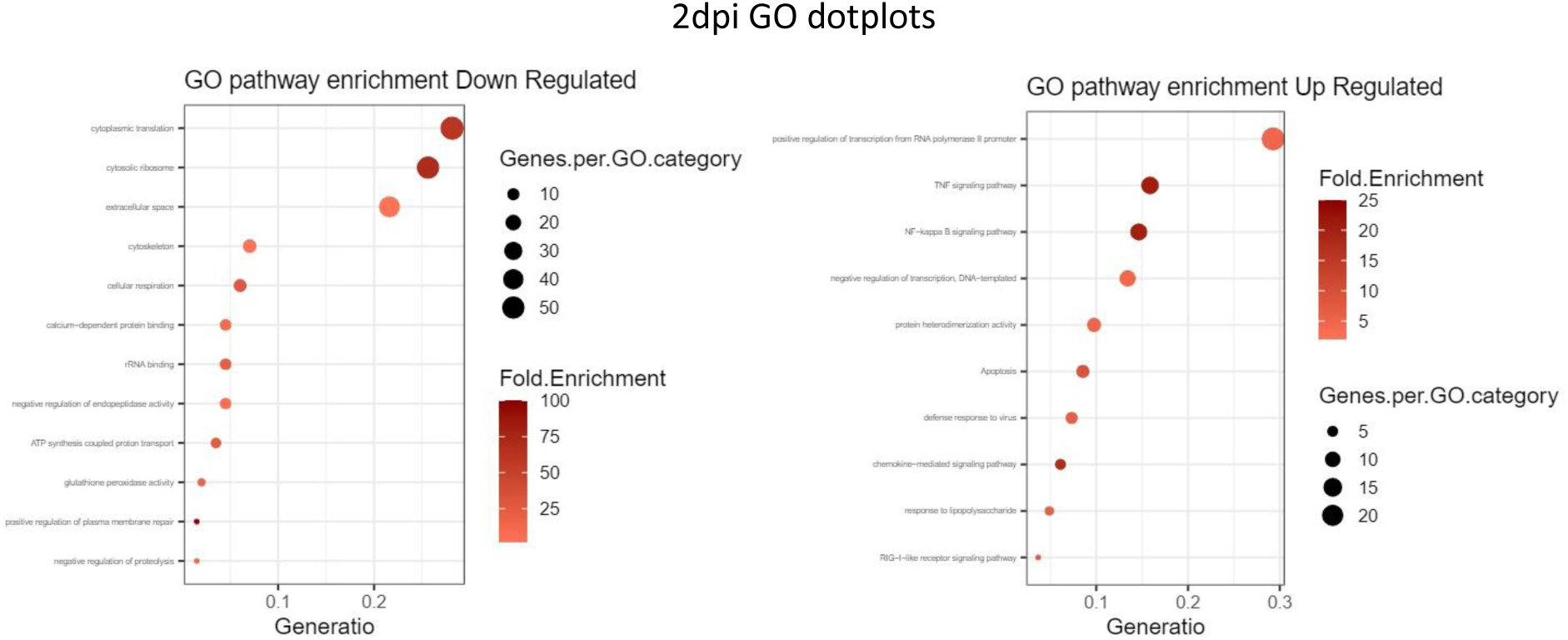

